# Parametric neural control differentiates top neural network models of primate visual cortex

**DOI:** 10.64898/2026.08.16.745063

**Authors:** Jacob S. Prince, Binxu Wang, Thomas Fel, Akshay V. Jagadeesh, Parisa A. Vaziri, George A. Alvarez, Margaret S. Livingstone, Talia Konkle

## Abstract

Leading deep neural network encoding models predict visual cortical responses with nearly indistinguishable accuracy, raising the strong inference that these models have converged on the same underlying brain-aligned parameterization of natural image space. Here we demonstrate that this is not the case. We introduce axis-aligned feature accentuation, which converts each model’s fitted encoding axis into graded stimulus perturbations that are predicted to parametrically control neural firing within and beyond the natural-image range. We generated over 27,500 controller stimuli from ten leading vision models and presented them to five macaques in closed-loop experiments targeting early, mid-, and high-level visual areas. Despite matched natural image predictivity, models diverged strongly in their ability to control neural firing using accentuated stimuli, revealing that most model encoding axes failed to capture the precise tuning of their corresponding neurons. The two adversarially trained models showed a consistent advantage, though adversarial robustness was only weakly predictive of neural control across other models. Instead, control was better predicted by the spatial frequency structure of the input gradient: the distribution of pixels influencing each encoding axis. Overall, these results establish neural control via axis-aligned feature accentuation as a causal method to assess the alignment between how neurons and models parameterize the visual world.

## Introduction

Deep neural network (DNN) encoding models have emerged as powerful tools for predicting visually evoked brain responses across species and measurement modalities, from single-unit activity in macaque V1, V4, and IT to fMRI, EEG, and MEG responses across the human visual hierarchy (1–5). These successes have motivated the view that modern vision models provide a computationally explicit framework for formulating and testing hypotheses about visual processing in the brain (6–10). However, neural encoding scores have increasingly saturated, with recent surveys showing that hundreds of distinct models achieve nearly identical scores despite substantial differences in architecture and training objectives (11, 5). These results raise a fundamental question: are these diverse models truly equal in their alignment (12), or do their similar prediction scores mask distinct representations and computations (13–18), some of which may be more aligned with biological vision than others?

To answer this question, recent work has focused on developing stricter criteria and underlying principles for measuring the alignment between models and neural data. For example, some have focused on updating the brain-model regression procedures commonly used to assess neural predictivity, either by including stringent forms of regularization that limit the flexibility of the mapping (19–22), by assessing bi-directional mappings between models and brains (23, 24), or by alternating data collection and model fitting to improve the scope of predictivity (25). Others have focused on using custom stimulus sets most likely to differentiate models in their degree of brain or behavioral alignment. Classic examples include “controversial” stimuli, where images are generated or selected to produce maximally divergent predictions across candidate models (26). Related approaches use targeted image sets to test whether models reproduce known behavioral signatures in humans and other primates, including error patterns (27), invariances (28), adversarial susceptibility (29), and shape vs. texture bias (30, 31, 15, 32). Despite these efforts, there remains limited consensus on how best to define and assess brain-model alignment.

As a possible source of new traction on this problem, we draw on studies of neural control (33–37). These efforts use model-synthesized stimuli to modulate the response of a neuron or neural population, often in an online closed-loop paradigm. One approach relies on evolutionary algorithms and generative image priors to synthesize stimuli that strongly drive visual neurons (38–41), without an explicit model of neural tuning. Alternative approaches to neural control first map DNN features to neural responses using a set of images for model fitting, then use the resulting “encoding axes” to synthesize new images that drive, suppress, or selectively modulate individual neurons or neural populations (34, 35, 42–45). These encoding-based control paradigms carry stronger inferential power than standard encoding benchmarks, because they intervene on the input according to a model’s own predictions, shifting model evaluation from the observational to the interventional level of causal inference (46, 47). To date, neural control efforts have largely been motivated by either specific applied goals (e.g., precision modulation to deliver targeted neural or behavioral therapies; 42), or by the scientific goal of using successful control to better understand neural tuning (38, 39, 33, 34, 48). However, the controller model is often a means to an end: success is defined by the quality of control, with relatively little emphasis on whether one model provides a more biologically aligned account than another.

Here, we treat model-based neural control as a critical assay of model-brain alignment. We start from the standard view that a neural encoding axis can be defined as a weighted combination of features in a DNN layer, reflecting an explicit hypothesis about the stimulus attributes to which a neural site is tuned. Critically, because the axis is defined over the features of a specific layer, it inherits a dependence on the model-specific cascade of computations that transforms input pixels through intermediate stages to that layer. To probe the full computational commitments of each encoding axis, we use a gradient-based stimulus synthesis technique called feature accentuation (49). Starting from any seed image, we can accentuate it along the encoding axis in fine increments, producing a parametric stimulus sweep. Competing models with different internal computations may therefore produce different accentuations. If an encoding model has captured the site’s true tuning, neural responses should vary systematically across the sweep, tracking the model’s predictions, and allowing us to put each model-defined encoding axis to the test in this parametric control paradigm.

Here we compared how well a set of 10 deep neural network models can parametrically control neural responses over 25 sites spanning V3, V4, central IT, anterior IT, and STS regions of the macaque visual system. These models differed in their architectures, objectives, input diets, and learning protocols, but all achieve high and nearly indistinguishable brain-predictivity scores under current standard natural-image evaluation procedures (5, 11). Despite this matched predictivity, we found that models differed dramatically in their ability to parametrically control neural responses along their encoding axes. The two adversarially trained models showed a consistent advantage, but the degree of adversarial sensitivity did not explain variation in control among the remaining models. Instead, control was better predicted by the spatial-frequency structure of the encoding axes’ input gradients, which can be estimated prospectively before any control experiment. Broadly, this work establishes a framework for converting an encoding model into a causal hypothesis: a specific and testable trajectory of stimuli that should modulate the neural response. In doing so, we expose previously hidden differences in brain alignment among the most predictive encoding models, demonstrating that precision comparisons of model-brain alignment require targeted neural control experiments that go beyond natural image benchmarks.

## Results

### A closed-loop framework for comparing DNN encoding models via parametric neural control

To test whether models that appear equivalent on natural images make different hypotheses about neural tuning, we designed a four-phase closed-loop model comparison experiment (Figure 1). This paradigm consisted of: (i) a calibration phase, in which we measured neural responses to a broad image set; (ii) offline fitting of DNN-to-neuron encoding models; (iii) synthesis of axis-aligned accentuated stimuli; and (iv) a control phase, in which the synthesized stimuli were presented back to the animals during neural recording.

**Figure 1.**
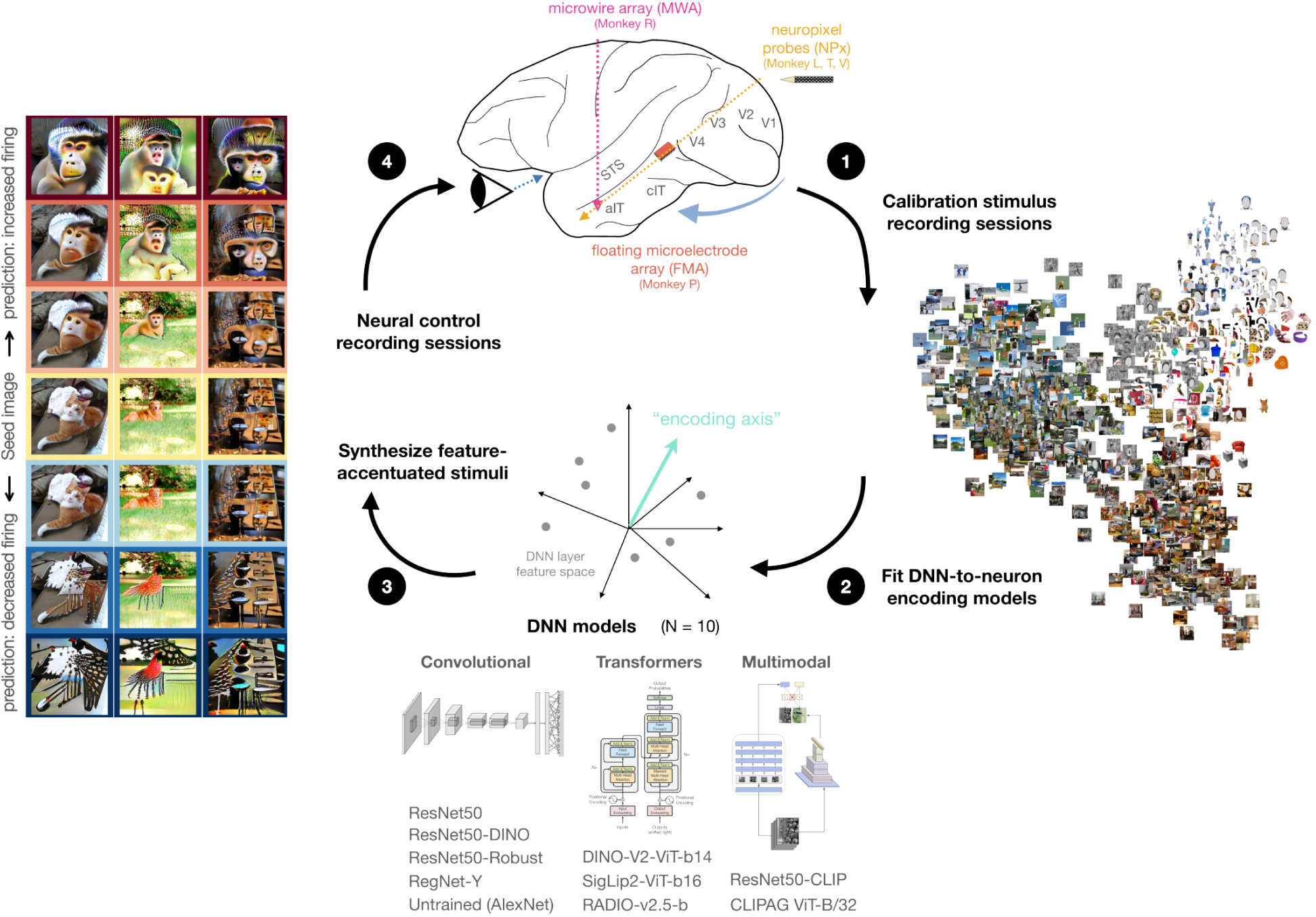
Closed-loop framework for evaluating DNN encoding models through parametric neural control. The experimental paradigm proceeded in four phases. (1) Neural responses to a diverse calibration image set were recorded from five macaques using Neuropixels probes in areas V2, V3, and V4, and STS, a chronic floating microelectrode array in central inferotemporal cortex (cIT), and a microwire array in anterior inferotemporal cortex (aIT). n = 969 calibration images. (2) Encoding models were fit separately for each DNN and each targeted neural site. The encoding axis was defined as the ridge-regression weight vector that best predicted a site’s responses from DNN feature space, with cross-validated layer selection. (3) Axis-aligned accentuated stimuli were synthesized by perturbing natural seed images along each encoding axis at graded increments, yielding image sweeps predicted to modulate firing from below to above the site’s calibration-image response range. (4) The synthesized stimuli were presented back to the animals during neural recording to assess each model’s capacity for parametric neural control. Ten DNN models were compared, spanning convolutional and transformer architectures, trained with supervised, self-supervised, contrastive, language-aligned, and adversarial training objectives.

We recorded neural responses from five macaques across visual cortical areas V3, V4, STS, central inferotemporal cortex (cIT), and anterior IT (aIT), using both chronic floating microelectrode arrays and Neuropixels probes (see Methods; Fig. S3). We selected five sites per animal with diverse, reliable tuning profiles as the targets for neural control, spanning a range of category preferences (Fig. S2, S3, S4; see Methods; mean calibration phase noise ceiling *r* = 0.88).

We selected ten deep neural networks for comparison, with a goal to span convolutional and transformer architectures, with supervised, self-supervised, contrastive, language-aligned, and adversarial training (50) objectives, and one untrained model that serves as a baseline reference (see Methods, Table 2). For each target site, we characterized its tuning with a hypothesized “encoding axis”, estimated separately for each DNN model using ridge regression with cross-validated layer selection (see Fig. S5).

In the third phase of the study, we used these encoding models to synthesize the feature-accentuated controller stimuli for the closed-loop experiments. Each “accentuated image sweep” targeted 11 predicted neural response levels, ranging from 25% below to 50% above the site’s native response range to natural images.

Finally, we collected neural responses to these accentuated controller stimuli, intermixed with the (initial) natural images for replication. This closed-loop design allowed us to evaluate each model’s fitted encoding axis not only in its predictive accuracy on held-out natural images, but also in its capacity to parametrically modulate neural responses.

### Feature accentuation along DNN encoding axes

To synthesize controller image stimuli, we adapted the feature accentuation procedure developed by Hamblin et al. (49) and Fel et al. (51; MACO; see Methods for full detail). Each image synthesis run takes as input a seed image, a neural encoding axis (expressed as a direction in the feature space of a DNN layer), and a desired target response. Using gradient-based updates, the algorithm returns an accentuated image: a modified version of the seed image that highlights or de-emphasizes evidence for a given feature direction in order to achieve the targeted level of activation. This allows us to generate sweeps of images that are predicted to progressively and parametrically activate a given neural site along the fitted encoding axis (Figure 2A–C).

**Figure 2.**
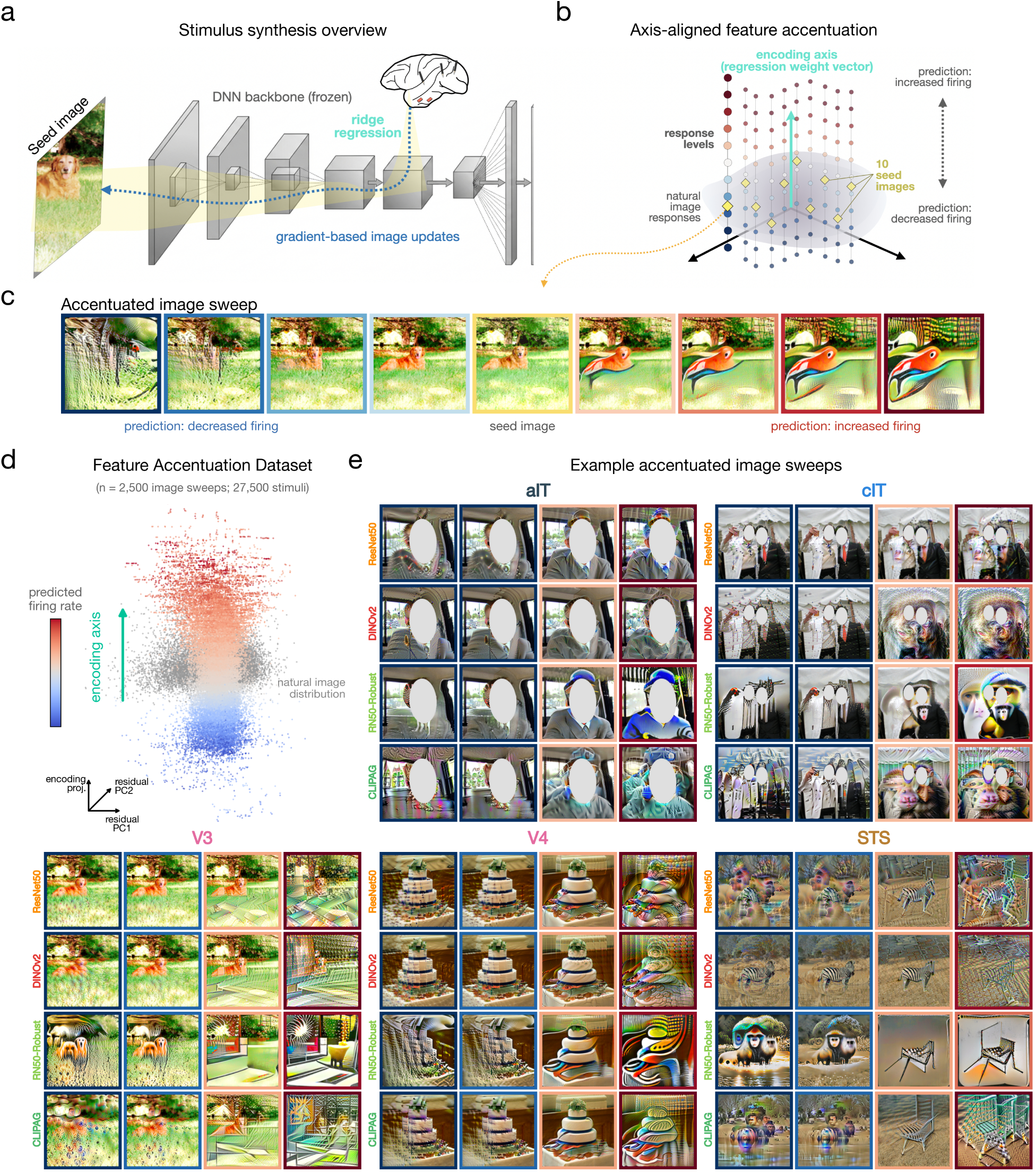
Feature accentuation along DNN encoding axes. (A) For each neural site and DNN model, a ridge regression weight vector (an “encoding axis”) was fit from layer activation PCs to recorded responses using the calibration stimuli. Gradient-based image updates were then propagated through the model to modify a seed image toward a target predicted response level along the encoding axis. (B) Schematic of axis-aligned feature accentuation in DNN layer feature space. The gray cloud indicates the distribution of natural image responses, and dots are colored by predicted firing rate. The cyan arrow indicates the encoding axis, and the two other axes reflect two orthogonal latent dimensions in the source layer feature space. Yellow diamonds indicate the embeddings of 10 seed images, each accentuated to 11 target response levels spanning below and above the natural-image response range. (C) Subset of an example accentuated image sweep for one seed image, model, and neural site. The original seed image is shown at center, with progressively decreased predicted firing to the left and increased predicted firing to the right. Border colors indicate predicted response levels (blue = low, red = high). (D) Full feature accentuation dataset visualized in the three-dimensional principal component space of an example encoder (ResNet50). n = 2500 image sweeps; 27,500 total stimuli. The vertical dimension reflects projection onto the encoding axis, and the two horizontal dimensions reflect the first two principal components of residual variance. The gray region indicates the natural-image response range from the calibration set. (E) Example accentuated image sweeps for one representative site per cortical area, shown for four DNN models. Rows correspond to models accentuating the same seed image for the same neural site, and columns show four predicted response levels from low to high firing rate.

Generating these sweeps is inherently ill-posed: for a given level of predicted activity, there are potentially infinite images that achieve it. For many vision models, unconstrained gradient-based optimization in pixel space typically resolves this underdetermination in one of two degenerate ways: either by covering the image in hallucinated high-frequency artifacts, or by settling on an imperceptible adversarial perturbation that shifts the prediction without altering anything a viewer would recognize as content (49, 51; but see 52). We therefore applied a multifaceted regularization scheme to stabilize the image updates while preserving movement along the encoding axis. Image updates were performed in the Fourier domain with explicit penalization of high spatial frequencies, and several forms of image augmentation (e.g., random cropping, additive noise) further stabilized the synthesis and prevented artifacts (see Methods for full details). To preserve fairness across models and avoid hand-tuning, we also developed an automated hyperparameter selection procedure to set regularization levels independently for each model-site combination (see Methods, Fig. S6).

This procedure reliably generated image sweeps that reached their full set of target activations (94.5% of stimuli; Figs. S7, S8) while remaining broadly axis-aligned (Fig. S9, S10). Qualitatively, the different models produced a diverse array of accentuations even for the same target site and seed image, with some emphasizing holistic structure and others enhancing textures or high-frequency content (Figure 2, Figs. S11–S16). These stimuli reflect genuinely divergent hypotheses about the underlying neural tuning.

### Strong natural image encoding does not imply accurate neural control

We first present the results from a single channel in anterior IT (aIT) and the ResNet50 deep neural network, which is one of the canonical visual encoders in our model set. How well does this model’s feature space predict this site’s responses to natural images, and critically, how well can it elicit parametrically controlled responses along its encoding axis?

As expected from numerous prior studies and benchmarks (11, 5), the best-layer feature space of ResNet50 supported a highly predictive encoding axis when fit on the diverse calibration set of natural images, achieving validation predictivity of *r* = 0.87 in the first recording phase (see Methods). This axis also remained stable in the second recording phase (*r* = 0.95 over the re-presented held-out images; Figure 3C).

**Figure 3.**
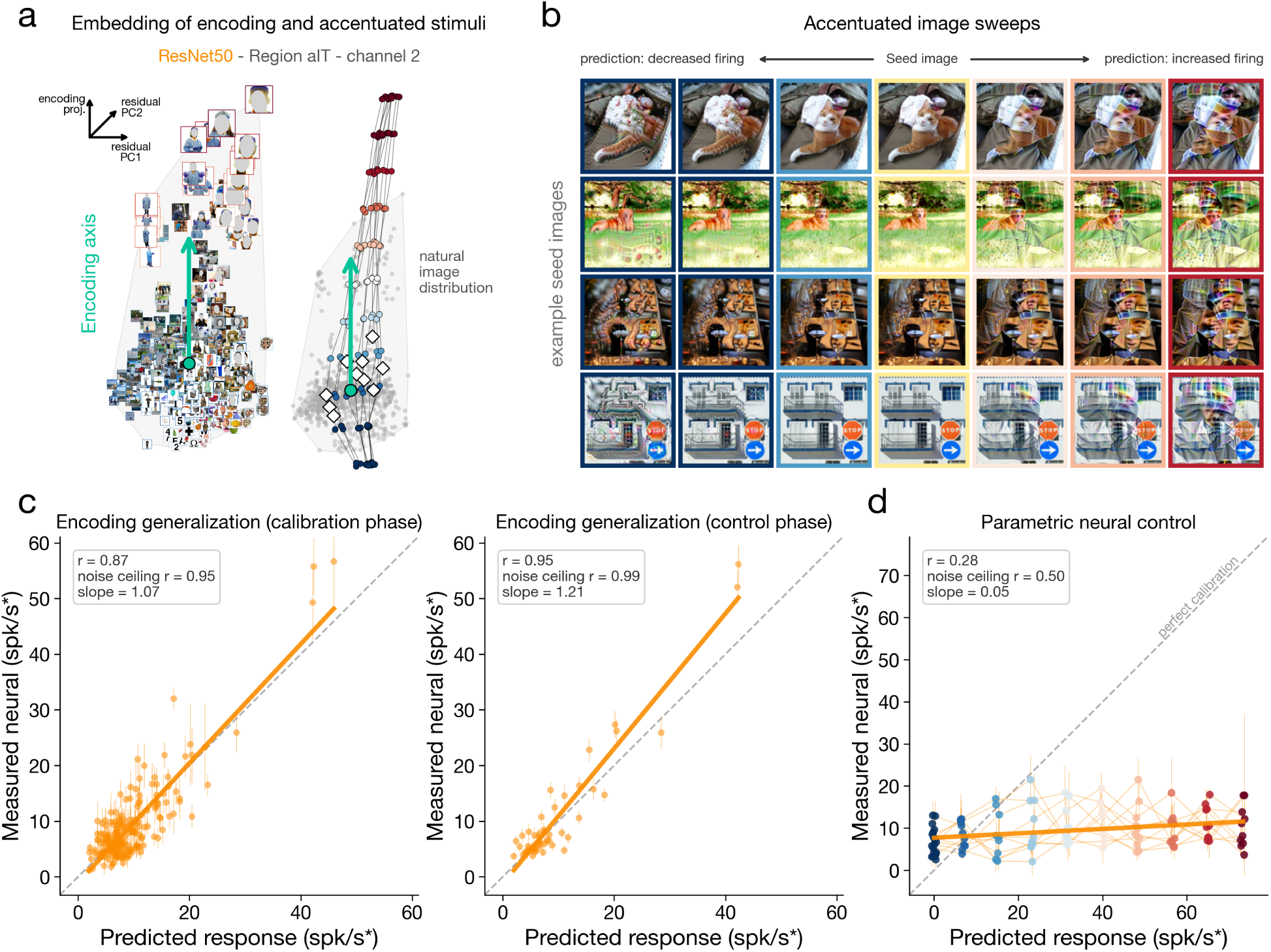
Strong natural image encoding does not imply accurate neural control. Results are shown for an example aIT site using the ResNet50 encoding model. (A) Visualization of site tuning in the best-fitting DNN layer feature space. Natural images are plotted in the same three-dimensional principal component space as in Figure 2D and colored by recorded firing rate. The strongest natural-image responses were elicited by images of human faces, particularly photographic portraits of lab members wearing personal protective equipment. The adjacent plot shows the 10 accentuated image sweeps projected into the same space, extending beyond the natural-image distribution along the encoding axis. (B) Example accentuated image sweeps for four seed images. Border colors indicate predicted response levels from low to high firing. (C) Encoding accuracy over natural images across phases. Scatter plots show predicted versus true neural firing rates for phase-1 validation images and phase-2 re-presented held-out images (the cross-session re-test set; see Methods). Dashed gray lines indicate identity, and orange lines indicate linear fits. (D) Parametric neural control over axis-aligned accentuated stimuli. Colored traces indicate seed image sweeps, and dots indicate individual accentuated images with vertical error bars indicating the SEM response over repeats.

However, we found that this model’s encoding axis was an exceedingly poor predictor of the site’s activity when tested over its axis-aligned accentuation images (*r* = 0.28; Figure 3D). While the stimuli were predicted to exert parametric control from 25% below to 50% above the natural image range, the actual neural responses showed these image sweeps led to effectively flat modulation (linear slope fit to neural responses = 0.05, where 1 would imply perfectly calibrated neural modulation; Figure 3D; see Methods). Notably, the neural responses to these accentuated stimuli were reliable across repeated presentations during the control sessions (110 stimuli, mean 4.0 repeats, Pearson *r* noise ceiling = 0.50; Fig. S31), arguing against measurement noise or session instability as the explanation for the control failure. Thus, high brain predictivity over a large and diverse set of natural images does not guarantee that the model’s encoding axis has a precise brain alignment under a more stringent test of parametric control.

Can any model successfully control neural responses along its encoding axis, and if so, to what degree can this capacity differentiate models in their brain alignment? The results from the 10 different models over 25 different sites are summarized in Figure 4. Considering the natural-image predictivity scores, as expected, nearly all models achieved high, tightly clustered predictive accuracy (mean calibration-phase validation *r* = 0.67 across the ten models; SD = 0.07 across model means and SD = 0.12 across the 250 site-model estimates; Figure 4A). The untrained model provided an important reference point: its lower encoding performance (*r* = 0.47) confirmed that this standard procedure can detect coarse differences in brain alignment, and its degree of predictivity replicates existing work suggesting that untrained models are capable of explaining meaningful variance in neural datasets via linear regression (53). However, among the nine trained models, natural-image predictivity offered limited separation (SD = 0.01).

**Figure 4.**
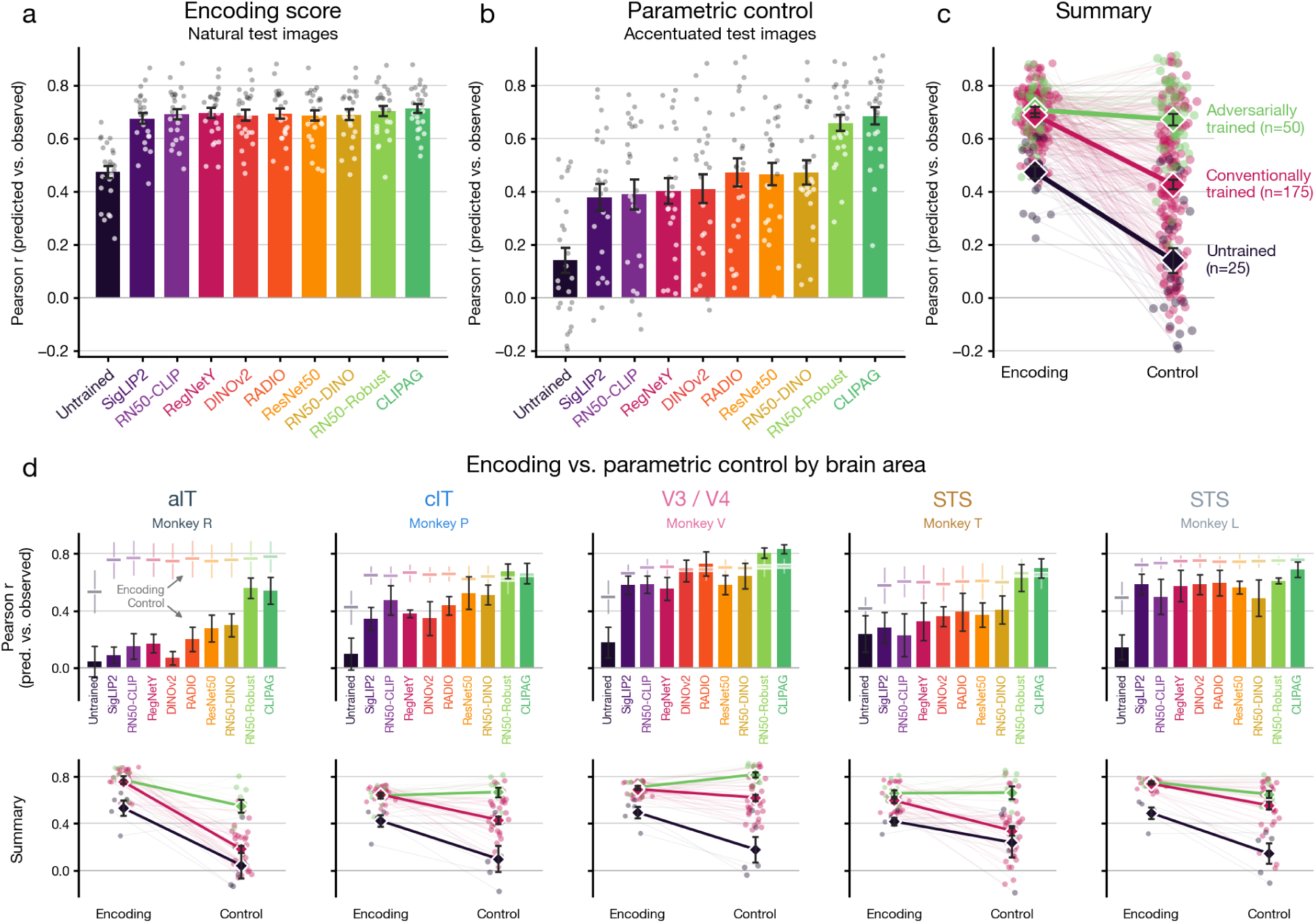
Models with saturated natural-image encoding diverge in parametric neural control. (A) Encoding performance over held-out natural images during the calibration phase. Bars show each model’s mean Pearson r across the 25 sites, dots show individual sites, and error bars indicate 1 SEM; n = 25 sites 10 models. Each color indicates one model. Across the ten model means, encoding r = 0.67 0.07. (B) Parametric control performance over accentuated stimuli. Same format as (A), but each model is evaluated on its own axis-aligned accentuated stimuli. Control predictivity is lower and more variable across models (across model means, r = 0.45 0.15). (C) Summary of Pearson correlations across the two experimental scenarios: natural-image encoding and parametric control. Each thin line indicates one model-site combination, colored by model family (adversarially trained, conventionally trained, and the untrained baseline). Large diamonds indicate family means 1 SEM, connected across conditions; only the adversarially trained family maintains high alignment under parametric control. (D) Encoding and control performance by brain area and animal. Top row shows control r over accentuated stimuli for each model, with that model’s mean calibration phase encoding r overlaid as a faint dash (thin vertical line = 1 SD across sites); bottom row summarizes the encoding-to-control change by model family, as in (C), with dots indicating individual site-model combinations. Error bars reflect 1 SEM: across the five sites of each animal for the bars (top row), and across that animal’s model-site combinations within each model-family for the grouped means (bottom row).

A different picture emerged when the same models were evaluated by their ability to modulate neural responses with accentuated stimuli (Figure 4B). Across the population, models systematically overestimated their neural control capacity: predictivity over accentuated stimuli dropped and spread substantially relative to natural images (*r* = 0.45; SD = 0.15 across model means and SD = 0.27 across the 250 model-site pairs), revealing systematic differences among models (repeated-measures ANOVA: *F* (9, 216) = 25.5, *p* = 1.4 × 10*^−^*^29^), with similar trends for the control slope outcome measure (Fig. S17).

Among the many possible model contrasts represented in our set, two models stood out in neural control: CLI-PAG and ResNet50-Robust. Although they differ in architecture, objective, and training data, both incorporate explicit adversarial robustness training, whereas none of the other included models do. Controllability varied substantially across neural sites (SD = 0.18 in mean control *r* across the 25 sites, range 0.09–0.74; Fig. S19), and several site-model combinations yielded impressive degrees of parametric control (e.g., *r* = 0.91, slope = 0.81 for CLIPAG in V3/V4, Monkey V channel 355; Fig. S22–S26).

These model differences in control were consistent across brain regions and animals. Figure 4D shows that the 9 trained models were nearly indistinguishable in brain predictivity, clustering tightly across models in both the calibration and control sessions (across-model range: calibration phase validation *r* = 0.67–0.71; control phase cross-session re-test *r* = 0.51–0.54 over re-presented held-out images, see Methods). But these models showed substantial variation in their parametric control capacity (RM-ANOVA per region: aIT *F* (9, 36) = 13.0, *p* = 7×10*^−^*^9^; cIT *F* (9, 36) = 4.8, *p* = 3×10*^−^*^4^; V3/V4 *F* (9, 36) = 9.4, *p* = 3×10*^−^*^7^; STS [Monkey T] *F* (9, 36) = 4.7, *p* = 4×10*^−^*^4^; STS [Monkey L] *F* (9, 36) = 6.6, *p* = 2×10*^−^*^5^). In general, the models that showed the best prediction of neural responses to the controller stimuli were consistent across areas (mean Spearman *ρ* = 0.69 ± 0.16 SD), and even higher when limiting analysis to aIT vs. cIT (Spearman *ρ* = 0.94); Fig. S21.

Taken together, these results reveal that many models can masquerade as similar in their brain alignment while relying on very different hierarchical computations internally. The feature accentuation process makes these differences explicit via the model-derived stimulus sweeps, which vary markedly in their ability to parametrically modulate neural responses.

### Input-gradient structure is a stronger predictor of neural control than adversarial robustness

Our ten models afford many possible contrasts, and a priori one might have expected the parametric control outcomes to organize along one of the familiar axes that distinguish them, for example separating convolutional from transformer architectures, supervised from self-supervised objectives, or language-aligned from vision-only models, with one class emerging as the more brain-aligned. We found no such dissociation: among the trained models, none of these standard groupings strongly separated neural control capacity (Fig. S30). Instead, the obvious common factor of the two best controllers was some form of explicit adversarial training (Figure 4C,D; see Figs. S31–S34 for statistical support). We confirmed this in a ‘controversial’ parametric control experiment pitting ResNet50 against its adversarially trained counterpart: on stimuli synthesized to maximize their predictive disagreement, only the adversarially trained model captured unique brain-aligned variance (see Figs. S36–S38). This result indicates that adversarial training directly increased ResNet50’s brain alignment.

How generalizable is the relationship between robustness and control? That is, does the degree of adversarial robustness of a model’s encoding axis predict its capacity for neural control across the full set of models, including those without explicit adversarial training? Somewhat surprisingly, adversarial sensitivity generalized poorly as a predictor of control beyond the adversarially trained models (Figure 5). We measured the adversarial robustness for all 250 axes, using projected gradient descent (50) on held-out natural images to maximally increase and decrease the predicted responses at each fixed perturbation budget *ɛ* (Figure 5A,B). Sensitivity correlated with control across the full model set (site-level slope *r* = −0.51, model-level *r* = −0.89) but this was carried entirely by the two adversarially-trained models and vanished once they were excluded (site-level slope *r* = −0.05, model-level *r* = −0.41; exact permutation *p* = 0.36; Figure 5D,E; see Figs. S40, S41 for similar results across alternative sensitivity measures and fast gradient sign method attacks; 54).

**Figure 5.**
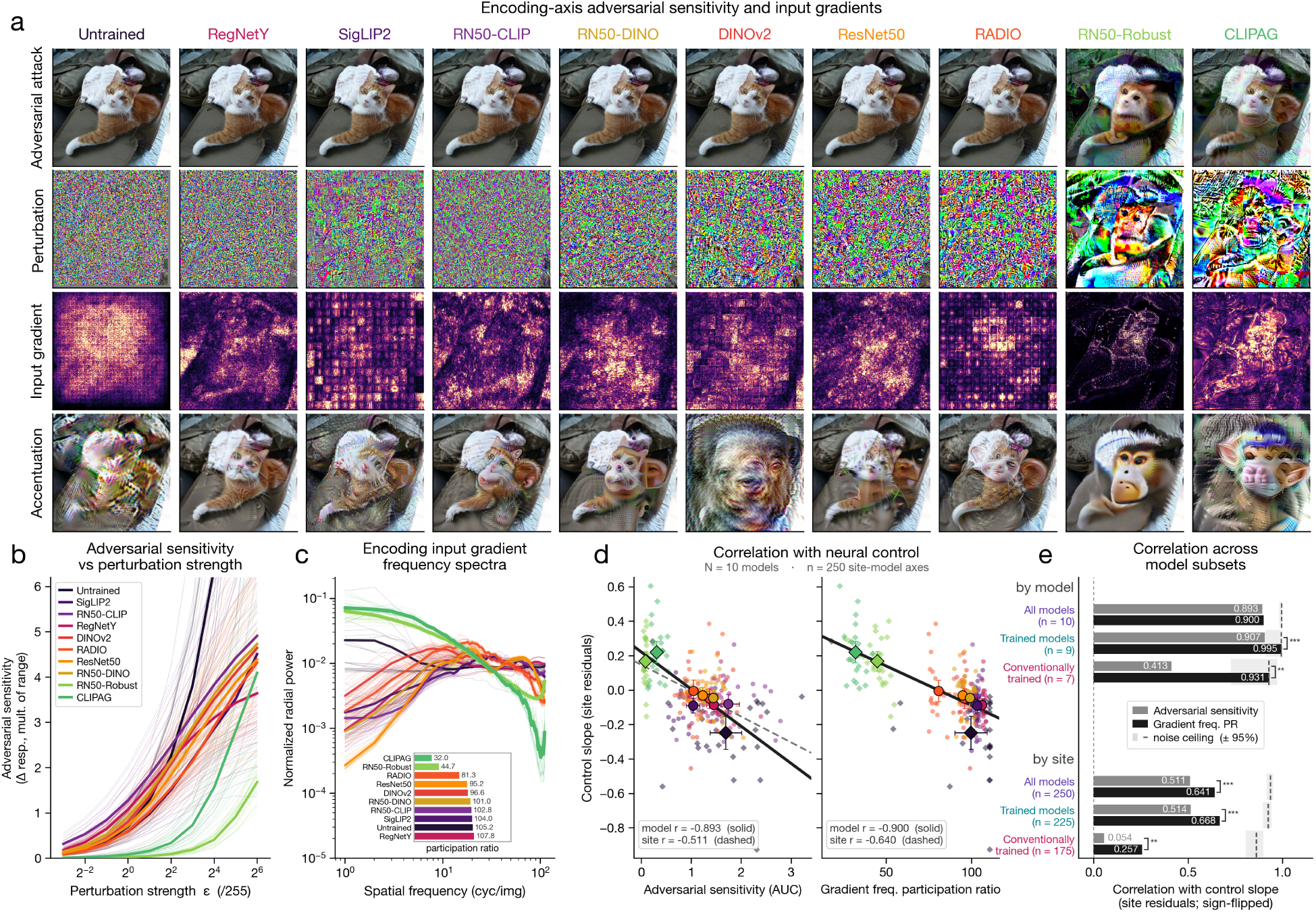
Input-gradient structure predicts parametric neural control beyond adversarial sensitivity. (A) Encoding-axis behavior for one face-selective cIT site (Monkey P) and seed image, with models ordered by gradient spectral concentration. Rows show the minimally perturbed image reaching +2 z under PGD, the corresponding perturbation, the input-gradient saliency map, and an axis-aligned accentuation reaching +3.1 z. Adversarially trained models show more coherent, low-frequency structure, whereas other models show more diffuse, high-frequency patterns. (B) Adversarial sensitivity across perturbation strengths ɛ (/255; log_2_ scale), quantified as the up-versus down-driven response swing relative to each axis’s natural response range. Thin lines indicate sites and bold lines model means; adversarially trained models are least sensitive and the untrained model most sensitive. (C) Model-mean radial power spectra of input gradients. The inset reports spectral participation ratio, with lower values indicating greater concentration; the adversarially trained models have the most concentrated spectra. (D) Site-residualized control slope versus adversarial sensitivity (log-ɛ AUC; left) and gradient spectral concentration (right) across 250 model-site axes. Small dots indicate individual axes, large dots model means ±1 SEM, diamonds the untrained and adversarially trained models, and solid and dashed lines model-and site-level fits. Both measures predict control across the full model set (site-level r = 0.51 and 0.64, respectively). (E) Correlations with control slope, with signs oriented so larger values indicate stronger prediction, computed across models (top) and sites (bottom) for all 10 models, the 9 trained models, and the 7 conventionally trained models. Dashed lines and shaded bands indicate split-half noise ceilings (±95%). Gradient spectral concentration remains predictive after excluding the adversarially trained models, whereas adversarial sensitivity does not (site-level r = 0.26 versus 0.05); brackets compare predictors (^∗^ p < 0.05, ^∗∗^ p < 0.01, ^∗∗∗^ p < 0.001).

However, we did identify a promising metric associated with neural control outcomes, related not to the overall degree of gradient-based adversarial sensitivity, but rather to the spatial structure in those gradient perturbations. Put more simply, when tracing the gradients from the encoding axis back down to the pixels, some pixels matter more than others in driving that axis. For some models, these consequential pixels were coherently focused on object and surface boundaries, while others showed more diffuse, high-frequency patterns spread across the image, and even grid structure from transformer input patchification (Figure 5A). We quantified this spatial structure with a spectral measure (based on the radial Fourier power spectrum of the input gradients over held-out natural images, using the participation ratio as a measure of low-frequency spectral concentration; see Methods). Gradient spectral concentration strongly predicted parametric neural control across the 250 individual axes (*r* = 0.64), and across the ten model means (*r* = 0.90; Figure 5D). Critically, and unlike adversarial sensitivity, the association between input gradient structure and control outcomes held even when removing the adversarially-trained models (site-level *r* = 0.26, exact permutation *p* = 0.0006; model-level *r* = 0.93, exact permutation *p* = 0.0016; Figure 5E). This relationship was robust to many variations, including alternate spectral concentration measures and probe image sets (Fig. S41, S45), and it generalized to held-out models, sites, and animals (*R*^2^ = 0.42–0.45, Fig. S48). Together, these results identify input-gradient spectral concentration as a strong prospective predictor of neural controllability.

### Implications for benchmarking and model comparison

We next turned to two practical questions: does pooling accentuated stimuli into a larger benchmark strengthen or weaken model separation, and, can carefully selected natural images achieve comparable discriminative power to synthesized stimuli?

We found that “personalized” accentuated stimuli (consisting of the 110 images synthesized separately for each model-site pair along that model’s own encoding axis) produced substantially stronger model differentiation than larger pooled stimulus sets. When all 5,500 accentuated images per monkey were aggregated into a global benchmark and every model was scored on the full set, model separation collapsed (SD = 0.04 vs. 0.11 for the personalized accentuations; Figure 6A, left), likely because most pooled images fall within a narrow band of low predicted activation for any given model (Figure 6B). Even in the pooled regime, a small but significant advantage remained for adversarially trained models (mean pooled *r* = 0.48 vs. 0.40; paired *t*(24) = 5.3, *p* = 2×10*^−^*^5^). Removing each model’s own accentuations (*n* = 550 total across five target sites) from the pooled set and evaluating on the remaining 4,950 images reduced model separation even further (SD = 0.02; Figure 6A, right), revealing that the discriminative power of the pooled benchmark derives almost entirely from the small fraction of personalized accentuated stimuli. These findings demonstrate that targeted synthetic stimulus sets have far greater diagnostic capacity than larger but untargeted natural image benchmarks.

**Figure 6.**
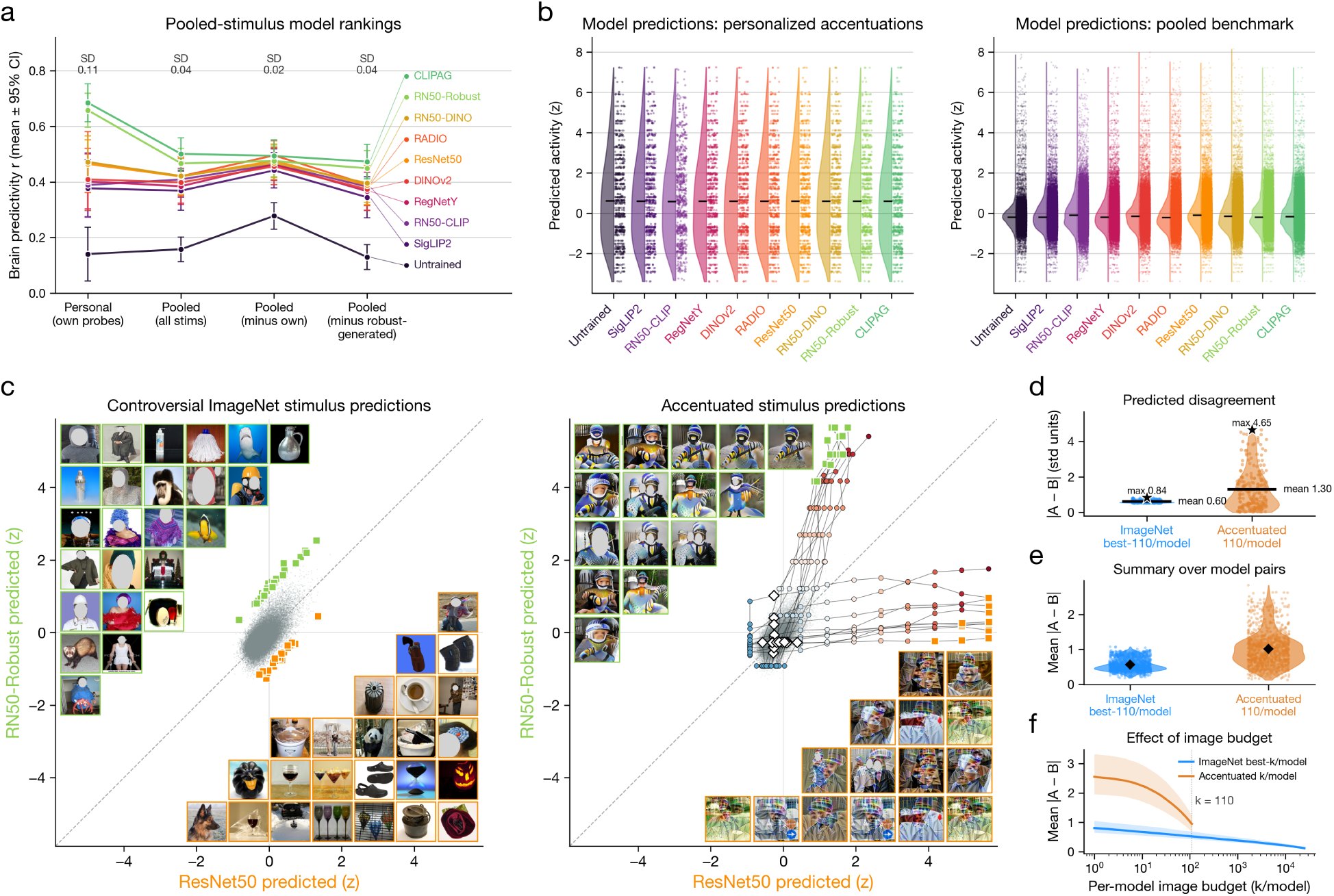
Accentuated controller stimuli differentiate models more efficiently than pooled or natural-image benchmarks. (A) Brain predictivity for each model under four evaluation regimes: personalized accentuations (n = 110 images), the pooled accentuated-stimulus benchmark (n = 5500 images), the pooled benchmark with each model’s own accentuations removed (n = 4950 images), and the pooled benchmark with all adversarially trained models’ accentuations removed (n = 4400 images). Set sizes are the design counts; all analyses use the stimuli actually presented to each animal (see Methods). Values indicate Pearson r, mean±95% CI over 25 sites. Model separation is largest for personalized accentuations and shrinks under pooling (across-model SD = 0.11, 0.04, 0.02, and 0.04 across the four regimes), but the two adversarially trained models remain top-ranked even when all their stimuli are excluded. (B) Distribution of predicted activation levels for each model over personalized accentuations and the full pooled benchmark. Predicted activations are z-scored. Personalized stimuli span broad predicted response ranges by construction, whereas most pooled stimuli cluster within a narrower low-activation range. (C) Predicted disagreement between ResNet50 and ResNet50-Robust for one example aIT site. Left: best-case ImageNet validation images selected to maximize predicted disagreement. Right: corresponding accentuated stimuli. Images are plotted by each model’s predicted response. (D) Predicted disagreement for the example model pair and site in (C), comparing the best-case ImageNet subset and accentuated stimulus set. Disagreement is measured as A B in standardized prediction units; the accentuated set produced roughly twice the mean disagreement of the best-case ImageNet subset (mean 1.30 versus 0.60, 2.1×). (E) Summary of predicted disagreement across all channel-model-pair combinations (n = 1125 combinations). Accentuated stimuli yield greater predicted disagreement than budget-matched best-case ImageNet subsets, exceeding the ImageNet baseline in 90.5% of combinations; the advantage holds after aggregating to the 25 sites (24 of 25; Wilcoxon signed-rank p = 1.2×10^−7^) and in all five animal means. (F) Mean predicted disagreement as a function of per-model image budget. Shaded regions indicate 95% confidence intervals.

Personalized accentuated stimuli also recovered far more discriminative power per image than natural images. To make the strongest possible case for natural images, we identified, for each model pair and channel, the 220 ImageNet validation stimuli (among 50,000) producing the largest predicted disagreement, matching the two models’ pooled accentuation budget (see Methods). For an example model pair (ResNet50 and ResNet50-Robust) at one aIT site, the best-case ImageNet subset achieved an average disagreement of 0.60 standardized units (see Methods), while the corresponding accentuations produced 1.30, more than twice as large (2.1×; Figure 6C–D). Across all 1,125 channel–model-pair combinations, accentuated stimuli yielded substantially greater disagreement than budget-matched best-case ImageNet subsets, exceeding the natural-image baseline in 90.5% of cases (24 of 25 sites, Wilcoxon *p* = 1.2×10^−7^; Figure 6E), and this held at every tested budget (Figure 6F). While we cannot rule out that internet-scale image sets might close this gap, feature accentuated stimuli certainly provide a vastly more data-efficient route to model comparison than natural-image search.

## Discussion

Despite near-equal encoding predictivity, the popular and top-performing DNN encoding backbones examined here varied dramatically in their ability to control neural responses via axis-aligned feature accentuation. All models overestimated their control capacity, revealing clear remaining gaps in our ability to model visual tuning in the ventral stream, with two adversarially trained models consistently showing the strongest parametric control across subjects, areas, and electrode types. This advantage was confirmed in a targeted controversial-stimulus experiment pitting standard and adversarially trained ResNet50 models directly against each other. Across the full model set, the degree of adversarial sensitivity was moderately predictive of control outcomes, but this relationship did not hold when considering only the seven conventionally trained models. Instead, control was more strongly predicted by the spectral structure of each encoding axis’s input gradients. Together, these results establish parametric neural control as a substantially more stringent test of model-brain alignment within this high-performing model set, and identify a prospective signature of control outcomes to be validated in future experiments.

Our closed-loop paradigm shifts model comparison up the ladder of causal inference (46, 55). Standard encoding procedures are observational: models are fit to responses elicited by a preexisting stimulus set, and we ask which patterns of variation they capture. Because the rich covariance structure of natural images allows many different feature sets to support equally predictive models (termed the Rashomon effect; 56–58), observational tests alone struggle to adjudicate among them. Presenting accentuated stimuli back to the animal corresponds to intervention: setting the input to a model-prescribed value and asking whether the predicted neural effect holds. On this framing, parametric axis-aligned control instantiates and tests a causal theory of what makes a neuron fire: *P* (Response|*do*(*I* = *A*(*M, λ***v**))), where an accentuation procedure *A* uses model *M* to synthesize an image at position *λ* along the encoding axis **v**. When neural responses track the predicted rates across a parametric sweep of these inputs, the model has isolated the causal effect of that stimulus dimension on the neuron. While natural-image benchmarks remain valuable for assessing coarse-scale model alignment, we contend that this interventional standard provides stronger leverage for identifying the inductive biases that drive precision model-brain alignment.

Our results also bear on current debates of representational convergence between neural systems. We observe that models tend to look more aligned under more flexible tests of similarity (Figure 4; 59, 60), while diverging under causal perturbation. This divergence qualifies recent broad claims of representational universality (12, 61–63) and the brain alignment of untrained architectures (53). Our findings instead support more tempered views in which convergence may be partial, shaped by shared ecological constraints (64), and strongly dependent on the procedure for testing alignment (e.g., the Aristotelian and Umwelt representation hypotheses; see 13, 14, 64–66). Many models appear to converge on linearly mappable features while differing in their input sensitivities and internal computations, and feature accentuation makes these algorithmic differences visible to experimenters.

### Implications of axis-dependent neural control measures for model comparison

A potential concern is that axis-aligned accentuation may inherently favor adversarially trained models, whose input gradients are argued to be more human-interpretable (28, 67). Under this view, the models themselves could be equally brain-aligned while our synthesis procedure simply extracts more effective controller stimuli from some than others. Several results argue against this explanation. First, adversarially trained models remained the strongest predictors even when evaluated only on stimuli generated by other models (Figure 6A). Second, the synthesis regularization strategies used here were originally developed to improve feature visualizations from non-robust models (49, 51), and appear to do exactly that: despite these networks’ susceptibility to imperceptible adversarial perturbations (Figure 5A,B), they produce accentuated stimuli with widespread, perceptible image changes (Figure 2, Figure 3, Figure 5A, Figs. S11, S12–S16), suggesting that our procedure, if anything, benefits conventionally trained models rather than disadvantaging them. Finally, the strongest prospective predictor of control outcomes, input-gradient spectral concentration (Figure 5D,E, Fig. S44), is an intrinsic property of the fitted model that can be computed without synthesizing accentuated stimuli. Together, these observations suggest that differences in neural control reflect properties of the models themselves rather than artifacts of our synthesis procedure.

Another question is how much these differences in neural control ultimately matter: if many models are effectively equivalent in their ability to predict responses to new natural images, what should we make of their divergent predictions for synthetic, non-ecological stimuli? The core issue at stake here is generalization: whether these models capture the true neural tuning function across the broader space of visual inputs. Encoding models can look highly successful when test images are drawn from the same distribution as the training set, especially when test images have close neighbors in the training data (68, 69). In this case, strong held-out predictive performance may reflect interpolation within nearby regions of a high-dimensional space, rather than recovery of the underlying tuning function. If the scientific goal is to identify that function, stronger tests require extrapolation and out-of-distribution generalization, for example by deliberately withholding regions or dimensions of a continuous stimulus space and asking whether the model correctly predicts responses there (70). Because image space is so vast, model-derived stimuli guarantee direct tests of the hypothesized model encoding axis. Relatedly, axis-aligned accentuation is only one way to use encoding models for targeted traversals of image space. Off-axis accentuations, for example, could probe whether the invariances predicted by an encoding axis also match the neuron’s null space (cf. 71, 28, 72). In sum, model differences that appear inconsequential for in-distribution prediction become consequential precisely when we ask the model to generalize; identifying the correct encoding axes is what makes such generalization possible.

What are the implications of these results for NeuroAI research, and particularly for the role of large-scale neural datasets? Our findings suggest that for modeling goals that depend on recovering true neural tuning, closed-loop experiments may be necessary. While such experiments can appear methodologically demanding, their requirements are increasingly tractable. In practice, they require stable recordings across multiple sessions, with model fitting and stimulus synthesis performed between sessions. Efficient, easy-to-use pipelines for generating model-driven stimulus hypotheses will further accelerate this empirical-computational loop. Fortunately, substantial progress has been made in maintaining recording stability across sessions, as demonstrated in recent large-scale standalone neural datasets (73–77). Importantly, however, closed-loop experiments do not require scale to be effective. Indeed, our results show that selectively targeting diagnostic stimuli can be more informative than broadly sampling large regions of image space. Thus, we suggest that incorporating model-derived stimuli into experimental pipelines can provide a valuable complement to large-scale, model-independent neural datasets.

### Robustness training and spectral signatures of brain alignment

A growing body of work has converged on the observation that adversarially trained models have distinctive forms of brain and behavioral alignment: they synthesize more recognizable metamers (28), generate image perturbations that shift model and human percepts in predicted directions (78, 67), modulate macaque IT activity via subtle image updates (42), and better predict human auditory responses in controversial tests against baseline models (79). Our results offer a refinement of what adversarial training is doing and which properties of the resulting models matter for brain alignment.

Adversarial sensitivity captures the overall magnitude of perturbation required to change a model’s prediction, but although it separated the robust models from the rest, it did not explain graded variation in control among conventionally trained models (Figure 5D,E). The stronger predictor instead captured the spatial organization of influential pixels, rather than only their aggregate sensitivity, providing a more nuanced characterization of how models are sensitive to subtle input changes. On this view, adversarial training is one route, currently the most effective, for producing this more brain-aligned gradient structure, which appears to provide a more general predictor of neural control than overall robustness itself (see related work on Perceptually Aligned Gradients; 80–84).

Given that we have identified a candidate prospective predictor of control, it will be valuable to test whether it generalizes across other models with mechanisms that are known to impact robustness. For example, several inductive biases including blurred (85) or noisy training regimes (86, 87), feedback architectures (88), extra-classical receptive fields (89), enhanced viewpoint diversity (90), and more developmentally realistic visual diets (91) all show representations that are naturally more adversarially robust. What is the impact on the input-gradient spectral structure of these models’ encoding fits? Using existing neural datasets, these models can be easily pre-screened for competitive natural-image brain predictivity and the degree of concentrated input-gradient spectra, and tested on the accentuated stimuli that are legible to the strongest-performing models identified here. This provides a clear path for asking which inductive biases yield the representational and gradient properties associated with successful neural control, while reserving new neural experiments for the most promising candidates.

However, we caution against interpreting well-structured input gradients as the core mechanism underlying effective neural control. A substantial proportion of variance in control remained unexplained by the predictors tested here (Fig. S48, S49), and spatial-frequency reliance can be dissociated from adversarial robustness (92, 80; see also 93). The gradient spectra may therefore serve as a proxy for aspects of hierarchical model-brain alignment that have not yet been characterized (94). Identifying those properties will likely require resolving the specific model subcircuits that support successful neural control, aided by advances in pruning (95) and interpretability (96–100) together with richer neural datasets.

## Conclusion

How should we identify the best models of the brain and neural processing? While the field will surely continue to produce many valid and defensible approaches, here we endorse a method that focuses on how precisely a model can capture neural tuning: a neuron’s particular set of invariances and sensitivities to variation across all of input space. The central tenet of this approach is that the tuning of individual neurons is not arbitrary, but instead reflects the computational role of that unit within the larger basis set of units contributing to visual processing. The current work shows that image space is simply too vast to fully characterize neural tuning across a diverse population of neurons by randomly sampling natural images, but that axis-aligned feature accentuation can translate encoding models into focused sweeps through natural image space that span predicted neural response levels, revealing whether the encoding axis is truly brain aligned. While leading DNN encoding models have saturated on neural predictivity for natural images, our method reveals that most are substantially misaligned with the brain, while a few show promising degrees of neural control in some neural sites. We propose that targeting precision neural control is a path toward more extensively capturing the tuning of neurons across the ventral stream. Doing so will surely impact theories of visual processing, but it also promises to improve prediction and control of the downstream behavior of the system, which would be important for practical applications such as neural decoding and neural prosthetics.

## Materials and Methods

### Animal subjects, neural recording, and behavioral task

#### Subjects and recording targets

We recorded from five awake, behaving macaques (*Macaca mulatta*; with abbreviated names “R,” “P,” “L,” “T,” and “V”), three male and two female, with body weights ranging from 3 to 14 kg. All procedures were approved by the Harvard Medical School Institutional Animal Care and Use Committee (protocol #ISO00001049) and conformed to NIH guidelines provided in the *Guide for the Care and Use of Laboratory Animals*.

Neural activity was acquired using two electrode platforms across the five subjects. To enable stable longitudinal recording, two animals were implanted with chronic multi-electrode arrays in inferotemporal (IT) cortex: a 64-channel microwire array (MWA; Microprobes for Life Science, Gaithersburg, MD) targeting left anterior IT (aIT) in Monkey R, and a 64-channel floating microelectrode array (FMA; Microprobes) targeting left central IT (cIT) in Monkey P. To bias recording toward reliable, category-selective populations in these two animals, implant sites were guided by functional magnetic resonance imaging (fMRI)-localized face-selective regions (Fig. S4). For dense, high-yield sampling in the remaining three animals, Neuropixels probes (Neuropixels 1.0, 383 channels, NHP Long, 45 mm; IMEC, Leuven, Belgium; 101) were used: CT-and MRI-guided implantations in Monkeys L and T targeted the left superior temporal sulcus (STS), starting in V1 and traversing through STS, whereas in Monkey V a single probe traversed V1, V2, V3, V4, posterior IT (pIT), and the upper bank of the STS, with analyzed units restricted to V3 and V4 (Fig. S3).

#### Behavioral task and stimulus presentation

To maintain stable gaze, each monkey performed a fixation task while seated in a primate chair, receiving juice-drop rewards as long as fixation was held within a 1^◦^ window around a central dot. Stimuli were presented on a calibrated liquid-crystal display (LCD) monitor 57 cm away from the animal. Gaze location was monitored continuously using infrared eye-tracking (ISCAN, Woburn, MA). Deviations from the target were detected using this system and this facilitated either delivering or withholding reward. Trials where fixation was broken were discarded online. Experimental control was implemented using the NIMH MonkeyLogic software (102). To evoke reliable, spatially matched responses, stimuli were presented in a rapid serial visual presentation (RSVP) fashion at 4–6 visual degrees and centered on each site’s mapped receptive field (RF); on every trial an image was shown for 250 ms followed by a 250 ms inter-stimulus interval (ISI), yielding a rapid serial sequence that maximized the number of unique stimuli per fixation while preserving a fixed stimulus-response epoch across sessions.

#### Signal preprocessing and normalization

To capture spiking activity at high temporal fidelity from the chronically implanted arrays, broadband FMA and MWA signals were amplified and sampled at 40 kHz with a Plexon acquisition system, and spiking events were detected online with OmniPlex software using a threshold-crossing criterion at 3.5 standard deviations. Spike timings were then binned into spike counts at 1 kHz. Most detected events reflected multi-unit spiking; where single units were isolated by waveform clustering, they were merged back into the corresponding channel to yield channel-level activity.

For the Neuropixels recordings in Monkeys L, T, and V, broadband data were acquired at 30 kHz using SpikeGLX software (Janelia Research) and preprocessed using CatGT. To obtain a continuous, threshold-free measure of aggregate spiking near each electrode, neuronal activity was quantified using the multi-unit activity envelope (MUAe), computed with custom code following established methods (103, 104): bandpass filtering (300–3000 Hz), rectification, lowpass filtering (200 Hz), and downsampling to 1 kHz. Because the MUAe captures a continuous and instantaneous envelope of local spiking (105, 106), it lives on a different scale from discrete spike rate and thus requires its own normalization.

To place all signals on a common temporal resolution and extract evoked responses, the raw neuronal activity, whether spike counts or MUAe, was smoothed with a 25 ms boxcar running average and then binned into 10 ms non-overlapping bins. For each stimulus, a scalar response was quantified by averaging activation in a fixed temporal window post-stimulus onset: [100, 400) ms for the array recordings in Monkeys R and P, and [80, 250) ms for the Neuropixels probes in Monkeys L, T, and V. These windows were chosen by grid searching over different options using pilot data and identifying the regime that maximized response reliability within each of the two recording regimes. To align response distributions across sessions (justified by the finding that neuronal selectivity in chronic recordings is highly stable), single-trial scalar responses within each recording day were pooled to estimate a per-channel mean *µ* and standard deviation *σ*, and each response was z-scored to a common scale:

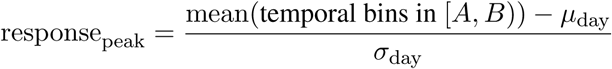

The z-scored activities were then concatenated across multiple recording days, and repeated trials per stimulus were averaged, yielding the average response matrix (stimuli×neuronal channels) that served as the regression target for all encoding models.

For the control (accentuation) sessions, the identical peak-window averaging, smoothing, and binning procedures were applied. There were two modifications for this phase. First, to remove artifacts, we applied a per-session, per-channel outlier rejection procedure. Specifically: a trial was discarded if, for any channel, its robust z-score, |*x* median|*/*(1.4826 · MAD), exceeded 15. Median and median absolute deviation (MAD) were computed over that channel for that session. Second, because we expected response gain to drift across the multi-day control phase of the study, we standardized the values in each control session using a fixed set of anchor images. For anchors we used the shared natural calibration images re-presented during the control phase, standardizing each session on all of those presented that day. For each session and channel, the mean and SD of single-trial responses over these anchor stimuli were used to compute z-score statistics that were then applied to that session’s activity. In some cases a session contained fewer than five anchor repeats, due to limited stimulus presentations. In these cases, the z-scoring statistics were estimated over all of that day’s trials, including accentuated stimuli.

We used these standardized z units to fit all encoding models and compute all control scores, and assess the resulting correlations and slope fits. As such, channels differing in baseline firing and response gain were able to be assessed and compared on a reasonably common scale. Where an interpretable firing-rate axis was needed, for display and for defining the firing floor described below, the standardization was inverted as spk*/*s^∗^ = *z σ* + *µ*, recovering the evoked rate in spikes per second (denoted spk/s^∗^, though MUAe activations reflect analog values and are not strictly spikes). Because the per-session statistics differ, we implemented this backward conversion by taking the average per-channel *µ* and *σ* across the relevant phase’s sessions, with values floored at 0 spk/s^∗^ given that firing rates cannot be negative.

### Closed-loop experimental design

#### Four-phase paradigm

To test whether deep neural network (DNN) encoding models matched in natural-image predictivity nonetheless differ in their causal control over primate visual cortex, we implemented a closed-loop paradigm in which each of ten candidate DNN models was used to synthesize graded stimuli predicted to parametrically drive or suppress the firing of individual recorded neural sites (Figure 1). The experiment was organized into four phases. In a first calibration (mapping) phase, the responses of each targeted neural site (a reliably recorded, responsive channel) were recorded to a fixed calibration set of natural and object-segmented images. In a second, offline phase carried out on a compute cluster, these responses were used to fit a per-site, per-model encoding axis. In a third, likewise offline phase, model-derived feature-accentuation stimuli were synthesized from each fitted axis (49, 51). In the fourth (control) phase, responses were recorded to the synthesized accentuation sweeps, graded stimulus sets designed to modulate predicted responses within and beyond the natural-image response range, so that the parametric controllability conferred by different models could be compared directly at the same sites.

Encoding axes were fit and control stimuli generated for 25 neural sites across the five macaques (Monkeys R, P, L, T, and V), spanning areas V3, V4, central and anterior inferotemporal cortex (cIT, aIT), and the superior temporal sulcus (STS). Calibration recordings were collected over consecutive-day sessions (3 for Monkey R, 3 for Monkey P, 6 for Monkey L, 6 for Monkey T, and 4 for Monkey V), each presenting the same 969 stimuli; control recordings followed over 4–9 sessions per animal, each beginning 4–8 days after that animal’s final calibration session (Table 1).

**Table 1.** Neural data acquired during the encoding and control experiments. Per animal: recording dates, session counts, total trials, unique stimuli, and mean trial repeats for the encoding (calibration) and control (accentuation) experiments, the number of recorded units, and the gap between the final encoding session and the first control session.

| Monkey | Enc. dates | Enc. sess. | Enc. trials | Enc. stim. | Enc. avg. rep. | Units | Ctrl. dates | Ctrl. sess. | Ctrl. trials | Ctrl. stim. | Ctrl. avg. rep. | Gap (d) |
| --- | --- | --- | --- | --- | --- | --- | --- | --- | --- | --- | --- | --- |
| R | Apr 28–30 | 3 | 8,241 | 969 | 8.50 | 64 | May 4–12 | 9 | 24,691 | 5,575 | 4.43 | 4 |
| P | Apr 28–30 | 3 | 3,164 | 969 | 3.27 | 64 | May 5–12 | 8 | 14,051 | 5,182 | 2.71 | 5 |
| L | Apr 26–May 1 | 6 | 4,736 | 969 | 4.89 | 383 | May 9–12 | 4 | 3,040 | 2,498 | 1.22 | 8 |
| T | Apr 26–May 1 | 6 | 6,002 | 969 | 6.19 | 383 | May 8–12 | 5 | 4,097 | 3,057 | 1.34 | 7 |
| V | Apr 26–29 | 4 | 16,260 | 969 | 16.78 | 383 | May 7–12 | 6 | 24,526 | 5,595 | 4.38 | 8 |

#### Calibration image set and data partitions

To sample both naturalistic scene statistics and isolated objects, the 969-image calibration set (Fig. S1) was assembled from three sources: 649 images from the Natural Scenes Dataset (NSD) Shared-1000 set of natural scenes (73), 259 from a set of segmented objects and animals on white backgrounds, and 61 from a functional localizer (fLoc) set (107). The identical set was presented to all five monkeys and used for encoding-model fitting, model evaluation, and cross-session normalization (Figure 1). For all fitting and evaluation we used a single fixed partition into 774 training images, on which encoding weights were fit, and 195 held-out validation images, used both for per-channel layer selection and for the reported encoding scores. Since we used this same set of held-out images to guide layer selection, we note that all calibration-phase encoding scores reported here reflect these post-selection validation scores rather than fully independent scores assessed using an entirely different set of held-out stimuli. To monitor the stability of brain responses and encoding axes across experimental phases (and help standardize responses to a common scale), we relied on a 100-image subset of the calibration set comprising 50 training and 50 validation images, drawn in a manner that roughly matched the composition of the full set across the three image sources. Both halves of this 100-image subset were re-presented during the control phase and together formed the anchoring set: each control session was standardized on all of the calibration images actually presented that day, rather than on a fixed subset. All 100 were re-presented in Monkeys R, P, and V, whereas Monkeys L and T received 49 and 46 respectively, reflecting their incomplete stimulus sets. To estimate cross-phase encoding generalization (the control-phase retest Pearson *r*), we used only the held-out validation images among these re-presented stimuli (50 for Monkeys R, P, and V; 24 for Monkey L; 22 for Monkey T), so that no model-training images contribute. These scores therefore reflect generalization to held-out stimuli measured in a separate session, several days after the encoding models were fit.

#### Channel reliability and site selection

To restrict closed-loop targeting to the most stable channels, we quantified reliability for every channel using the calibration recordings. We used two different measures: a split-half procedure that correlated odd-and even-trial condition means across the 969 calibration conditions, followed by Spearman–Brown correction; and a complementary noise-ceiling signal-to-noise ratio (“NCSNR”; 73) estimated over conditions that had at least three repeats. These two estimates of reliability were highly convergent. Five channels with high calibration reliability were retained per animal (Monkey R [0, 2, 9, 15, 19], Monkey P [0, 8, 24, 40, 47], Monkey V [9, 79, 151, 331, 355], Monkey L [81, 282, 286, 306, 342], and Monkey T [56, 74, 120, 168, 204]), yielding 25 total sites. To encourage that our control experiments would target sites with non-redundant tuning, a candidate site was excluded if its tuning profile over the 969 calibration images correlated above *r* = 0.9 (Pearson) with an already-selected site. The noise-ceiling reliability of the 25 selected sites is reported in Fig. S2. To confirm stable tuning across the transition to control, per-channel Pearson correlations were computed between calibration and control responses over the shared natural images re-presented in both phases (Fig. S2).

### Deep neural network models

#### Desiderata for model selection

To sample the space of leading computational models of primate high-level vision, we assembled a fixed set of ten DNN models spanning a range of architectures and training objectives. Four desiderata guided this selection. First, we included the standard vision encoders most widely used as backbones in neural encoding studies (for example, ResNet50). Second, we included models empirically shown to predict high-level visual responses well, whether or not they are widely adopted in the field, drawing on the BrainScore benchmarks (11) and on the large-scale Natural Scenes Dataset (NSD) fMRI model benchmark of Conwell et al. (5). Third, we incorporated several models that were receiving widespread attention in the fields of computer vision and cognitive neuroscience as of Spring 2025. Finally, when possible we selected models that shared some inductive bias: either a training set (e.g. ImageNet), architecture (e.g. ResNet50), or task, such that differences in neural control could be better attributed to specific learning pressures.

#### Overview of the chosen models

The ten models were as follows. **ResNet50** was included as the field’s de facto standard image encoder. An adversarially trained **ResNet50-Robust** was included both as a top-performing BrainScore model of inferotemporal (IT) cortex and because adversarially trained models have been studied extensively in relation to the human brain and behavior (28, 42). **ResNet50-CLIP**, the top-scoring model of human occipitotemporal cortex (OTC) responses in the Conwell et al. (5) NSD benchmark and itself widely studied against visual brain data, was included as a contrastive language-image variant on the same convolutional backbone, and **CLIPAG** (ViT-B/32) as an adversarially robustified fine-tuning of CLIP. **ResNet50-DINO**, a self-supervised convolutional network and a precursor to DINOv2, was included to span self-supervision on the shared ResNet50 backbone. Among transformer-based systems, we included **DINOv2** (ViT-B/14, with register tokens), a self-supervised visual foundation model of rapidly growing influence; **SigLIP2** (ViT-B/16), a language-image model whose sigmoid loss enables very large training batch sizes; and **RADIO v2.5-B**, a multi-task vision model that distills several foundation models, including DINOv2 and SigLIP, into a single backbone. The convolutional **RegNetY-640** (a SEER-trained RegNet) was included as another top performer in the Conwell et al. (5) benchmark. Finally, **AlexNet** was selected because, as of 2 May 2025, it was the overall top-ranked model averaged across the BrainScore neural_vision benchmarks, driven by its leading performance on the V1 and V2 benchmarks available at that time; as of 21 July 2026 it remains the top-ranked model on the V1 benchmarks. On inspection, however, the model variant served through the BrainScore API (“alexnet_training_seed_01”) carries weights inconsistent with a correctly trained network (near-zero weight correlation to a reference AlexNet across all layers, and untrained-level classification and linear-probe accuracy, in contrast to a correctly loaded checkpoint and the standard torchvision AlexNet, which both reach normal accuracy), and it is precisely this effectively untrained variant that attains those high BrainScores. As such, the model functions in our study as an “**Untrained**” baseline, while still being relevant as a “leading” model in light of its performance on BrainScore. The architecture and training objective of each model are summarized in Table 2.

**Table 2.** Deep neural network models and their training objectives. Architecture family (convolutional network, CNN; vision transformer, ViT) and the training/objective properties of each of the ten models.

| Model | Architecture | Supervised classification | Self-supervised / contrastive | Adversarial training | Distillation | CLIP / language alignment |
| --- | --- | --- | --- | --- | --- | --- |
| Untrained (AlexNet) 108, 109 | CNN |  |  |  |  |  |
| ResNet50 110 | CNN | ✓ |  |  |  |  |
| ResNet50-Robust 50, 111 | CNN | ✓ |  | ✓ |  |  |
| ResNet50-CLIP 112 | CNN |  |  |  |  | ✓ |
| ResNet50-DINO 113 | CNN |  | ✓ |  |  |  |
| RegNetY 114, 115 | CNN |  | ✓ |  |  |  |
| CLIPAG 116 | ViT |  |  | ✓ |  | ✓ |
| SigLIP2 117 | ViT |  |  |  |  | ✓ |
| DINOv2 118, 119 | ViT |  | ✓ |  |  |  |
| RADIO v2.5 120, 121 | ViT |  |  |  | ✓ |  |

#### Model groupings

To summarize and further examine this post hoc observation, we grouped the models according to architecture and the presence of explicit adversarial robustness training. The three families were: convolutional neural networks without adversarial training (CNNs; ResNet50, RegNetY-640, ResNet50-CLIP, ResNet50-DINO), vision transformers without adversarial training (DINOv2, SigLIP2, and RADIO v2.5), and adversarially trained models (ResNet50-Robust and CLIPAG). We treated the untrained AlexNet control as its own group. These families therefore isolated adversarial training as a property of interest while enabling us to compare models whose baseline natural-image predictivity was approximately matched (Figure 4; Figure 5).

### Encoding-model fitting

#### Feature extraction and dimensionality reduction

To predict the firing of each neural site using each candidate DNN, we fit an encoding model that mapped model features onto the site’s repeat-averaged responses to the calibration stimulus set. For each model, post-activations were extracted from major network blocks, rather than from every constituent internal computational stage. Because the flattened block activations were high-dimensional, we used linear dimensionality reduction before regression to stabilize the fits and reduce overfitting: principal component analysis (PCA) was used to project each flattened layer activation onto its top 750 components. We set the PCA dimensionality to 750, just below the maximum rank of 774 imposed by the number of calibration images for training, thereby capturing nearly all available variance. We flattened and transformed the full spatial feature tensor for convolutional networks, and for vision transformers all tokens (patch, register, and class) were flattened and concatenated before PCA.

#### Ridge regression and per-channel layer selection

To map the PCA-reduced features onto neural responses, we fit ridge regression with built-in cross-validation (RidgeCV function; scikit-learn, 122) on the 774 training images over a 14-value logarithmic penalty grid spanning 10^−4^ to 10^9^, with a separate regularization value selected for each recording channel via RidgeCV’s efficient leave-one-out generalized cross-validation. We used the 195 validation images from the calibration set to identify the most predictive layer of every model, per channel separately, by measuring the coefficient of determination (*R*^2^ between predicted and observed brain responses; Fig. S5); we report each model’s encoding score as the validation-set Pearson correlation and *R*^2^ between predicted and measured held-out responses (Figure 3; Figure 4).

Because both PCA and ridge regression are linear, the resulting encoding model is an end-to-end differentiable function spanning from pixel inputs to the measurement sites. We directly take advantage of this property for our accentuation synthesis. Each image is preprocessed and passed through the frozen backbone to the encoding source layer, where its latent representation is transformed by the fitted PCA projection and passed through the ridge readout to compute the predicted response for a given unit, while preserving gradient flow back through the entire graph.

### Feature accentuation and controller-stimulus synthesis

#### Overview of axis-aligned feature accentuation

To synthesize stimuli predicted to parametrically drive or suppress activity at each site while preserving natural-image structure, we used a feature-accentuation procedure that optimizes an image to elicit a specified response from a fitted encoding model (49, 51). Because each encoding model is a linear read-out of deep-network features, optimizing this scalar response amounts to displacing the image to a particular degree along the fitted encoding axis (Figure 2).

Generating a single accentuated stimulus for a given site-model pairing involved gradient-based optimization where an input “seed” image was iteratively passed through the model and updated via back-propagation to achieve a target level of predicted neural activity. Each synthesis was initialized from a natural seed image resized to 224×224 px, scaled to [0, 1], and upsampled to a 1024×1024-px optimization canvas. A given forward pass involved first transforming the image using the model’s native preprocessing (resize to 224×224 px and channel normalization, with a bicubic resize plus a 224-px center crop for the ViT-, CLIP-, RegNet-, and SigLIP-family backbones), then passing it through the frozen backbone up to the fitted layer, and through the fitted PCA projection and ridge read-out, yielding the scalar predicted response of the target unit as the objective *f* (x). Using auto-differentiation machinery, we computed the gradient of this objective towards pixel inputs ∇_x_*f* (x), and thereby optimized the image to drive the predicted response toward a specified target level. Repeating this optimization procedure until convergence and sweeping across a graded series of target levels produced one “accentuated image sweep” per site × model × seed image.

#### Regularization of stimulus synthesis

Optimizing an image directly against a frozen encoding model can exploit non-natural, high-frequency directions and yield noisy or otherwise uninterpretable visualizations unless appropriate regularization is imposed (123–125). Several complementary forms of regularization were used to constrain and improve the quality of image synthesis (Fig. S6). Note that no explicit pixel budget term was used; regularization arose entirely from the image parameterization and the augmentations described here.

##### Decorrelated-spectrum parameterization

Rather than optimizing pixels, images were parameterized in a decorrelated Fourier space, following the feature-visualization lineage of Olah et al. (125). The seed was mapped into this space by min-max normalization to [0, 1], an inverse-sigmoid (logit) transform, RGB decorrelation via a fixed color matrix, and a forward 2D FFT; at each step the spectrum was inverted back to an image by applying an inverse 2D FFT, spatial-mean subtraction, RGB re-correlation, a sigmoid nonlinearity, and rescaling to [0, 1].

##### Frequency-decay reweighting

To further suppress spurious high-frequency detail, Fourier-space gradients were reweighted by a frequency-decay envelope proportional to *f* ^−^*^α^*, with the decay exponent *α* tunable over 1.2–2.5. This has been shown to dramatically improve the quality of feature visualizations in prior work (51).

##### Stochastic crop augmentation

To promote robustness of the accentuated features to small viewing changes, each optimization step evaluated the objective over a batch of eight stochastically augmented crops, whose centers were jittered and whose extents were drawn from a box-size range of [0.90, 0.95], each resampled to 224 × 224 px and corrupted by additive noise of tunable amplitude 0.05–0.30 (51, 49).

##### Asymmetric objective and early stopping

To terminate image synthesis once the prediction target for a given accentuation instance was reached, we relied on an asymmetric objective. Specifically, we scored the clean, full-resolution accentuated image at each step and once the predicted activation level exceeded the target for the first time (either in the positive direction for drive stimuli or in the negative direction for suppress stimuli), we reversed the synthesis direction with tenfold reduced update magnitude, until the score fell within 0.01 absolute magnitude of the target. The stimulus was synthesized using NAdam at a learning rate of 12.0 for up to 6,000 steps, stopping early once the above criterion was reached, and the result was saved at 1024 × 1024 px.

#### Target schedule

To place sweeps from different models on a common, response-referenced scale, we defined an 11-level target schedule for each neural site. The lower and upper anchors were the 1st and 99th percentiles (*q*_01_ and *q*_99_) of the site’s repeat-averaged responses across the 969 encoding images. Defining the response bandwidth as the difference between *q*_99_ and *q*_01_, the full schedule extended 0.25 times the bandwidth below *q*_01_ and 0.50 times the bandwidth above *q*_99_, thereby probing both within-and beyond-range activation. The 11 levels were evenly spaced across this interval, yielding a fixed split of six within-range and five beyond-range levels, with two on the suppression side and three on the drive side.

#### Automated hyperparameter selection

To ensure that syntheses consistently reached their precise predicted activation targets, we developed an automated procedure that was used to identify a suitable regularization regime independently for every model-channel combination (10 models×25 channels) before the full set of accentuated stimuli were generated. The tested hyperparameter pairs specified a noise-augmentation magnitude and a frequency-decay exponent *α* (see above); larger values impose stronger regularization. Each regime was scored by synthesizing accentuations across a 3×11 grid of three seed images crossed with all 11 target levels and counting a cell successful when its final prediction fell within 0.1 of target. The candidate regularization levels were ordered from most to least strong regularization (e.g. highest noise augmentation and steepest frequency-decay exponent run first), and the search stepped down this list and stopped at the first ladder step where the success rate over all 33 cells exceeded 95%. This prioritized maintaining a relatively strong level of regularization that would still enable targets to be reached consistently. If no regime succeeded against this criterion, the highest-scoring set of hyperparameters was retained. This search was run independently for each model-site combination (Fig. S6).

#### Final dataset overview

To provide natural starting points for each accentuation sweep that were not used during model fitting, ten seed images were drawn exclusively from the 50-image re-test subset of held-out validation images, spanning the natural-image response range and balanced 6 animate images (containing people or animals) with 4 containing only inanimate content. Synthesizing an accentuation for every seed × site × level combination yielded 5 sites×10 seeds×11 levels = 550 stimuli per model, 5,500 per subject, and 27,500 across the five macaques, each resized to 425×425 px for presentation to match the resolution of the NSD calibration stimuli. For three site-model combinations from Monkey L, a duplicated synthesis run produced a second image variant for 277 different design cells. Of these, one variant entered the presented set for 139 and both variants for 51 (the remainder were never presented due to the monkey not completing the full experiment). Because image synthesis involves some degree of randomness, we compared each variant pair at the pixel level. Those that reconverged to essentially the same final output (pixel-wise *r* ≥ 0.90; 57 of the 277) were collapsed onto the presented variant, and where both variants of a pair were presented (10 cells), the responses were pooled and treated as additional trial repeats. The pairs that diverged into distinct images were retained as extra unique stimuli. For reference, even adjacent levels of the same sweep, which are distinct stimuli by design, never exceeded a pixel-wise *r* of ≈ 0.74. A synthesis was scored as reaching its target when its miss, the absolute difference between the achieved (re-encoded) prediction and the target level, fell within 5% of that site’s target-schedule range; by this criterion 94.5% of syntheses reached target (Fig. S7). We conducted various analyses and exclusions to ensure fairness in model comparison (see below), given that some individual accentuations failed to reach their synthesis targets and that some stimuli were not presented during control sessions due to monkeys not finishing the experiment.

### Encoding-axis alignment of accentuation sweeps

To assess how faithfully the accentuation sweeps traversed their fitted encoding axis, we decomposed each sweep’s net latent displacement into a component lying along the axis and a component lying off it, following the on-axis/off-axis analysis of 42. Across the full set of 2,500 sweeps (25 sites×10 models×10 seed images), each sweep was represented in the 750-dimensional PCA feature space by its net displacement *d* between the most-suppressive and most-driving levels, and its encoding axis by the unit read-out vector a = *w/||w||* This displacement was split into the directional modulation (DM), the magnitude of travel along the encoding axis, and the direction-orthogonal modulation (DOM), the residual travel over the 749 dimensions orthogonal to the axis:

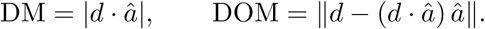

Because DOM aggregates 749 dimensions where as DM measures a single one, the two were also compared on a per-dimension basis (DM against 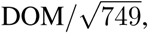 the scaling expected if the displacement *d* was spread isotropically in 750d, 42). Finally, we asked whether the encoding axis was the dominant individual direction of traversal. For each model-site pair, we first projected out the component parallel to the encoding axis (*a*) from the centered activation matrix over the calibration stimuli, then ran PCA on the resulting 749-dimensional residual data, yielding a new orthonormal basis of off-axis directions. We then projected each sweep displacement **d** onto these directions, and the on-axis magnitude of DM was compared with the largest absolute projection onto any one of the off-axis components (Fig. S9).

### Quantifying neural control

#### Outcome measures

We quantified neural control outcomes by computing two complementary summary statistics using the accentuated image responses for each site-model combination. For the main analyses reported here, we pooled each model’s accentuations for a given site across all ten seed images and eleven target levels before computing the summary statistics, rather than scoring each seed sweep separately and averaging sweep-level scores. Our results broadly do not depend on this choice (Fig. S18). We defined the control score as the Pearson correlation (*r*) between the model’s predicted responses and the measured neural responses over this pooled set of stimuli, and the control gain as the slope of the ordinary-least-squares regression of measured responses on predicted responses. Because both quantities were computed in z-scored response space, with predicted responses on the calibration-phase z scale and measured responses on the control-phase z scale, these slope values were directly comparable across experimental sites. An outcome score was returned as undefined if fewer than three stimuli were available for evaluation.

Because overall controllability varied substantially across neural sites (Fig. S19, S20), analyses relating encoding-axis properties to control across the 250 site × model experiments (e.g., adversarial sensitivity and gradient spectral concentration; Figure 5) used site-residualized outcomes: from each site-model control score or slope, we subtracted that site’s mean outcome computed over the nine trained models. This step removes any shared site-level variation in reliability and/or intrinsic controllability that may otherwise dominate the observed effects. After residualization, the remaining variation necessarily reflects differences among models tested at the same sites. We excluded the untrained model’s data from the computation of the per-site reference mean so that its uniformly poor control could not have an outsized effect on the distribution within which the trained models were compared.

#### Noise ceiling and ceiling-normalized control

To estimate the maximum control correlation expected given trial-to-trial response variability, we computed a Pearson-r noise ceiling for each stimulus set using the signal/noise decomposition of 73. Let *r* denote a single-trial response and *r̄* the response averaged across trials for each stimulus. We first estimated the trial-noise variance as the average within-stimulus variance across repeated trials,

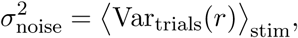

and the total variance across the trial-averaged stimulus responses as

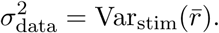

Because 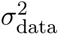 is computed from trial-averaged responses, it contains only a reduced contribution from trial noise. We therefore estimated the stimulus-driven signal variance as

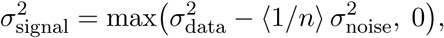

where *n* is the number of repeats for a stimulus and 〈1*/n*〉 is the mean reciprocal repeat count across stimuli (equal to 1*/n* when all stimuli have the same number of repeats).

We then defined the noise-ceiling signal-to-noise ratio as

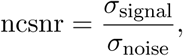

from which the expected Pearson correlation between the measured trial-averaged responses and their underlying noise-free responses follows as

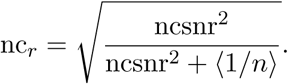

Equivalently, 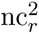 gives the variance-explained noise ceiling described in 73 (Fig. S2).

For each stimulus set, trial-noise variance was estimated from same-day repeats of the anchor images, rather than from the accentuated images, which were largely presented only once. A noise ceiling could not be estimated for Monkeys L and T because their STS recordings contained insufficient same-day anchor repeats. Ceiling-referenced analyses were therefore restricted to Monkeys R, P, and V, comprising 150 of the 250 site × model combinations.

Estimating these ceilings allowed us to express control outcomes as a fraction of the maximum level that could in principle be attained given the signal-to-noise ratio of brain responses to a given model’s accentuated images, for a target site (Fig. S31).

#### Firing floor and correction of predicted responses

All of our control metrics relied on post-hoc predicted responses obtained by passing each presented image back through its generating model’s frozen backbone, PCA projection, and ridge readout. Because the target levels per model extended below the calibration range for each neural site (see above), it was possible that some predicted response levels in standardized (z) units could imply negative firing rates. We therefore clamped predictions at a site-specific firing floor before computing control outcome measures. In z units, the firing floor was defined as the more negative of the session-averaged z-score corresponding to zero firing (-*µ/σ*) and the minimum z-score observed across calibration trials, the latter verified to correspond to true zero-spike epochs, so that the floor never sits above a level the neuron actually reached. Thus, for a subset of accentuated images, each prediction was replaced by max(prediction, floor). This clamp was applied only to model predictions, not measured responses, and only at the point of comparing those predictions with brain responses. It affected only chronic-array spiking data from Monkeys R and P (floor∼ -1 to 2 z), and not Neuropixels MUAe data from Monkeys V, L, and T (floor∼ -10 to 25 z). Control results were essentially unchanged whether sub-floor predictions were clamped, left unmodified, or excluded.

#### Model differentiation

To ask whether models diverged in their parametric control capacity, control scores were computed over the full accentuation sweeps for each of the 25 neural sites crossed with all ten models, pooling the presented accentuated stimuli from the different seeds together for each site-model combination. No synthesis target-achievement filter was applied for this main comparison, such that all presented stimuli contributed to the outcome measures. Per-model control performance and analysis across model families and cortical areas is summarized in Figure 4.

To test whether model differentiation depended on the personalized accentuated stimuli obtained from each model, we compared model separation across three probe regimes of increasing scope. Such analyses were possible for two reasons: first, any presented image can be scored by any model post hoc; and second, because all targeted sites were recorded simultaneously, every accentuated image elicited measured responses across all sites, regardless of the site for which it was synthesized. The regimes were: “personalized”, where each model was scored only on its own accentuations for its own site; “pooled”, where each model was scored on all of the accentuations targeting sites from a given animal (a benchmark roughly an order of magnitude larger); and “pooled-minus-own”, the pooled set with a model’s own accentuations removed.

In each regime, differentiation between models was summarized as the across-model standard deviation of the control scores (Figure 6A). To ensure fairness for this analysis, the pooled predicted-response distributions were evaluated only using the (site, seed, level) cells at which all ten models were on-target within a strict tolerance (2% of the channel range) on the firing-floor-clamped achieved responses. This ensured that no model’s accentuations reached further along the response axis than another’s (Figure 6B).

As a further index of model control capacity, we identified super-stimuli: images that drove firing outside the natural-image range to levels above *q*_99_ were labeled super-drive and those below *q*_01_ were labeled super-suppress stimuli. We summarized the super-stimulus proportion across all 25 sites and ten models (Figs. S27–S29).

#### Filtering for fair cross-model comparison

Because a small proportion of accentuations failed to reach their synthesis targets, and not all stimuli were presented to every animal, we implemented several exclusion regimes to perform model comparison in a way that would ensure fairness. As a baseline, every presented stimulus was included regardless of whether it hit its target. Then, we defined three exclusion regimes. The “per-model” regime entailed that each model’s off-target accentuations were dropped, retaining only its own successful syntheses. In the “level-drop” regime, entire target levels were excluded by their scheduled rank rather than by whether models achieved them or not. This ensured that all models were scored over the same range of predicted response levels. In the strictest (“intersection”) regime, a stimulus was retained only if all ten models successfully reached the accentuation target at that same site, seed, and level combination, such that no model was scored over a wider response range than any other. The structure of the underlying synthesis failures is documented in Fig. S8, and the retention and control outcomes under each exclusion regime are reported in Fig. S35.

### Controversial-accentuation experiment

To adjudicate between two models whose natural-image predictivity was closely matched, we adapted the controversial-stimulus method of Golan et al. (26) to the accentuation setting (Fig. S36). We sought to create a set of controversial accentuated stimuli, where two candidate models would make divergent predictions, and then the neural response would effectively discriminate between them. As such we replaced our single-model accentuation objective with a dual objective defined as the difference between two models’ predicted responses for the target unit. The stimulus synthesis procedure was modified to create images that would drive up one model’s readout while suppressing the other’s, with the procedure repeated and the sign reversed in order to create two different controversial scenarios, one favoring the predictions of each model.

This procedure relied on the same decorrelated-spectrum parameterization and random crop augmentation as the single-model accentuation pipeline, but this time we did not use a graded target level schedule nor an early-stopping criterion. Instead, each image was optimized for a fixed 2,048 NAdam steps (learning rate = 5.0) at a stimulus size of 768 × 768 pixels. We used a relatively heavy regularization regime (noise level = 0.30, spectral decay rate *α* = 2.5), and the final output of each one of these runs was retained as the test stimulus.

Our experiment contrasted a standard ResNet50 against its adversarially trained variant, ResNet50-Robust. We chose these because of their shared architecture and training task, and roughly matched encoding predictivity, given that the key variable would be whether adversarial robustness training was employed. For both models, the encoding axis was fit from the same model layer (layer 4, block 1), over PCA-reduced features, with cross-validation used to identify the optimal regularization *λ* per site as in the main experiments. For our controversial test, we targeted seven aIT sites in Monkey R (channels 1, 9, 15, 16, 25, 37, and 44). For each site we generated one controversial accentuated stimulus per model over 10 different seed images, yielding 7×2×10 = 140 total images (Fig. S37). We followed the same two-phase experimental flow, where accentuations were presented in a control phase that was subsequent to the initial calibration phase. Encoding axes for this experiment were fit with cross-validated Lasso regression (in place of the RidgeCV procedure used elsewhere), and the same Lasso readouts were used for scoring. We compared the models’ performance by scoring each stimulus within both encoding readouts, such that each model would have a predicted response level for all 140 stimuli. The key outcome measure was whether the brain responses over the 20 site-targeted accentuations better tracked the predictions of baseline ResNet50 or ResNet50-Robust. As such, there were 40 total model-image-response datapoints available to plot per neural site (Fig. S36, S38). We summarized these outcomes with Pearson *r* scores as well as OLS slope fits, and tested for significant differences between models over the seven sites with a Wilcoxon signed-rank test (chosen over a paired *t*-test given the small number of sites).

### Analysis of adversarial sensitivity and input gradient spectra

#### Adversarial sensitivity of encoding axes

To quantify the degree to which each encoding axis was sensitive to small pixel-level input perturbations, we relied on standard procedures for deriving adversarial attacks. This analysis leveraged 100 held-out NSD natural scene images that were not used elsewhere in the study.

For each encoding axis and image, we performed projected gradient descent (PGD, *L*_∞_ norm, for 50 optimization steps) to maximally increase (and separately decrease) the predicted neural response in a given site. We repeated this procedure across a grid of fixed perturbation budgets *ɛ*, that were log-spaced from 0.125*/*255 to 16*/*255. At each budget level, for each model-site encoding axis, we measured the resulting difference between the increases and decreases in response achieved by the strongest attacks (i.e., the response “swing”). We then standardized these swing values by expressing them as fractions of that axis’ natural (calibration) response range, such that they would be directly comparable across models and sites. This yielded a trajectory of different sensitivity values across *ɛ* levels, per model-site axis. To summarize adversarial sensitivity for each of these, we considered many different summary statistics (Fig. S41), and results were not strongly dependent on this choice. As a default, we relied on the area under this normalized swing curve (AUC) across log *ɛ*; lower values here denote a more perturbation-robust axis (Figure 5; Fig. S39, S40). These supplementary figures further document whether the relationship between robustness and control is dependent on the attack norm (*L*_∞_ vs. *L*_2_), the attack method (PGD vs. fast gradient sign method; FGSM), and the integration window over *ɛ* values.

#### Input-space gradients

To characterize the input-level gradient structure of each encoding model, we analyzed the spatial frequency properties of gradient maps derived from each of the 250 encoding models. To do so, we computed the gradient of each model’s predicted neural response with respect to the input image pixels, which is well defined because each encoding model is end-to-end differentiable. We used 100 held-out NSD stimuli that were not used in the main experiments to assess these gradient properties, and the same set was used to measure adversarial sensitivity (see above). Each gradient map comprised one 3×224×224 (RGB×height × width) tensor per image and therefore we obtained a 100×3×224×224 stack per site×model, across all 25 sites and all ten models. For the example galleries, gradients were additionally evaluated at the ten natural seed images along which each accentuation sweep steps; for the gallery site (Monkey P, cIT, channel 8), each gradient was rendered as a saliency map, the L2 magnitude across the color channels, shown beside that model’s top accentuated stimulus for the same seed (Figure 5A; Fig. S43).

#### Spectral analysis of gradients

To place these gradients on a common spectral axis, each per-image gradient map was reduced to a radially averaged 2-D Fourier power spectrum (a 2-D FFT followed by radial binning of |*F*|^2^ over integer-radius rings about the image center, with frequency in cycles per image), and the 100 held-out-image profiles were averaged per site×model. This computation is illustrated step by step in Fig. S42. A seed-based estimate computed identically over the ten synthesis seeds agrees almost perfectly with the held-out-image estimate (*r* = 0.98 across the 250 axes; Fig. S45). As a natural-image reference, the identical computation was applied to the 969 calibration images. A standard way to characterize spectral signatures like these is the slope of their decay in log-log space, which presumes that power falls off as a power law of spatial frequency (1*/f^α^*). Visual inspection indicated that these gradient spectra were not well described by a single power law, so for our main analyses we summarized each spectrum by its participation ratio (PR),

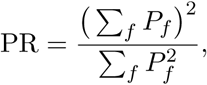

the effective number of frequency bins over which the radial power *P_f_* is spread, computed over radial frequencies from 1 to 111 cycles/image (from just above the DC term up to the 112 cycles/image along-axis Nyquist; the DC term and the sparsely sampled corner frequencies beyond the Nyquist are excluded; bins spaced evenly on a linear axis, one per integer-radius ring, each the average power over that ring).

Intuitively, the PR equals *N* for a perfectly uniform (white) spectrum over *N* bins and decreases toward 1 as power concentrates into fewer bins. We chose this measure as the default spectral summary statistic due to several advantages: it is scale-invariant and assumes nothing about the shape of the spectrum (whereas e.g. a linear slope fit to estimate rate of spectral decay in log-space would require stronger assumptions).

That said, we note that the PR measures the unevenness of the spectrum across frequency bins, and does not specify where the power is concentrated; a spectrum peaked at high frequencies could in principle also yield a low PR. In our study, we observe that lower PR closely tracks lower spectral centroid and steeper rates of decay, indicating greater low-frequency content (Figure 5C). Where a signed “concentration” predictor is reported, it is oriented so that higher values denote greater concentration (lower PR). We also tested several other measures of spectral characteristics for a relationship with neural control outcomes (e.g. coefficient of variation, Wiener spectral flatness, normalized spectral entropy, median frequency, and log-log spectral-decay slope; see 126–128; Fig. S41, S45–S47). Results were broadly unchanged across these different summary measures.

#### Predicting parametric control

We compared several different predictors in their association with downstream control outcomes. Our primary analyses focused on the 250 site-model experiments (25 sites×10 models), where control outcomes (Pearson *r* and slope) were residualized by subtracting each site’s mean, isolating differences among models within sites from variation in overall controllability across sites.

Beyond our main analysis of gradient spectra and adversarial robustness (Figure 5), we examined the following larger set of predictors: alternative gradient-spectrum measures (e.g. spectral coefficient of variation, Wiener flatness), alternative adversarial sensitivity measures, calibration-and control-phase encoding accuracy, properties of the accentuated stimuli such as their spatial frequency content, readout-layer effective dimensionality, ImageNet linear-probe accuracy, and encoding axis weight norm (Fig. S49).

For Figure 5D–E, predictor-control relationships were quantified with Pearson correlation across site-residualized model-site axes and across model means, using three different model groupings: all 10 models, the nine trained models, and the seven conventionally trained models. Significance was then assessed by permuting model identities while preserving the crossed site × model structure (see below). Correlation noise ceilings were estimated by repeatedly splitting each experiment’s 10 seeds, recomputing control outcomes in each half, and applying the Spearman–Brown correction.

To assess out-of-sample generalization with pre-specified models, we separately fit fixed one-and two-predictor linear models of the site-residualized control outcomes over the nine trained models: gradient spectral concentration alone, adversarial sensitivity alone, and each with natural-image encoding accuracy added, using a shared intercept and global slopes, with predictors z-scored within each training fold. Out-of-fold *R*^2^ was evaluated under progressively harder partitions: random 10-fold, leave-one-model-out, leave-one-site-out, and leave-one-animal-out (Fig. S48). Cross-validated *R*^2^ values were referenced to a split-half explainable-variance ceiling for each outcome, estimated by repeatedly dividing each experiment’s accentuated stimuli into halves, recomputing the outcome for each half, correlating the resulting estimates across experiments, and applying the Spearman–Brown correction; this ceiling represents the maximum coefficient of determination attainable given measurement noise.

### Disagreement of natural versus accentuated stimulus predictions

We asked whether the personalized accentuations were capable of separating pairs of encoding models more effectively than an equal number of natural images (Figure 6). For each of the 25 sites and 45 unique model pairs, we computed the predicted responses to the pair’s 220 personalized accentuations: 110 generated for each model at that site (10 seeds×11 levels). Predictions from each model–site encoding axis were then standardized using its own 1st–99th percentile response range over the 50,000 ImageNet-1k validation images (129), to facilitate direct comparison between all models. Disagreement was then calculated image by image as the absolute difference between the two standardized predictions, |*A − B*|, one value obtained from each model in the pair.

In Figure 6E,F, we scaled this analysis to the full set of 25×45 = 1,125 total unique site-model-pair combinations: we compared the mean disagreement over the 220 personalized accentuations with that over the 220 ImageNet images producing the largest |*A − B*| from among the full validation set. For the specific example in Figure 6C,D, the ImageNet set was instead balanced directionally, using the 110 images most favoring each model, for the sake of visualization. These predicted-disagreement analyses use all synthesized accentuations, whereas the measured-response benchmarking (Figure 6A,B) uses only the stimuli actually presented to each animal.

### Statistical analysis

#### Overview

Statistical analyses were carried out in Python using the NumPy and SciPy libraries, and, for repeated-measures ANOVA, the statsmodels library. Correlations are Pearson unless specified as Spearman rank correlations, and all hypothesis tests were two-sided. Uncertainty is reported as specified in each figure: the standard error of the mean, 95% confidence intervals from image-level bootstrap resampling, or a bootstrap standard error over experiments for cross-validated statistics. Because the comparisons below each address a distinct, pre-specified hypothesis over a small number of model families, cortical areas, or sites, *p*-values are reported without correction for multiple comparisons; as a stronger test, the key family effects were additionally required to hold in direction across every animal and area.

#### Repeated-measures ANOVA and discriminability index

To quantify whether the ten models differed in their control outcomes, we fit a one-way repeated-measures ANOVA to the per-experiment control scores (Pearson control *r*), with model as the within-subject factor (10 levels) and neural site treated as the repeated subject (25 sites; using the statsmodels AnovaRM function). Then, the same repeated-measures design was fit separately for each of the five monkeys (model factor, 5 sites per area) to confirm that model differences held region by region (Figure 4B,D). As a simple and complementary measure of model differentiation, we calculated the standard deviation of the per-model mean control scores.

#### Group and cross-area comparisons

To compare model families, we averaged control scores within the adversarially trained family and within the standard model families at each site, and the two family means were compared across the 25 sites with a paired *t*-test (we report outcomes both including and excluding the Untrained control from the standard model group). An identical paired contrast was applied to the ceiling-normalized control scores (Figure 4D; Fig. S31).

To quantify how consistent model rankings were between monkeys and areas, the per-model mean control score was computed within each of the five monkeys, and rank correlation between areas was summarized as the pairwise Spearman correlation across the ten area pairs (mean ± SD), together with the anterior-versus-central IT pair (Figure 4; Fig. S21).

In the controversial-accentuation experiment, we compared the two candidate models’ per-site prediction accuracies across the seven targeted sites with a Wilcoxon signed-rank test (see Controversial-accentuation experiment; Fig. S38).

To test whether the accentuated stimuli separated models more effectively than large sets of natural images, we compared each model-pair’s level of predicted disagreement against the best-case disagreement arising from a budget-matched subset of ImageNet validation stimuli. To test for significant levels of disagreement, we averaged over the 45 model-pair difference values per site and then tested for an effect across the 25 sites with a Wilcoxon signed-rank test, with the five animal means checked for sign consistency; the 1,125 site × model-pair combinations were used only for the descriptive fraction of combinations in which accentuation exceeded the natural-image baseline (Figure 6).

#### Model-level permutation inference

We used model-label permutation tests to test for relationships between different predictors (e.g. gradient spectral concentration or adversarial sensitivity) and parametric control outcome scores. This approach accounted for the crossed 25-site×10-model design by treating models as the key unit of inference rather than the 250 individual site-model observations, which were not independent (this avoids potentially inflating effect sizes).

For the model-level permutation tests, we permuted the model labels assigned to the predictor values and recomputed their correlation with the fixed model-level control outcomes. For site-level tests, we did the same model-label permutation across all sites before recomputing the correlation with the site-residualized control outcome values. This approach preserved the crossed site × model structure. We then calculated two-sided *p*-values as the proportion of permuted correlations that were at least as extreme as the observed correlation. We calculated permutation *p*-values from the exact null distribution obtained by evaluating all 7! = 5,040 assignments of the seven conventionally trained models. These exact permutation tests were used for the model-subset values reported in the main text and Fig. S41C.

## Acknowledgments

We thank all members of the Harvard Vision Lab for helpful conversations throughout the project and Greta Tuckute for feedback on the manuscript. We also thank Nikolas McNeal, N. Apurva Ratan Murty (Georgia Tech), Carlos R. Ponce (HMS), David Klindt, Benjamin Cowley (CSHL), Sandro Romani, Ann Hermundstad, Stephan Saalfeld (Janelia), Tim Gardner (U. Oregon), A. David Redish, Thomas Naselaris and Kendrick Kay (U. Minnesota) for inspiring questions and discussions during our presentations of early versions of this work.

## Author contributions

Conceptualization: JSP, BW, TF, AVJ, GAA, TK. Methodology: JSP, BW, TF, AVJ, MSL, TK. Investigation: AVJ, MSL. (NHP electrophysiology recording by MSL; data preprocessing led by AVJ; encoding models led by BW; feature accentuation led by TF; data analysis led by JSP.) Visualization: JSP, BW, TF, PAV, TK. Supervision: GAA, TK. Writing – original draft: JSP, BW, PAV, TK. Writing – review and editing: JSP, BW, TF, GAA, MSL, TK.

## Data, code, and materials availability

Experimental stimuli were drawn from three sources: the Shared-1000 subset of the Natural Scenes Dataset (NSD; 73; http://naturalscenesdataset.org), the fLoc functional localizer set (107; http://vpnl.stanford.edu/fLoc), and a set of segmented objects and animals presented on white backgrounds, which originated in the stimulus set of Konkle et al. (130) and was drawn here from the subset used by Vinken et al. (131). The ImageNet-1k validation split (129; https://image-net.org) was used as the large natural-image benchmark against which the accentuated stimuli were compared. All ten deep network models were used as publicly released, without retraining or fine-tuning; their architectures, training objectives, and source references are given in Table 2, and the corpora on which each model was pretrained are documented in those references. Neural recordings, the fitted per-site encoding models, and all synthesized stimuli (27,720 accentuation stimuli and the 140 controversial-accentuation stimuli) will be made available upon publication. All code needed to reproduce the analyses and figures in this paper is available at https://github.com/jacob-prince/parametric-neural-control.

## References

[1] D. L. K. Yamins, H. Hong, C. F. Cadieu, E. A. Solomon, D. Seibert, and J. J. DiCarlo. Performance-optimized hierarchical models predict neural responses in higher visual cortex. Proceedings of the National Academy of Sciences, 111(23):8619–8624, 2014.

[2] U. Güçlü and M. A. J. van Gerven. Deep neural networks reveal a gradient in the complexity of neural representations across the ventral stream. Journal of Neuroscience, 35(27):10005–10014, 2015.

[3] R. M. Cichy, A. Khosla, D. Pantazis, A. Torralba, and A. Oliva. Comparison of deep neural networks to spatio-temporal cortical dynamics of human visual object recognition reveals hierarchical correspondence. Scientific Reports, 6:27755, 2016.

[4] S. A. Cadena, G. H. Denfield, E. Y. Walker, L. A. Gatys, A. S. Tolias, M. Bethge, and A. S. Ecker. Deep convolutional models improve predictions of macaque V1 responses to natural images. PLoS Computational Biology, 15(4):e1006897, 2019.

[5] C. Conwell, J. S. Prince, K. N. Kay, G. A. Alvarez, and T. Konkle. A large-scale examination of inductive biases shaping high-level visual representation in brains and machines. Nature Communications, 15(1):9383, 2024.

[6] N. Kriegeskorte. Deep neural networks: a new framework for modeling biological vision and brain information processing. Annual Review of Vision Science, 1:417–446, 2015.

[7] D. L. K. Yamins and J. J. DiCarlo. Using goal-driven deep learning models to understand sensory cortex. Nature Neuroscience, 19(3):356–365, 2016.

[8] B. A. Richards, T. P. Lillicrap, P. Beaudoin, Y. Bengio, R. Bogacz, A. Christensen, C. Clopath, R. P. Costa, A. de Berker, S. Ganguli, C. J. Gillon, D. Hafner, A. Kepecs, N. Kriegeskorte, P. Latham, G. W. Lindsay, K. D. Miller, R. Naud, C. C. Pack, P. Poirazi, P. Roelfsema, J. Sacramento, A. Saxe, B. Scellier, A. C. Schapiro, W. Senn, G. Wayne, D. Yamins, F. Zenke, J. Zylberberg, D. Therien, and K. P. Kording. A deep learning framework for neuroscience. Nature Neuroscience, 22(11):1761–1770, 2019.

[9] A. Doerig, R. P. Sommers, K. Seeliger, B. Richards, J. Ismael, G. W. Lindsay, K. P. Kording, T. Konkle, M. A. J. van Gerven, N. Kriegeskorte, and T. C. Kietzmann. The neuroconnectionist research programme. Nature Reviews Neuroscience, 24(7):431–450, 2023.

[10] L.-M. Schmitt, D. T. Dong, E. Çelik, F. P. de Lange, and M. Toneva. A decade of comparing brains to DNNs: Progress and perspectives. Trends in Cognitive Sciences, 2026.

[11] M. Schrimpf, J. Kubilius, H. Hong, N. J. Majaj, R. Rajalingham, E. B. Issa, K. Kar, P. Bashivan, J. Prescott-Roy, F. Geiger, K. Schmidt, D. L. K. Yamins, and J. J. DiCarlo. Brain-score: Which artificial neural network for object recognition is most brain-like? bioRxiv, page 407007, 2018.

[12] M. Huh, B. Cheung, T. Wang, and P. Isola. Position: The platonic representation hypothesis. In Proceedings of the 41st International Conference on Machine Learning, volume 235 of Proceedings of Machine Learning Research, pages 20617–20642, 2024.

[13] F. Gröger, S. Wen, and M. Brbić. Revisiting the platonic representation hypothesis: An aristotelian view. arXiv preprint arXiv:2602.14486, 2026. Accepted at ICML 2026.

[14] A. S. Koepke, D. Zverev, S. Ginosar, and A. A. Efros. Back into plato’s cave: Examining cross-modal representational convergence at scale. arXiv preprint arXiv:2604.18572, 2026.

[15] F. R. Doshi, T. Fel, T. Konkle, and G. Alvarez. Visual anagrams reveal hidden differences in holistic shape processing across vision models. In *Advances in Neural Information Processing Systems*, volume 38, pages 84840–84867, 2025.

[16] M. Raghu, T. Unterthiner, S. Kornblith, C. Zhang, and A. Dosovitskiy. Do vision transformers see like convolutional neural networks? Advances in Neural Information Processing Systems, 34:12116–12128, 2021.

[17] S. Kornblith, M. Norouzi, H. Lee, and G. Hinton. Similarity of neural network representations revisited. In Proceedings of the 36th International Conference on Machine Learning, volume 97, pages 3519–3529. PMLR, 2019.

[18] J. Feather, M. Khosla, N. A. Ratan Murty, and A. Nayebi. Brain-model evaluations need the NeuroAI Turing test. arXiv preprint arXiv:2502.16238, 2025.

[19] J. S. Prince, C. Conwell, G. A. Alvarez, and T. Konkle. A case for sparse positive alignment of neural systems. In ICLR 2024 Workshop on Representational Alignment (Re-Align), 2024.

[20] M. Khosla and A. H. Williams. Soft matching distance: A metric on neural representations that captures single-neuron tuning. In Proceedings of UniReps: the First Workshop on Unifying Representations in Neural Models, volume 243, pages 326–341. PMLR, 2024.

[21] S. Shah and M. Khosla. Representational alignment across model layers and brain regions with multi-level optimal transport. In The Fourteenth International Conference on Learning Representations, 2026.

[22] C. Kapoor, A. H. Williams, and M. Khosla. Partial soft-matching distance for neural representational comparison with partial unit correspondence. In International Conference on Learning Representations, 2026.

[23] S. Muzellec and K. Kar. Reverse predictivity for bidirectional comparison of neural networks and biological brains. Nature Machine Intelligence, 8(3):474–488, 2026.

[24] I. Thobani, J. Sagastuy-Brena, A. Nayebi, J. Prince, R. Cao, and D. Yamins. Model-brain comparison using inter-animal transforms. arXiv preprint arXiv:2510.02523, 2025. Extended version of a paper in the Proceedings of the 2025 Cognitive Computational Neuroscience conference.

[25] B. R. Cowley, P. L. Stan, J. W. Pillow, and M. A. Smith. Compact deep neural network models of the visual cortex. Nature, 652(8111):947–954, 2026.

[26] T. Golan, P. C. Raju, and N. Kriegeskorte. Controversial stimuli: Pitting neural networks against each other as models of human cognition. Proceedings of the National Academy of Sciences, 117(47):29330–29337, 2020.

[27] R. Rajalingham, E. B. Issa, P. Bashivan, K. Kar, K. Schmidt, and J. J. DiCarlo. Large-scale, high-resolution comparison of the core visual object recognition behavior of humans, monkeys, and state-of-the-art deep artificial neural networks. Journal of Neuroscience, 38(33):7255–7269, 2018.

[28] J. Feather, G. Leclerc, A. Mądry, and J. H. McDermott. Model metamers reveal divergent invariances between biological and artificial neural networks. Nature Neuroscience, 26(11):2017–2034, 2023.

[29] C. Guo, M. Lee, G. Leclerc, J. Dapello, Y. Rao, A. Mądry, and J. DiCarlo. Adversarially trained neural representations may already be as robust as corresponding biological neural representations. In Proceedings of the 39th International Conference on Machine Learning, volume 162 of Proceedings of Machine Learning Research, pages 8072–8081. PMLR, 2022.

[30] R. Geirhos, P. Rubisch, C. Michaelis, M. Bethge, F. A. Wichmann, and W. Brendel. ImageNet-trained CNNs are biased towards texture; increasing shape bias improves accuracy and robustness. In International Conference on Learning Representations, 2019.

[31] K. Hermann, T. Chen, and S. Kornblith. The origins and prevalence of texture bias in convolutional neural networks. Advances in Neural Information Processing Systems, 33:19000–19015, 2020.

[32] M. Kato and B. J. He. Systematic image perturbations reveal persistent gaps between human and machine vision. bioRxiv, 2026.

[33] C. R. Ponce, W. Xiao, P. F. Schade, T. S. Hartmann, G. Kreiman, and M. S. Livingstone. Evolving images for visual neurons using a deep generative network reveals coding principles and neuronal preferences. Cell, 177(4):999–1009.e10, 2019.

[34] P. Bashivan, K. Kar, and J. J. DiCarlo. Neural population control via deep image synthesis. Science, 364(6439):eaav9436, 2019.

[35] E. Y. Walker, F. H. Sinz, E. Cobos, T. Muhammad, E. Froudarakis, P. G. Fahey, A. S. Ecker, J. Reimer, X. Pitkow, and A. S. Tolias. Inception loops discover what excites neurons most using deep predictive models. Nature Neuroscience, 22(12):2060–2065, 2019.

[36] Z. Gu, K. Jamison, M. R. Sabuncu, and A. Kuceyeski. Human brain responses are modulated when exposed to optimized natural images or synthetically generated images. Communications Biology, 6(1):1076, 2023.

[37] M. M. Henderson, A. F. Luo, S. Park, M. J. Tarr, and L. Wehbe. Diffusion-based stimulus optimization reveals functional organization across higher visual cortex. bioRxiv, 2026.

[38] Y. Yamane, E. T. Carlson, K. C. Bowman, Z. Wang, and C. E. Connor. A neural code for three-dimensional object shape in macaque inferotemporal cortex. Nature Neuroscience, 11(11):1352–1360, 2008.

[39] C.-C. Hung, E. T. Carlson, and C. E. Connor. Medial axis shape coding in macaque inferotemporal cortex. Neuron, 74(6):1099–1113, 2012.

[40] B. Wang and C. R. Ponce. Tuning landscapes of the ventral stream. Cell Reports, 41(6):111595, 2022.

[41] B. Wang and C. R. Ponce. Neuronal tuning aligns dynamically with object and texture manifolds across the visual hierarchy. Nature Neuroscience, 29(4):864–875, 2026.

[42] G. Gaziv, S. Goulding, A. Ayvazian-Hancock, Y. Bai, and J. J. DiCarlo. Noninvasive precision modulation of high-level neural population activity via natural vision perturbations. arXiv preprint arXiv:2506.05633, 2025.

[43] A. F. Luo, M. M. Henderson, L. Wehbe, and M. J. Tarr. Brain diffusion for visual exploration: Cortical discovery using large scale generative models. In *Advances in Neural Information Processing Systems*, volume 36, 2023.

[44] G. Tuckute, A. Sathe, S. Srikant, M. Taliaferro, M. Wang, M. Schrimpf, K. Kay, and E. Fedorenko. Driving and suppressing the human language network using large language models. Nature Human Behaviour, 8(3):544–561, 2024.

[45] M. W. Shinkle and M. D. Lescroart. Visualizing and controlling cortical responses using voxel-weighted activation maximization. In Proceedings of the IEEE/CVF Conference on Computer Vision and Pattern Recognition (CVPR) Workshops, pages 4864–4868, 2025.

[46] J. Pearl. Causality: Models, Reasoning, and Inference. Cambridge University Press, 2 edition, 2009.

[47] A. Motiwala, J. Soldado-Magraner, A. P. Batista, M. A. Smith, and B. M. Yu. Brain–computer interfaces as a causal probe for scientific inquiry. Trends in Cognitive Sciences, 30(1):40–53, 2026.

[48] P. Papale, D. De Luca, and P. R. Roelfsema. Deep generative networks reveal the tuning of neurons in it and predict their influence on visual perception. bioRxiv, 2024.

[49] C. Hamblin, T. Fel, S. Saha, T. Konkle, and G. Alvarez. Feature accentuation: Revealing ‘what’ features respond to in natural images. arXiv preprint arXiv:2402.10039, 2024.

[50] A. Mądry, A. Makelov, L. Schmidt, D. Tsipras, and A. Vladu. Towards deep learning models resistant to adversarial attacks. In International Conference on Learning Representations, 2018.

[51] T. Fel, T. Boissin, V. Boutin, A. Picard, P. Novello, J. Colin, D. Linsley, T. Rousseau, R. Cadène, L. Goetschalckx, L. Gardes, and T. Serre. Unlocking feature visualization for deep network with MAg-nitude constrained optimization. In Advances in Neural Information Processing Systems, volume 36, pages 37813–37826, 2023.

[52] S. Santurkar, A. Ilyas, D. Tsipras, L. Engstrom, B. Tran, and A. Mądry. Image synthesis with a single (robust) classifier. In Advances in Neural Information Processing Systems, volume 32, pages 1260–1271, 2019.

[53] A. Kazemian, E. Elmoznino, and M. F. Bonner. Convolutional architectures are cortex-aligned de novo. Nature Machine Intelligence, 7(11):1834–1844, 2025.

[54] I. J. Goodfellow, J. Shlens, and C. Szegedy. Explaining and harnessing adversarial examples. In International Conference on Learning Representations, 2015.

[55] S. Gershman. Trading places: What happens when neuroscience turns into machine learning, and machine learning turns into neuroscience? The Transmitter, Mar. 2026.

[56] L. Breiman. Statistical modeling: The two cultures (with comments and a rejoinder by the author). Statistical Science, 16(3):199–231, 2001.

[57] C. Rudin, C. Zhong, L. Semenova, M. Seltzer, R. Parr, J. Liu, S. Katta, J. Donnelly, H. Chen, and Z. Boner. Position: Amazing things come from having many good models. In Proceedings of the 41st International Conference on Machine Learning, volume 235, pages 42783–42795. PMLR, 2024.

[58] J. S. Bowers, G. Puebla, S. Thorat, K. Tsetsos, and C. J. H. Ludwig. On the misuse of prediction in NeuroAI: A case study of Centaur. PsyArXiv preprint, 2026.

[59] I. Avitan and T. Golan. Model–behavior alignment under flexible evaluation: When the best-fitting model isn’t the right one. In Advances in Neural Information Processing Systems, volume 38, pages 12081–12120, 2025.

[60] C. Kapoor, S. Srivastava, and M. Khosla. Bridging critical gaps in convergent learning: How representational alignment evolves across layers, training, and distribution shifts. In Advances in Neural Information Processing Systems, volume 38, pages 155718–155732, 2025.

[61] Z. Chen and M. F. Bonner. Universal dimensions of visual representation. Science Advances, 11(27):eadw7697, 2025.

[62] F. P. Mahner, J. Roth, K. C. Lam, M. F. Bonner, F. Pereira, and M. N. Hebart. Characterizing universal object representations across vision models. arXiv preprint arXiv:2605.13675, 2026.

[63] M. Khosla, A. H. Williams, J. McDermott, and N. Kanwisher. Privileged representational axes in biological and artificial neural networks. bioRxiv, 2024.

[64] V. Bosch, R. Sommers, A. Doerig, and T. C. Kietzmann. The umwelt representation hypothesis: Rethinking universality. arXiv preprint arXiv:2604.17960, 2026.

[65] A. Soni, S. Srivastava, M. R. Maechler, K. P. Kording, and M. Khosla. Conclusions drawn from neural network to brain alignment depend strongly on the chosen similarity measure. bioRxiv, 2026.

[66] J. Wu, S. Saha, Y. Bo, and M. Khosla. Comparing and integrating different notions of representational correspondence in neural systems. arXiv preprint arXiv:2509.21628, 2025.

[67] G. Gaziv, M. J. Lee, and J. J. DiCarlo. Strong and precise modulation of human percepts via robustified ANNs. In Advances in Neural Information Processing Systems, volume 36, 2023.

[68] R. Geirhos, J.-H. Jacobsen, C. Michaelis, R. Zemel, W. Brendel, M. Bethge, and F. A. Wichmann. Shortcut learning in deep neural networks. Nature Machine Intelligence, 2(11):665–673, 2020.

[69] J. Roth and M. N. Hebart. How to sample the world for understanding the visual system. In 8th Annual Conference on Cognitive Computational Neuroscience, 2025.

[70] S. Madan, W. Xiao, M. Cao, H. Pfister, M. Livingstone, and G. Kreiman. Benchmarking out-of-distribution generalization capabilities of DNN-based encoding models for the ventral visual cortex. In Advances in Neural Information Processing Systems, volume 37, pages 89249–89277, 2024.

[71] L. Chang and D. Y. Tsao. The code for facial identity in the primate brain. Cell, 169(6):1013–1028.e14, 2017.

[72] B. Wang and C. R. Ponce. On the level sets and invariance of neural tuning landscapes. In Proceedings of the 1st NeurIPS Workshop on Symmetry and Geometry in Neural Representations, volume 197 of Proceedings of Machine Learning Research, pages 278–300, 2023.

[73] E. J. Allen, G. St-Yves, Y. Wu, J. L. Breedlove, J. S. Prince, L. T. Dowdle, M. Nau, B. Caron, F. Pestilli, I. Charest, J. B. Hutchinson, T. Naselaris, and K. Kay. A massive 7t fmri dataset to bridge cognitive neuroscience and artificial intelligence. Nature Neuroscience, 25(1):116–126, 2022.

[74] M. N. Hebart, O. Contier, L. Teichmann, A. H. Rockter, C. Y. Zheng, A. Kidder, A. Corriveau, M. Vaziri-Pashkam, and C. I. Baker. THINGS-data, a multimodal collection of large-scale datasets for investigating object representations in human brain and behavior. eLife, 12:e82580, 2023.

[75] M. St-Laurent, B. Pinsard, O. Contier, E. DuPre, K. Seeliger, V. Borghesani, J. A. Boyle, L. Bellec, and M. N. Hebart. CNeuroMod-THINGS, a densely-sampled fMRI dataset for visual neuroscience. Scientific Data, 13:141, 2026.

[76] J. Zerbe, J. Roth, M. M. Mell, P. Herholz, T. Knapen, and M. N. Hebart. LAION-fMRI: A densely sampled 7T-fMRI dataset providing broad coverage of natural image diversity. In Vision Sciences Society Annual Meeting, 2026.

[77] Y. Li, X. Liu, W. Li, J. Yang, B. Gong, W. Jin, Z. Gong, K. Wang, J. Luo, Z. Zhao, and P. Bao. Triple-N dataset: large-scale fMRI-guided dense recordings of nonhuman primate neural responses to natural scenes. Nature Neuroscience, 29(8):1999–2011, 2026.

[78] N. McNeal and N. A. Ratan Murty. Targeted perturbations reveal brain-like local coding axes in robustified, but not standard, ANN-based brain models. arXiv preprint arXiv:2509.23333, 2025.

[79] D. Skrill, J. Feather, and S. V. Norman-Haignere. Neural prediction decorrelation reveals that adversarial robustness substantially improves DNN prediction accuracy across the entire human auditory cortex. bioRxiv, 2026.

[80] D. Yin, R. Gontijo Lopes, J. Shlens, E. D. Cubuk, and J. Gilmer. A Fourier perspective on model robustness in computer vision. In Advances in Neural Information Processing Systems, volume 32, 2019.

[81] T. Garity, T. Fel, G. A. Alvarez, and T. Serre. The role of frequency in shaping features from artificial vision models. In Conference on Cognitive Computational Neuroscience, 2024.

[82] N. C. L. Kong, E. Margalit, J. L. Gardner, and A. M. Norcia. Increasing neural network robustness improves match to macaque V1 eigenspectrum, spatial frequency preference and predictivity. PLoS Computational Biology, 18(1):e1009739, 2022.

[83] R. Ganz, B. Kawar, and M. Elad. Do perceptually aligned gradients imply robustness? In Proceedings of the 40th International Conference on Machine Learning, volume 202 of Proceedings of Machine Learning Research, pages 10628–10648, 2023.

[84] S. Srinivas, S. Bordt, and H. Lakkaraju. Which models have perceptually-aligned gradients? an explanation via off-manifold robustness. In Advances in Neural Information Processing Systems, volume 36, pages 21172–21195, 2023.

[85] H. Jang and F. Tong. Improved modeling of human vision by incorporating robustness to blur in convolutional neural networks. Nature Communications, 15(1):1989, 2024.

[86] J. Dapello, T. Marques, M. Schrimpf, F. Geiger, D. D. Cox, and J. J. DiCarlo. Simulating a primary visual cortex at the front of CNNs improves robustness to image perturbations. In Advances in Neural Information Processing Systems, volume 33, 2020.

[87] A. Nayebi and S. Ganguli. Biologically inspired protection of deep networks from adversarial attacks. arXiv preprint arXiv:1703.09202, 2017.

[88] T. Konkle and G. Alvarez. Cognitive steering in deep neural networks via long-range modulatory feedback connections. In Advances in Neural Information Processing Systems, volume 36, 2023.

[89] E. U. R. Mohammed, E. Bagheri, Soniya, A. Narayan, and Y. Mohsenzadeh. Integrating non-classical receptive fields of the primary visual cortex into CNNs enhances adversarial robustness. In Conference on Cognitive Computational Neuroscience, Amsterdam, The Netherlands, 2025.

[90] Y. Luo and N. Müller. Viewpoint diversity improves convolutional neural network generalization and robustness. In 8th Annual Conference on Cognitive Computational Neuroscience, Amsterdam, The Netherlands, 2025.

[91] Z. Lu, S. Thorat, R. M. Cichy, and T. C. Kietzmann. Adopting a human developmental visual diet yields robust and shape-based AI vision. Nature Machine Intelligence, 8(5):735–748, 2026.

[92] Z. Shao, T. Ren, C. Wang, L. Isik, and D. M. Beck. Dissociating spatial frequency reliance from adversarial robustness advantages in neurally guided deep convolutional neural networks. arXiv preprint arXiv:2605.04443, 2026.

[93] A. Salvatore, S. Fort, and S. Ganguli. Solving adversarial examples requires solving exponential misalignment. arXiv preprint arXiv:2603.03507, 2026.

[94] J. Raugel, M. Seitzer, M. Szafraniec, H. V. Vo, J. Rapin, P. Labatut, P. Bojanowski, V. Wyart, and J.-R. King. Misalignment between backpropagation and the hierarchy of brain responses to images. arXiv preprint arXiv:2605.28693, 2026.

[95] J. W. Andrade and T. Konkle. Isolating sparse, category-computing circuits in deep neural networks. In Conference on Cognitive Computational Neuroscience, 2025.

[96] S. Black, L. Sharkey, L. Grinsztajn, E. Winsor, D. Braun, J. Merizian, K. Parker, C. R. Guevara, B. Millidge, G. Alfour, and C. Leahy. Interpreting neural networks through the polytope lens. arXiv preprint arXiv:2211.12312, 2022.

[97] D. Wurgaft, C. Rager, M. Kowal, V. Shyam, S. Feucht, U. Bhalla, T. Haklay, E. Bigelow, R. Sarfati, T. McGrath, O. Lewis, J. Merullo, N. Goodman, T. Fel, A. Geiger, and E. S. Lubana. Manifold steering reveals the shared geometry of neural network representation and behavior. arXiv preprint arXiv:2605.05115, 2026.

[98] T. Fel, M. Kowal, M. Jacobs, D. Hazra, U. Bhalla, L. Sharkey, L. Bushnaq, S. Grant, T. Haklay, T. Icard, C. Rager, M. Pearce, D. Wurgaft, A. Swann, F. Doshi, S. Boppana, C. Tigges, N. Cammarata, T. Serre, V. Shyam, O. Lewis, T. McGrath, J. Merullo, E. S. Lubana, and A. Geiger. Structuring sparsity: Block-sparse featurizers capture visual concept manifolds. arXiv preprint arXiv:2606.25234, 2026.

[99] S. Liu, H. Issa, A. Longon, L. Gorton, M. Khosla, A. Williams, and D. Klindt. Similarity of neural network representations in superposition. arXiv preprint arXiv:2604.00208, 2026.

[100] A. Longon, D. Klindt, and M. Khosla. Superposition disentanglement of neural representations reveals hidden alignment. arXiv preprint arXiv:2510.03186, 2025.

[101] J. J. Jun, N. A. Steinmetz, J. H. Siegle, D. J. Denman, M. Bauza, B. Barbarits, A. K. Lee, C. A. Anastassiou, A. Andrei, Ç. Aydın, M. Barbic, T. J. Blanche, V. Bonin, J. Couto, B. Dutta, S. L. Gratiy, D. A. Gutnisky, M. Häusser, B. Karsh, P. Ledochowitsch, C. Mora Lopez, C. Mitelut, S. Musa, M. Okun, M. Pachitariu, J. Putzeys, P. D. Rich, C. Rossant, W.-l. Sun, K. Svoboda, M. Carandini, K. D. Harris, C. Koch, J. O’Keefe, and T. D. Harris. Fully integrated silicon probes for high-density recording of neural activity. Nature, 551(7679):232–236, 2017.

[102] J. Hwang, A. R. Mitz, and E. A. Murray. NIMH MonkeyLogic: Behavioral control and data acquisition in MATLAB. Journal of Neuroscience Methods, 323:13–21, 2019.

[103] H. Supèr and P. R. Roelfsema. Chronic multiunit recordings in behaving animals: advantages and limitations. Progress in Brain Research, 147:263–282, 2005.

[104] X. Chen, F. Wang, E. Fernandez, and P. R. Roelfsema. Shape perception via a high-channel-count neuroprosthesis in monkey visual cortex. Science, 370(6521):1191–1196, 2020.

[105] A. D. Legatt, J. Arezzo, and H. G. Vaughan, Jr. Averaged multiple unit activity as an estimate of phasic changes in local neuronal activity: effects of volume-conducted potentials. Journal of Neuroscience Methods, 2(2):203–217, 1980.

[106] R. Bauer, M. Brosch, and R. Eckhorn. Different rules of spatial summation from beyond the receptive field for spike rates and oscillation amplitudes in cat visual cortex. Brain Research, 669(2):291–297, 1995.

[107] A. Stigliani, K. S. Weiner, and K. Grill-Spector. Temporal processing capacity in high-level visual cortex is domain specific. Journal of Neuroscience, 35(36):12412–12424, 2015.

[108] A. Krizhevsky, I. Sutskever, and G. E. Hinton. Imagenet classification with deep convolutional neural networks. In Advances in Neural Information Processing Systems, volume 25, 2012.

[109] J. Mehrer, C. J. Spoerer, N. Kriegeskorte, and T. C. Kietzmann. Individual differences among deep neural network models. Nature Communications, 11(1):5725, 2020.

[110] K. He, X. Zhang, S. Ren, and J. Sun. Deep residual learning for image recognition. In Proceedings of the IEEE Conference on Computer Vision and Pattern Recognition (CVPR), pages 770–778, 2016.

[111] H. Salman, A. Ilyas, L. Engstrom, A. Kapoor, and A. Mądry. Do adversarially robust imagenet models transfer better? In Advances in Neural Information Processing Systems, volume 33, 2020.

[112] A. Radford, J. W. Kim, C. Hallacy, A. Ramesh, G. Goh, S. Agarwal, G. Sastry, A. Askell, P. Mishkin, J. Clark, G. Krueger, and I. Sutskever. Learning transferable visual models from natural language supervision. In Proceedings of the 38th International Conference on Machine Learning, volume 139 of Proceedings of Machine Learning Research, pages 8748–8763. PMLR, 2021.

[113] M. Caron, H. Touvron, I. Misra, H. Jégou, J. Mairal, P. Bojanowski, and A. Joulin. Emerging properties in self-supervised vision transformers. In Proceedings of the IEEE/CVF International Conference on Computer Vision (ICCV), pages 9650–9660, 2021.

[114] I. Radosavovic, R. P. Kosaraju, R. Girshick, K. He, and P. Dollár. Designing network design spaces. In Proceedings of the IEEE/CVF Conference on Computer Vision and Pattern Recognition (CVPR), pages 10425–10433, 2020.

[115] P. Goyal, M. Caron, B. Lefaudeux, M. Xu, P. Wang, V. Pai, M. Singh, V. Liptchinsky, I. Misra, A. Joulin, and P. Bojanowski. Self-supervised pretraining of visual features in the wild. arXiv preprint arXiv:2103.01988, 2021.

[116] R. Ganz and M. Elad. CLIPAG: Towards generator-free text-to-image generation. In Proceedings of the IEEE/CVF Winter Conference on Applications of Computer Vision (WACV), pages 3843–3853, 2024.

[117] M. Tschannen, A. Gritsenko, X. Wang, M. F. Naeem, I. Alabdulmohsin, N. Parthasarathy, T. Evans, L. Beyer, Y. Xia, B. Mustafa, O. Hénaff, J. Harmsen, A. Steiner, and X. Zhai. SigLIP 2: Multilingual vision-language encoders with improved semantic understanding, localization, and dense features. arXiv preprint arXiv:2502.14786, 2025.

[118] M. Oquab, T. Darcet, T. Moutakanni, H. Vo, M. Szafraniec, V. Khalidov, P. Fernandez, D. Haziza, F. Massa, A. El-Nouby, M. Assran, N. Ballas, W. Galuba, R. Howes, P.-Y. Huang, S.-W. Li, I. Misra, M. Rabbat, V. Sharma, G. Synnaeve, H. Xu, H. Jégou, J. Mairal, P. Labatut, A. Joulin, and P. Bojanowski. DINOv2: Learning robust visual features without supervision. Transactions on Machine Learning Research, 2024.

[119] T. Darcet, M. Oquab, J. Mairal, and P. Bojanowski. Vision transformers need registers. In International Conference on Learning Representations (ICLR), 2024.

[120] M. Ranzinger, G. Heinrich, J. Kautz, and P. Molchanov. AM-RADIO: Agglomerative vision foundation model – reduce all domains into one. In Proceedings of the IEEE/CVF Conference on Computer Vision and Pattern Recognition (CVPR), pages 12490–12500, 2024.

[121] G. Heinrich, M. Ranzinger, H. Yin, Y. Lu, J. Kautz, A. Tao, B. Catanzaro, and P. Molchanov. RADIOv2.5: Improved baselines for agglomerative vision foundation models. In Proceedings of the IEEE/CVF Conference on Computer Vision and Pattern Recognition (CVPR), pages 22487–22497, 2025.

[122] F. Pedregosa, G. Varoquaux, A. Gramfort, V. Michel, B. Thirion, O. Grisel, M. Blondel, P. Prettenhofer, R. Weiss, V. Dubourg, J. Vanderplas, A. Passos, D. Cournapeau, M. Brucher, M. Perrot, and É. Duchesnay. Scikit-learn: machine learning in python. Journal of Machine Learning Research, 12:2825–2830, 2011.

[123] A. Nguyen, J. Yosinski, and J. Clune. Deep neural networks are easily fooled: High confidence predictions for unrecognizable images. In Proceedings of the IEEE Conference on Computer Vision and Pattern Recognition, pages 427–436, 2015.

[124] J. Yosinski, J. Clune, A. Nguyen, T. Fuchs, and H. Lipson. Understanding neural networks through deep visualization. In ICML Deep Learning Workshop, 2015. arXiv:1506.06579.

[125] C. Olah, A. Mordvintsev, and L. Schubert. Feature visualization. Distill, 2(11):e7, 2017.

[126] A. H. Gray Jr and J. D. Markel. A spectral-flatness measure for studying the autocorrelation method of linear prediction of speech analysis. IEEE Transactions on Acoustics, Speech, and Signal Processing, 22(3):207–217, 1974.

[127] J. D. Johnston. Transform coding of audio signals using perceptual noise criteria. IEEE Journal on Selected Areas in Communications, 6(2):314–323, 1988.

[128] L. Liu, B. Liu, H. Huang, and A. C. Bovik. No-reference image quality assessment based on spatial and spectral entropies. Signal Processing: Image Communication, 29(8):856–863, 2014.

[129] O. Russakovsky, J. Deng, H. Su, J. Krause, S. Satheesh, S. Ma, Z. Huang, A. Karpathy, A. Khosla, M. Bernstein, A. C. Berg, and L. Fei-Fei. ImageNet large scale visual recognition challenge. International Journal of Computer Vision, 115(3):211–252, 2015.

[130] T. Konkle, T. F. Brady, G. A. Alvarez, and A. Oliva. Conceptual distinctiveness supports detailed visual long-term memory for real-world objects. Journal of Experimental Psychology: General, 139(3):558–578, 2010.

[131] K. Vinken, J. S. Prince, T. Konkle, and M. S. Livingstone. The neural code for “face cells” is not face-specific. Science Advances, 9(35):eadg1736, 2023.

