## Supplementary Materials for "Parametric neural control differentiates top neural network models of primate visual cortex"

### Contents: Supplementary Figures


*Experimental framework and calibration*

**S1** Calibration image set . . . 4
**S2** Response reliability and NSD noise ceiling . . . 5
**S3** Array and Neuropixels placement . . . 7
**S4** Site category-selectivity . . . 8
**S5** Per-channel encoding-layer selection . . . 9

*Feature accentuation*

**S6** Automated synthesis-regularization selection . . . 10
**S7** Feature-accentuation success rate . . . 10
**S8** Structure of synthesis failures . . . 11
**S9** Encoding-axis alignment of accentuation sweeps . . . 11
**S10** Accentuation sweeps in the read-out-aligned embedding . . . 12
**S11** Suppress and drive extremes across sites . . . 13
**S12–16** Accentuation sweep galleries . . . 14

*Parametric neural control*

**S17** Model differences under the control-slope measure . . . 19
**S18** Control outcomes under alternative scoring metrics and per-seed aggregation . . . 19
**S19** Controllability variation across sites . . . 20
**S20** Per-site control score (all 10 models) . . . 20
**S21** Cross-area consistency of model rankings . . . 21
**S22–26** High-control example sweeps . . . 22
**S27** Super-stimulus proportion . . . 27
**S28** Super-stimulus response scatter, Monkey V . . . 27
**S29** Super-stimulus response scatter, all sites . . . 28
**S30** Neural control across model inductive bias groups . . . 29
**S31** Reliability of control-phase responses . . . 30
**S32** Synthesis regularization and parametric control . . . 31
**S33** Control outcomes after regressing out the effect of synthesis hyperparameters . . . 32
**S34** Relationship between axis alignment and neural control outcomes . . . 32
**S35** Control outcomes across accentuated-image exclusion regimes . . . 33

*Controversial accentuations*

**S36** Controversial-stimulus experiment . . . 34
**S37** Controversial-accentuation stimuli . . . 35
**S38** Controversial-accentuation results . . . 36

*Adversarial sensitivity and input-gradient structure*

**S39** Encoding-axis adversarial sensitivity and neural control . . . 37
**S40** Adversarial attack-method comparisons . . . 38
**S41** Alternative metrics for estimating adversarial sensitivity and gradient spectral concentration . . . 39
**S42** Computing gradient spectral participation ratio (PR) . . . 40
**S43** Input-gradient map galleries by seed image . . . 42
**S44** Encoding-gradient geometry and neural control . . . 43

**S45** Robustness of gradient spectral participation ratio to probe images and analysis choices . . . 44
**S46** Gradient spectra and their prediction of control across encoding training diets . . . 45
**S47** Gradient spectral participation ratio and control within matched regularization regimes . . . 46
**S48** Cross-validated prediction of parametric control among the trained models . . . 47
**S49** Predicting neural control: single-predictor leaderboard and cross-validated best model . . . 48

### Experimental framework and calibration

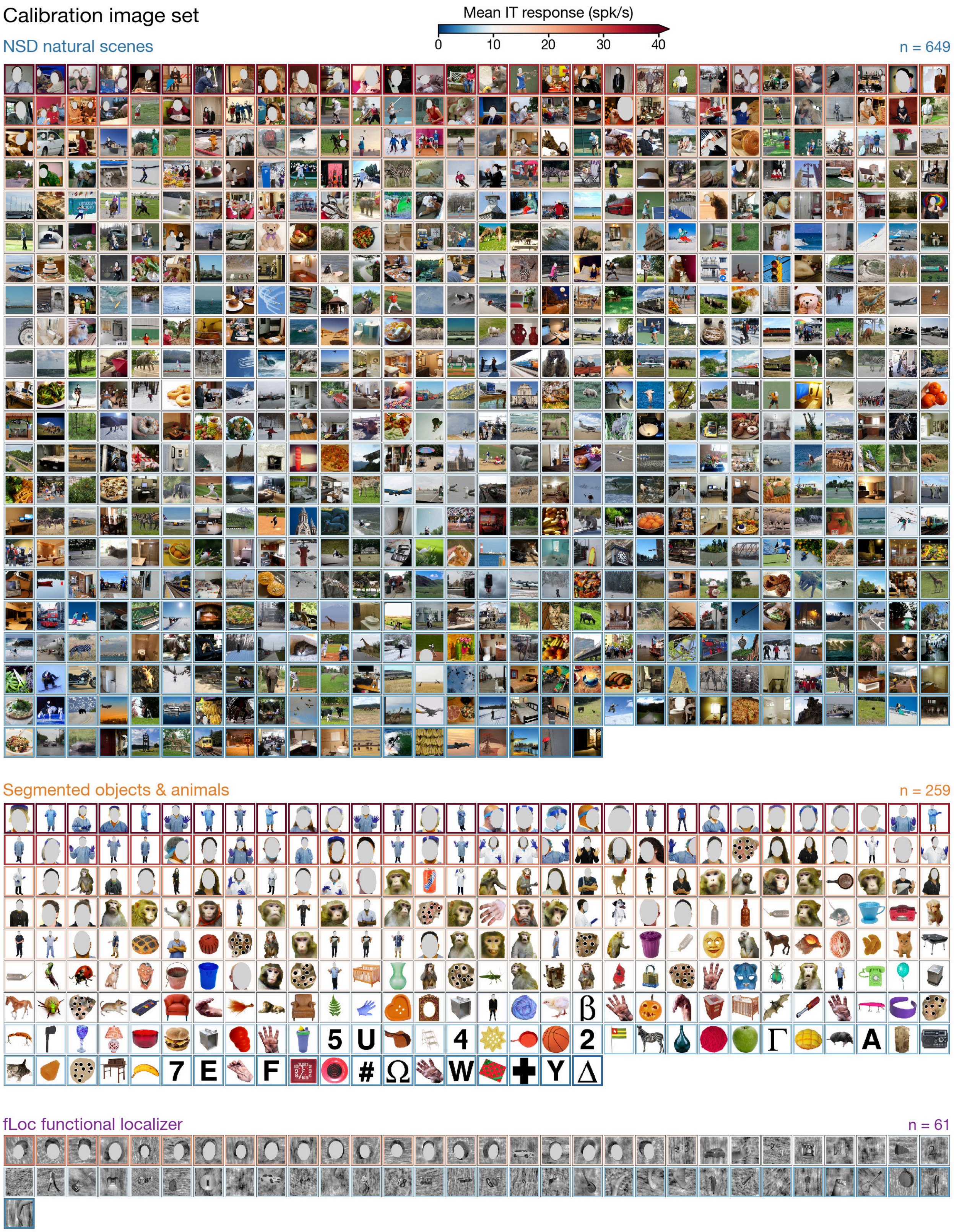

***Supplementary Figure 1. Calibration image set.*** *The full mosaic of the* 969 *natural and synthetic images used to fit the encoding models is shown, grouped by the source of the images. Within each source, images are sorted by mean IT response (high to low), with borders coding the corresponding firing rates averaged across the five aIT electrodes from Monkey R (blue = low, white = median, red = high spk/s). The three sources are NSD natural scenes* ($n = 649$)*, segmented objects and animals on white backgrounds* ($n = 259$)*, and fLoc functional-localizer images* ($n = 61$)*. The strongest responses were driven by images of human faces (see the warm-bordered thumbnails in the segmented-object and fLoc blocks).*

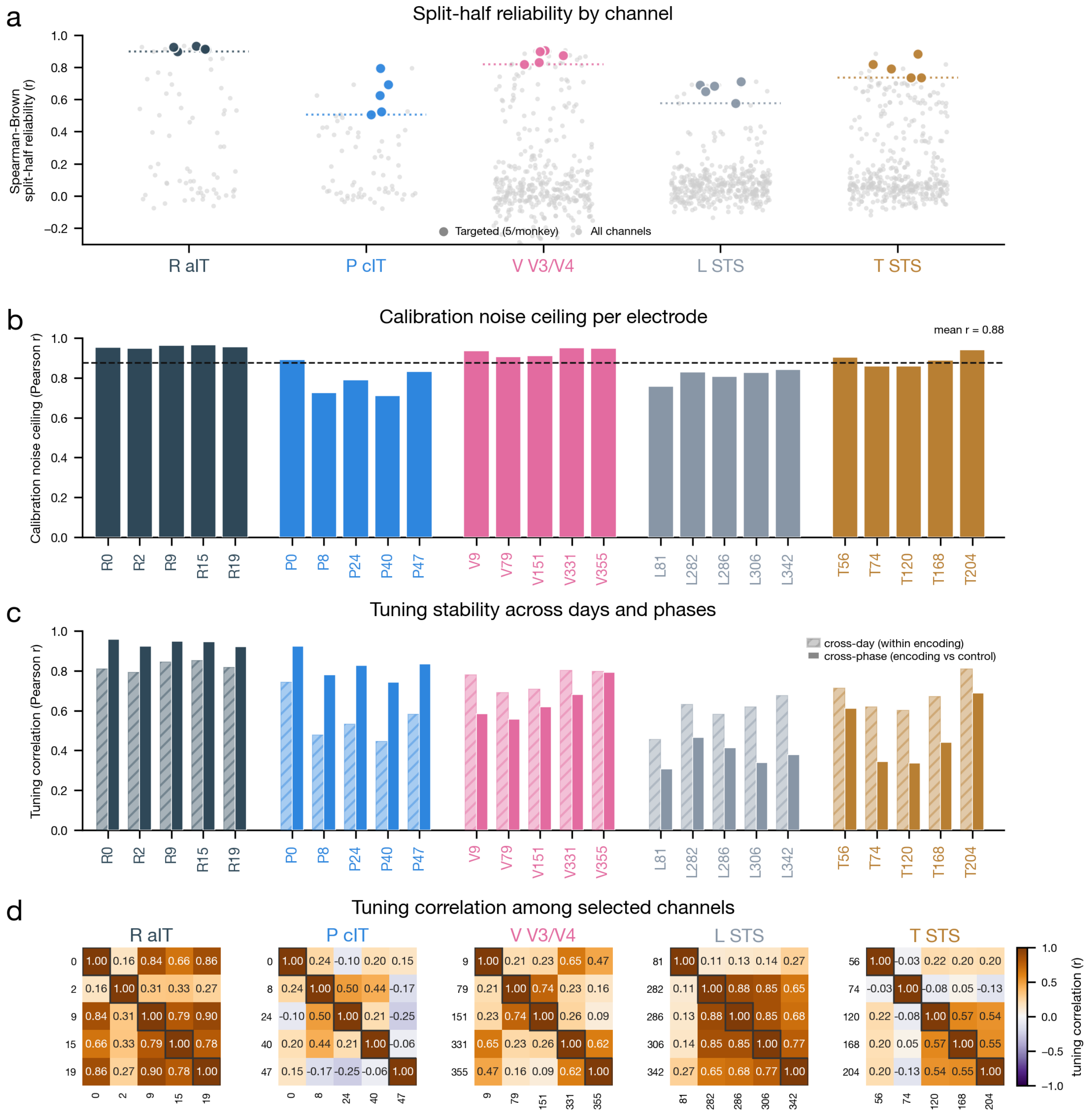


***Supplementary Figure 2. Response reliability and NSD noise ceiling.*** *Characterization of how the 25 targeted electrodes were selected and how reliably they respond, across the 5 macaques (Monkey R aIT, Monkey P cIT, Monkey V V3/V4, Monkey L STS, Monkey T STS). (A) Split-half response reliability by channel. For each animal, we plot the Spearman-Brown-corrected split-half reliability* ($r$; $2r/(1+r)$ *over repeats occurring during the calibration phase) for every array/probe channel (gray). The 5 targeted sites are highlighted per animal and a dotted line marks the minimum*

*reliability of the 5 selected sites. (B) Calibration noise ceiling per electrode. Bars show the per-site ceiling (Pearson* $r$*), the maximum model-data correlation for the trial-averaged calibration response, computed from the same calibration data as (A) as* $\sqrt{r_{\text{split-half}}}$ *(the NSD* $\mathrm{nc\_r}$ *identity; Allen et al., 2022), for each of the 25 sites, grouped and colored by animal; all five animals have a valid ceiling; control-phase noise ceilings, used for the control analyses, are reported separately. Dashed line: mean* $r = 0.88$ *across all 25 sites. (C) Tuning stability measured per site: cross-day (within the calibration phase, dashed fill) and cross-phase (calibration vs control phases, solid fill) Pearson* $r$ *of responses over the shared calibration images, grouped by animal. (D) The tuning correlations among the 5 targeted channels per animal:* $5 \times 5$ *correlation matrices (Pearson* $r$*) over calibration-phase images, confirming that selected channels are reliable and have relatively diverse tuning profiles.*

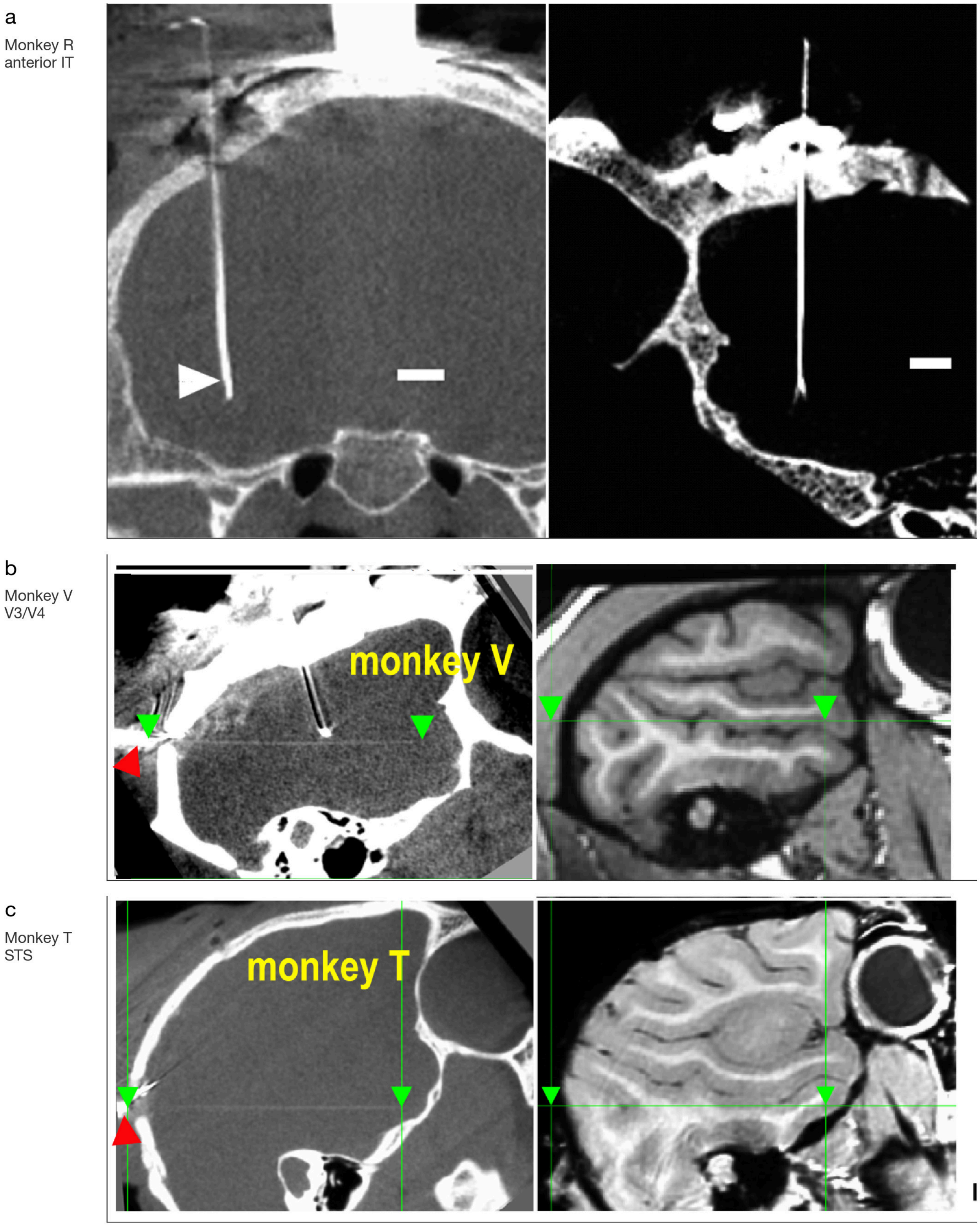


***Supplementary Figure 3. Array and Neuropixels probe placement across recorded animals.*** *Intra-operative CT and structural MRI sections showing probe tracks for the recorded sites. (A) Monkey R, anterior IT: two CT sections; the white arrowhead marks the chronic microwire array tip. (B) Monkey V, V3/V4: CT section (left) and structural MRI (right); green arrowheads mark Neuropixels probe entry and target. (C) Monkey T, STS: same format as (B).*

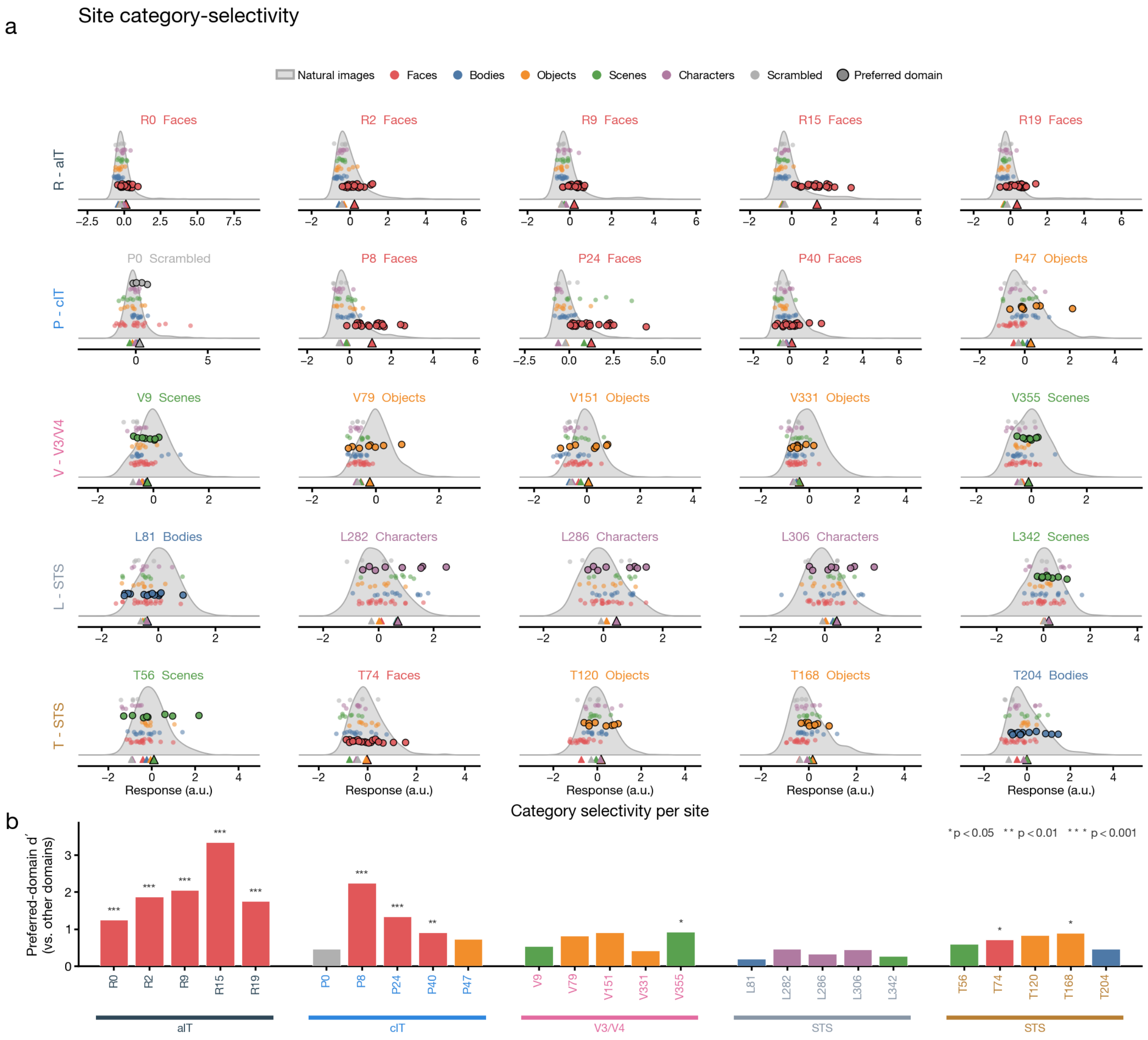


***Supplementary Figure 4. Category selectivity of recorded sites.*** *fLoc category responses of the $n = 25$ recorded sites, shown against each site's calibration-phase natural-image response distribution. (A) In-context response distributions, one panel per site, grouped by animal and area (Monkey R/aIT, P/cIT, V/V3/V4, L/STS, T/STS; 5 sites each). Gray shading is the kernel density of natural-image responses; jittered dots are the six fLoc category responses (Faces, Bodies, Objects, Scenes, Characters, Scrambled), with triangles marking category means. Each site's preferred domain (largest mean) is highlighted with black-edged markers and named in the panel title. (B) Summary of preferred-domain selectivity across all 25 sites. Bar height is the preferred-domain $d'$ (preferred category versus all other categories), colored by preferred domain; $x$-tick labels and region brackets are colored by animal. Asterisks denote a Welch $t$-test of preferred versus other categories ($*\,p < 0.05$, $*\,*\;p < 0.01$, $*\,*\,*\,p < 0.001$). aIT and cIT sites, targeted to face patches, show the strongest and most consistent selectivity, whereas V3/V4 and STS sites are more mixed.*

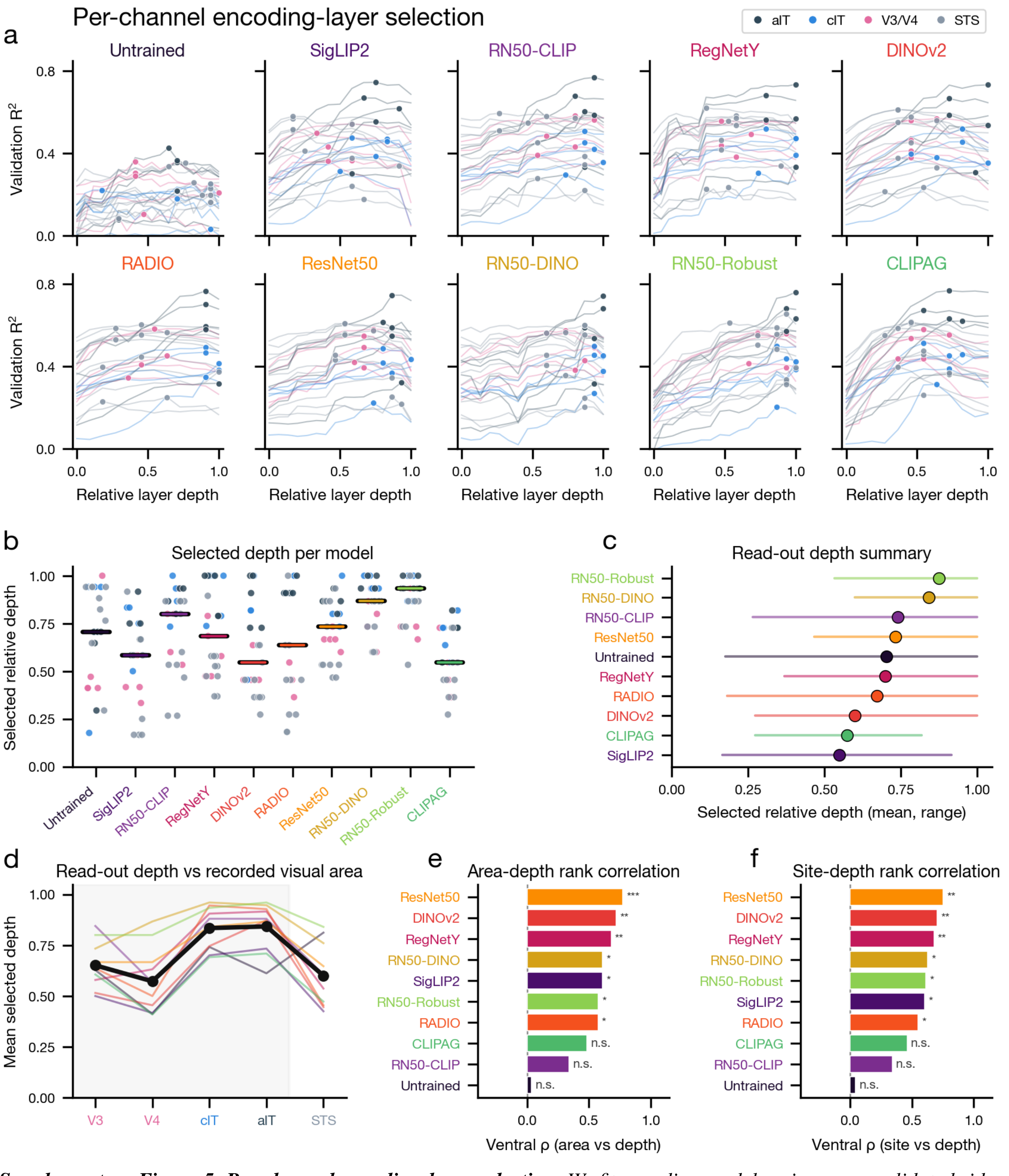


***Supplementary Figure 5. Per-channel encoding-layer selection.*** *We fit encoding models using cross-validated ridge regression for each of the* 250 *model-site pairs. In each case, the best-predicting backbone layer was chosen per channel via held-out validation* $R^2$*. Site markers are colored by recorded region (aIT, cIT, V3/V4, STS); models comprise* 10 *networks spanning three trained-model families and an Untrained baseline. (A) Validation* $R^2$ *versus relative layer depth for each model (one panel per model, one line per site,* $n = 25$ *sites), with the selected peak block marked. Curves generally rise then plateau, with peaks concentrated at intermediate-to-late depths. (B) Selected relative depth per model (swarm of* $n = 25$ *sites, colored by region; black-over-color bar = per-model median). (C) Read-out depth summary: per-model mean selected relative depth (dot) with min-max range (line), sorted by mean. (D) Mean selected depth per recorded visual area (V3, V4, cIT, aIT, STS), one line per model (colored) plus grand mean (black); ventral*

*areas shaded. Read-out depth is generally greater for more anterior ventral stream areas. (E) Per-model Spearman rank correlation between selected depth and coarse ventral area rank (V3 < V4 < cIT < aIT), sorted; bars colored by model, with significance stars (*** $p < 10^{-3}$, ** $p < 10^{-2}$, * $p < 0.05$, n.s. otherwise). (F) Same as (E), but over a finer ventral-stream ordering, with monkey V's 5 sites resolved by their location along the Neuropixels probe. Adversarially trained models are shown in bold.*

### Feature accentuation

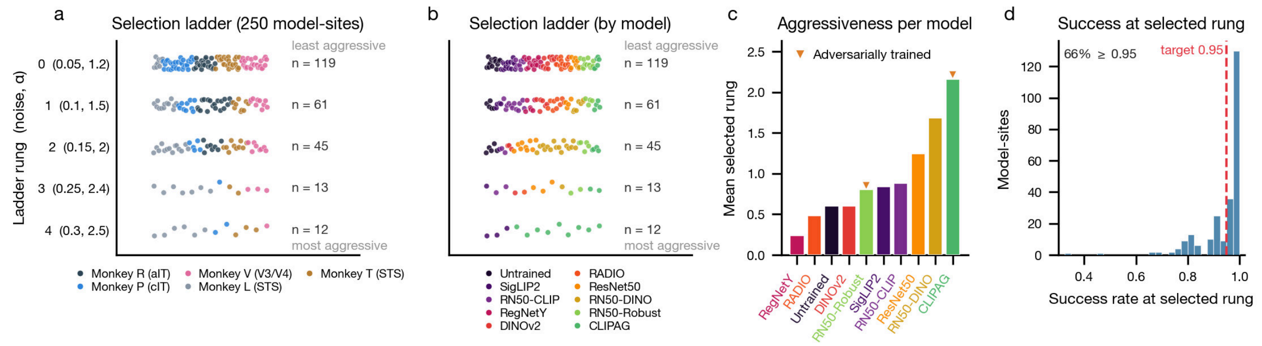


***Supplementary Figure 6. Automated synthesis-regularization selection.*** *An automated procedure selected the synthesis regularization independently for each of the 250 model-site combinations, stepping down a five-rung ladder of (augmentation-noise, spectral-decay $\alpha$) regimes ordered most to least aggressive and stopping at the first rung whose synthesis success rate exceeded* 0.95*. (A) The selected rung for all $n = 250$ model-sites (25 sites $\times$ 10 models), dodged within each rung and colored by animal (R/aIT, P/cIT, V/V3/V4, L and T/STS); rung labels give the (noise, $\alpha$) pair, with rung* 0 *least aggressive. Most sites land on the two least-aggressive rungs ($n = 119$ and $n = 61$). (B) The same selection ladder recolored by model (legend), showing that every model spreads across the rungs. (C) Per-model mean selected rung (0 = least aggressive), colored by model and ordered ascending; adversarially trained models (orange triangles) are marked. Models differ only modestly in the aggressiveness they require. (D) Distribution of the achieved success rate at the selected rung across the* 250 *model-sites; dashed red line marks the* 0.95 *target. The procedure returns the best-available regime, so* 66% *of sites clear the* 0.95 *line.*

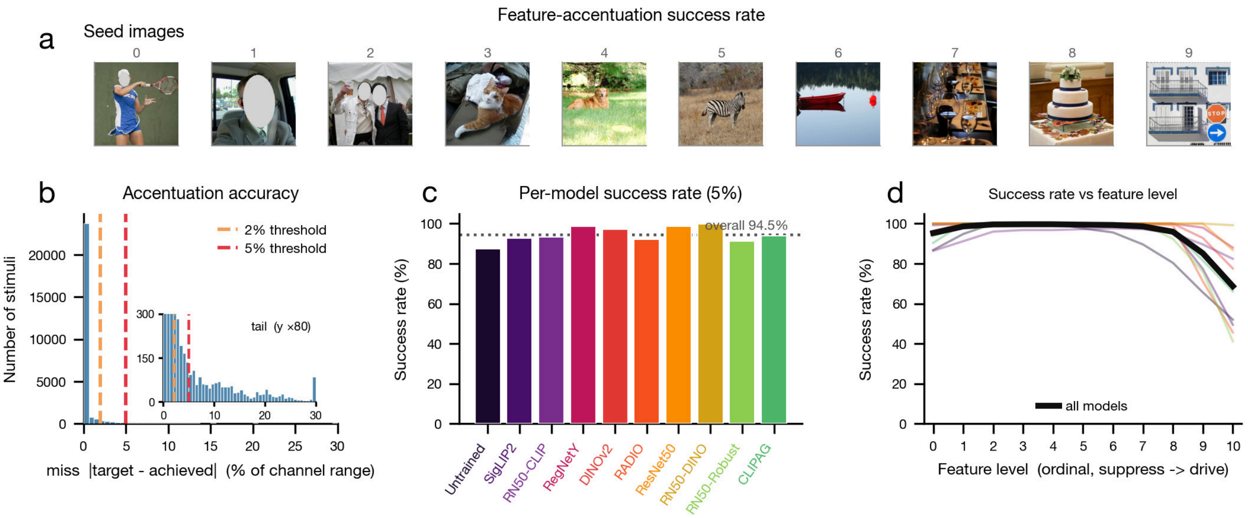


***Supplementary Figure 7. Feature-accentuation success rate.*** *Across all synthesized stimuli, the achieved level (each generating model's post-hoc predicted response for the saved image) is compared to its intended target, and a synthesis is scored successful when the miss (i.e., the relative error), $|\text{target} - \text{achieved}|/\text{range}$, falls within tolerance $\tau$, where* range *is the per-channel target range. Overall success was* 94.5% *at $\tau = 5\%$ and* 90.7% *at $\tau = 2\%$. (A) The ten seed*

*images used in feature accentuation. (B) Histograms are shown of the miss quantities (% of channel target range) pooled over all stimuli, with dashed lines marking the* 2% *(orange) and* 5% *(red) thresholds; note that a majority of the stimuli concentrate near zero. Inset magnifies the $y$-axis ($\times 80$) to expose the tail beyond the main peak. (C) Per-model success rate at $\tau = 5\%$ for the 10 models, colored by model, with the dotted line and label marking the overall rate (94.5%); rates are consistently high across all models. (D) Success rate at $\tau = 5\%$ versus the 11 ordinal drive levels (suppress to drive), one faint trace per model and the bold black trace pooling all models; success stays near ceiling through the mid-range and falls off only at the most extreme drive levels.*

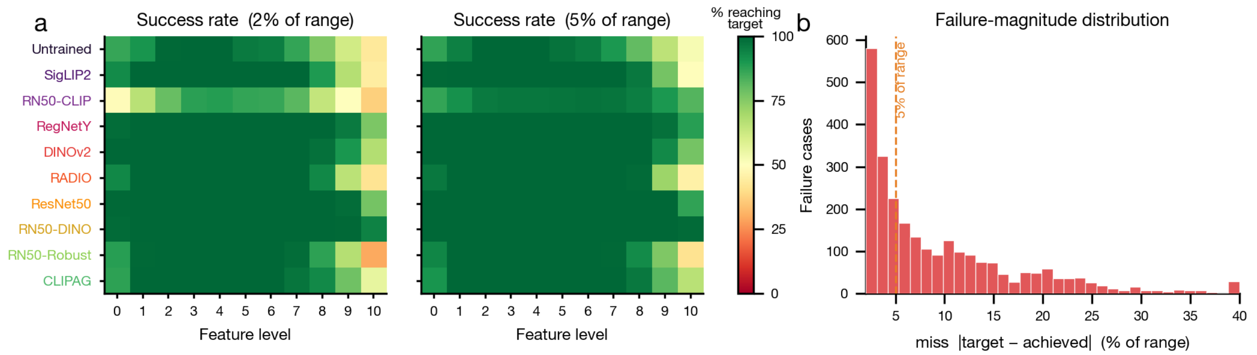


***Supplementary Figure 8. Structure of synthesis failures.*** *A synthesized image "fails" when its own post-hoc prediction misses the intended target by more than a fraction $\tau$ of the channel's target range ($\mathrm{relerr} = |\mathrm{target}-\mathrm{achieved}|/\mathrm{range} > \tau$); the target grid is shared across models within a channel, so this criterion is comparable across models. (A) Success rate ($100\times$ the fraction of stimuli with $\mathrm{relerr} \leq \tau$), shown as model ($y$, 10 models colored by family with adversarially trained models in bold) by feature level ($x$, 11 target levels 0–10) heatmaps (RdYlGn, 0–100%) at the strict ($\tau = 2\%$ of range) and lenient ($\tau = 5\%$) tolerances, pooled over all sites and seeds across the 5 macaques. Failures tend to be non-uniform, instead concentrating at the strongest drive levels and disproportionately arising from specific models (e.g. RN50-CLIP). Relaxing the success criterion threshold to $\tau = 5\%$ mitigates many error cases. (B) Distribution of miss magnitude $|\mathrm{target} - \mathrm{achieved}|$ (% of range) among failure cases only ($\mathrm{relerr} > 2\%$, so the success spike near 0 is excluded to expose the tail); dashed orange line marks the 5% tolerance. Most misses are small, with a long right tail.*

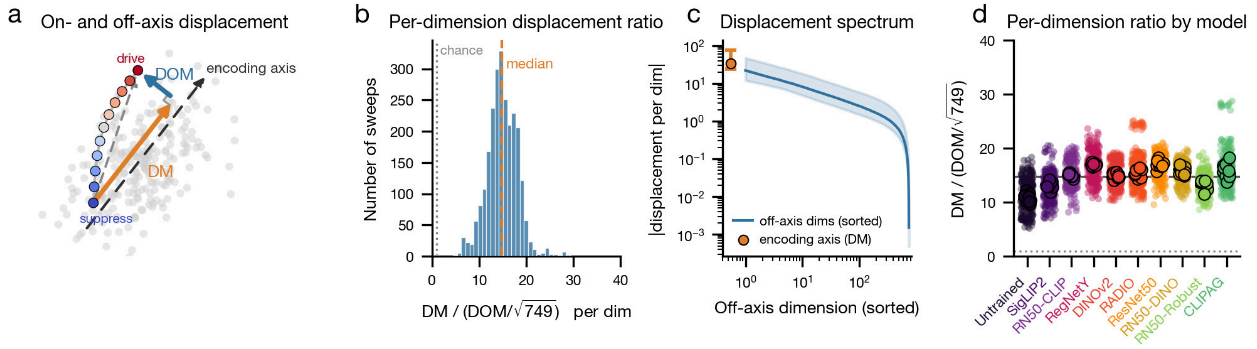


***Supplementary Figure 9. Encoding-axis alignment of accentuation sweeps.*** *Each sweep's net latent displacement $d$ in the 750-dimensional PCA feature space is decomposed into an on-axis component along the fitted encoding axis $\hat{a}$, $\mathrm{DM} = |d \cdot \hat{a}|$, and an off-axis residual over the remaining 749 dimensions, $\mathrm{DOM} = \|d - (d \cdot \hat{a})\,\hat{a}\|$. DM stands for directional modulation and DOM stands for direction-orthogonal modulation. Because DOM pools 749 dimensions, alignment is quantified per representational dimension as $\mathrm{DM}/(\mathrm{DOM}/\sqrt{749})$. (A) We provide a schematic of the on- and off-axis decomposition in a toy-example latent space: an 11-level accentuation sweep (blue "suppress" to red "drive") is split into its DM (orange) and DOM (blue) components. (B) Distributions of the per-dimension ratio across stimulus sweeps (orange dashed median $\approx 15$; gray dotted line at $x = 1$ marks the level where DM equals the root-mean-square (RMS) per-dimension off-axis displacement, $\mathrm{DM} = \mathrm{DOM}/\sqrt{749}$). $n = 2{,}500$ sweeps (25 sites $\times$ 10 models $\times$ 10 seed images). (C) In log-log space we show the distributions (over sweeps) of displacement either along the encoding axis (DM, orange point, median with IQR) or for the sorted off-axis dimensions (blue line, median with shaded IQR band); note that the displacement along the encoding axis is the single most-modulated dimension in 87.2%*

*of sweeps. (D) Per-dimension ratio by model, one column per model (dots jittered, colored by model, adversarially trained models in bold; large dots mark the* 10 *seeds of a representative channel per model). Black-and-color median bars per model; gray dashed line is the overall median and gray dotted line marks the same* $x = 1$ *reference as in (B). Alignment sits uniformly well above this reference across all* 10 *models (including the Untrained baseline) (median* $\approx 15\times$ *greater per-dimension displacement along the encoding axis than the root-mean-square (RMS) off-axis displacement per dimension).*

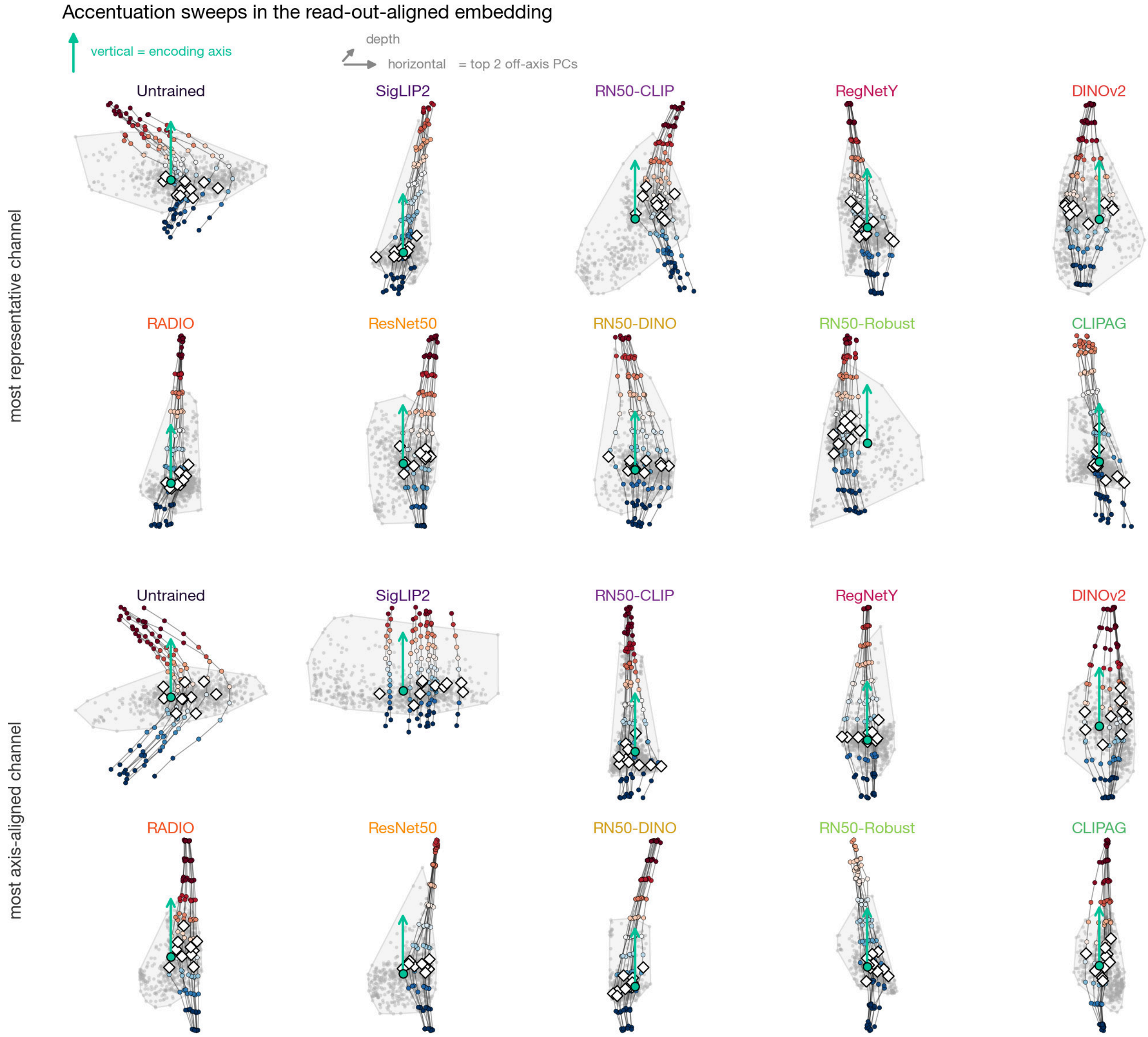


***Supplementary Figure 10. Accentuation sweeps in the read-out-aligned embedding.*** *We plot galleries of feature-accentuation sweeps for a set of example sites, projected into a three-dimensional read-out-aligned embedding: the vertical direction represents the fitted encoding axis (teal arrow), and the horizontal and depth axes represent the top two principal components of the residual* 749*-dimensional space. In every panel, gray dots and the shaded convex hull show the natural calibration images, with colored dots tracing accentuation sweeps derived from all ten seed images (dot color shows predicted firing rate; thin black lines connect levels within a sweep, and white diamonds mark the natural seed images). Top two rows show the most representative channel per model, i.e. the site whose mean on-axis/off-axis per-dimension displacement ratio* $\mathrm{DM}/(\mathrm{DOM}/\sqrt{n_{off}})$ *is closest to that model's mean. The bottom two rows show the most axis-aligned channel per model, i.e. the site with the largest mean* $\mathrm{DM}/(\mathrm{DOM}/\sqrt{n_{off}})$.

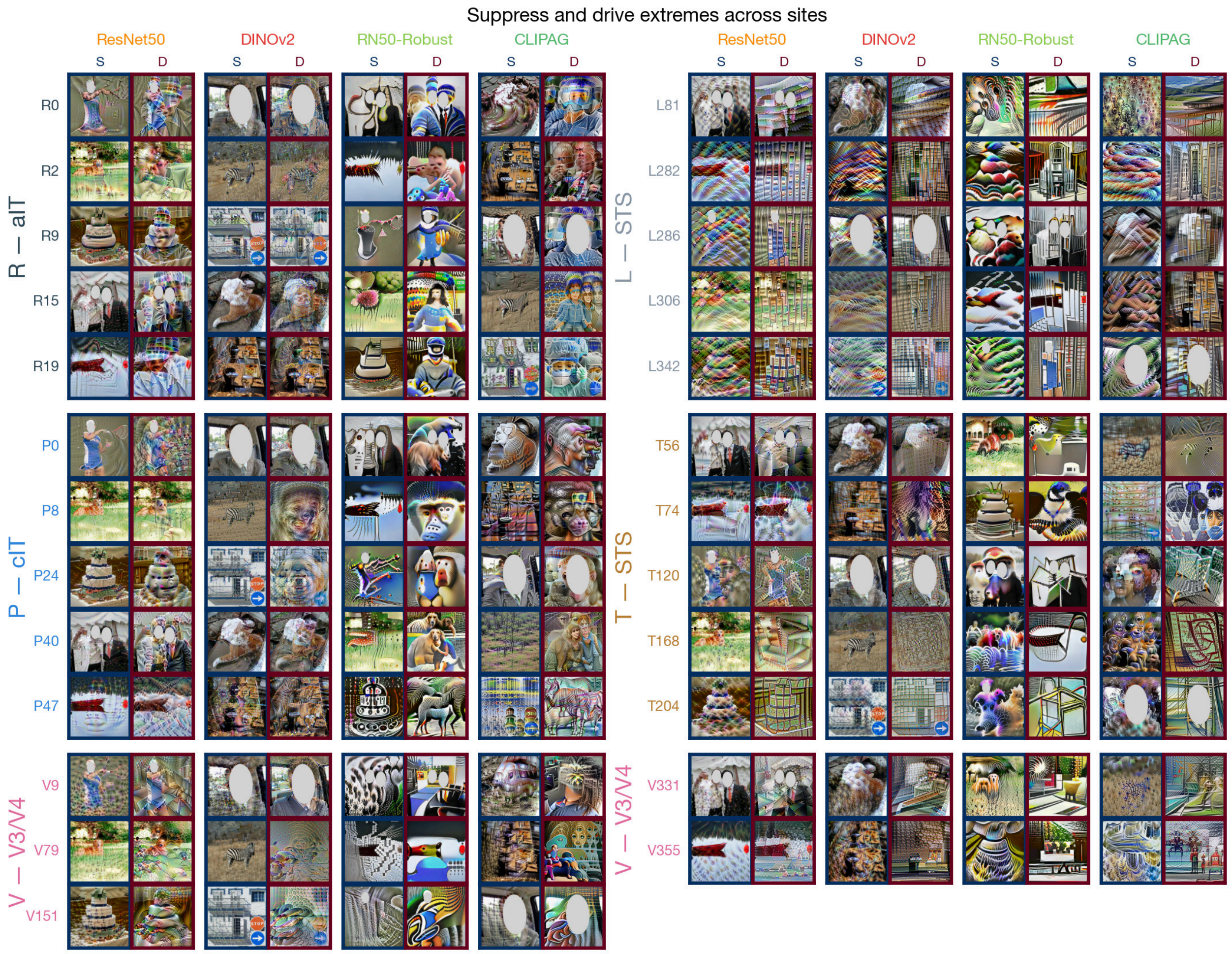


***Supplementary Figure 11. Suppress and drive extremes across sites.*** *We show galleries of the accentuated images predicted to be most strongly suppressing and most strongly driving, generated for each of the 25 recorded sites, across four representative models (ResNet50, DINOv2, RN50-Robust, and CLIPAG). Each cell shows the accentuated stimulus at the lowest predicted level (level 0, S) or highest predicted level (level 10, D) for that (site, model) pair; the seed image rotates across (site, model) pairs to sample the full seed set. Border color encodes predicted response level.*

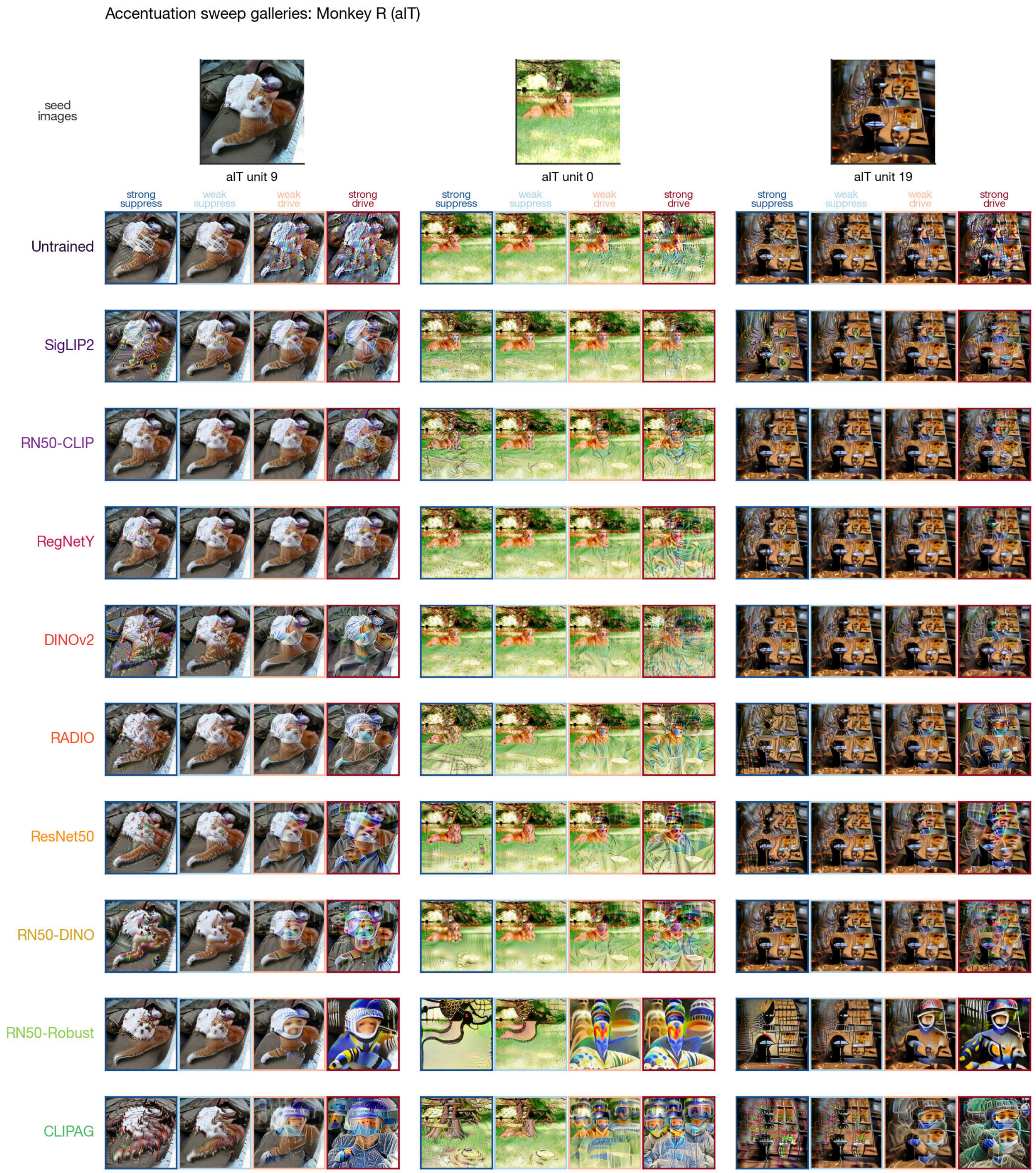


***Supplementary Figure 12. Accentuation sweep galleries – Monkey R (aIT).*** *Here we show, for each monkey, example accentuated stimuli for three recorded sites (one seed image of the ten is chosen for plotting per site). Rows represent the ten models and columns represent four target levels along each model's accentuation axis (strong suppress, weak suppress, weak drive, strong drive). Because the columns walk the same suppress-to-drive sweep for every model, the grid exposes how different models modify the same image to achieve the target response. Images are the raw synthesized accentuation stimuli; the level ordering per (model, site, seed) is taken from that site-model's personalized sweep.*

Accentuation sweep galleries: Monkey P (cIT)

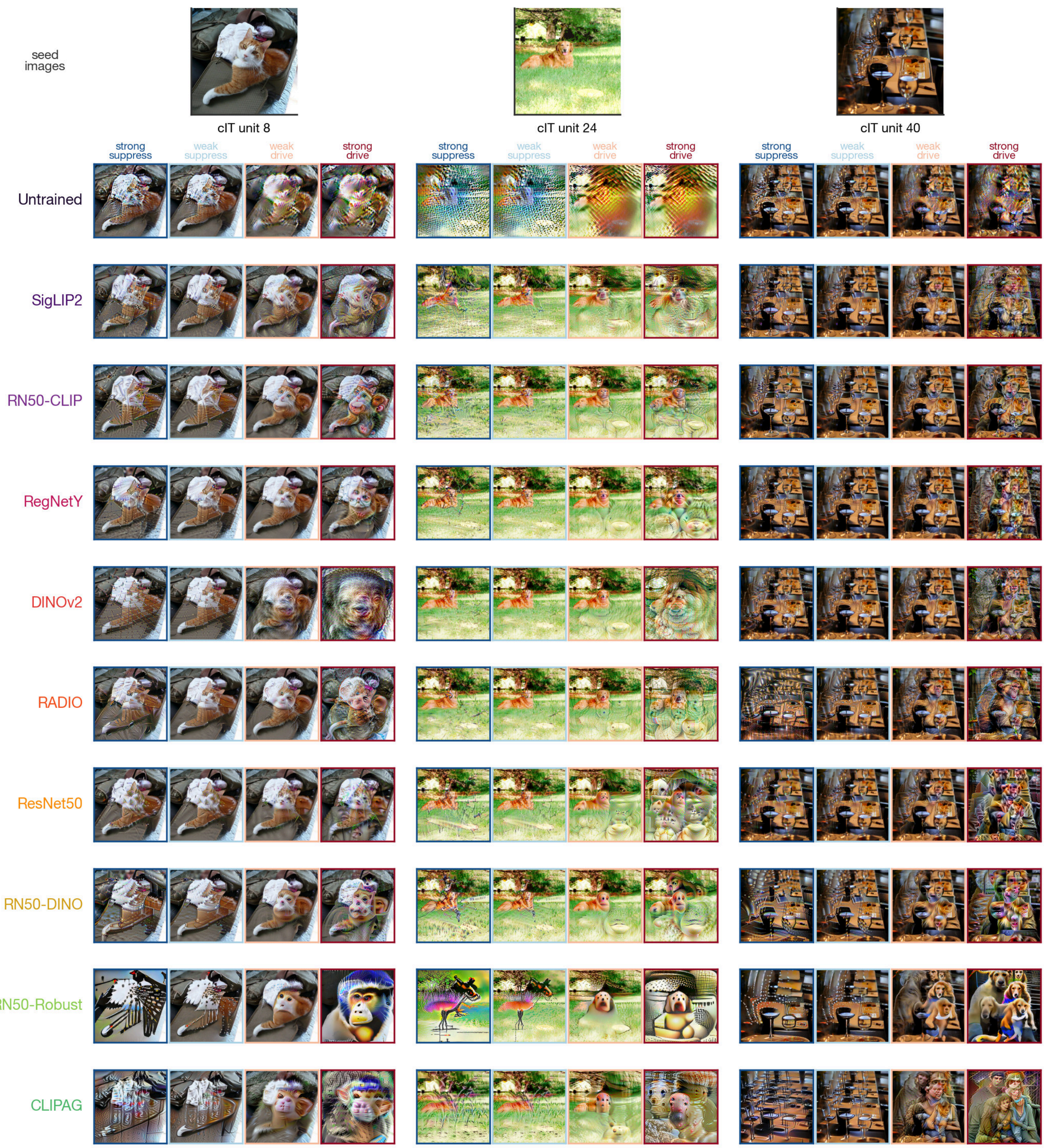


***Supplementary Figure 13. Accentuation sweep galleries – Monkey P (cIT).*** *As in Supplementary Fig. 12.*

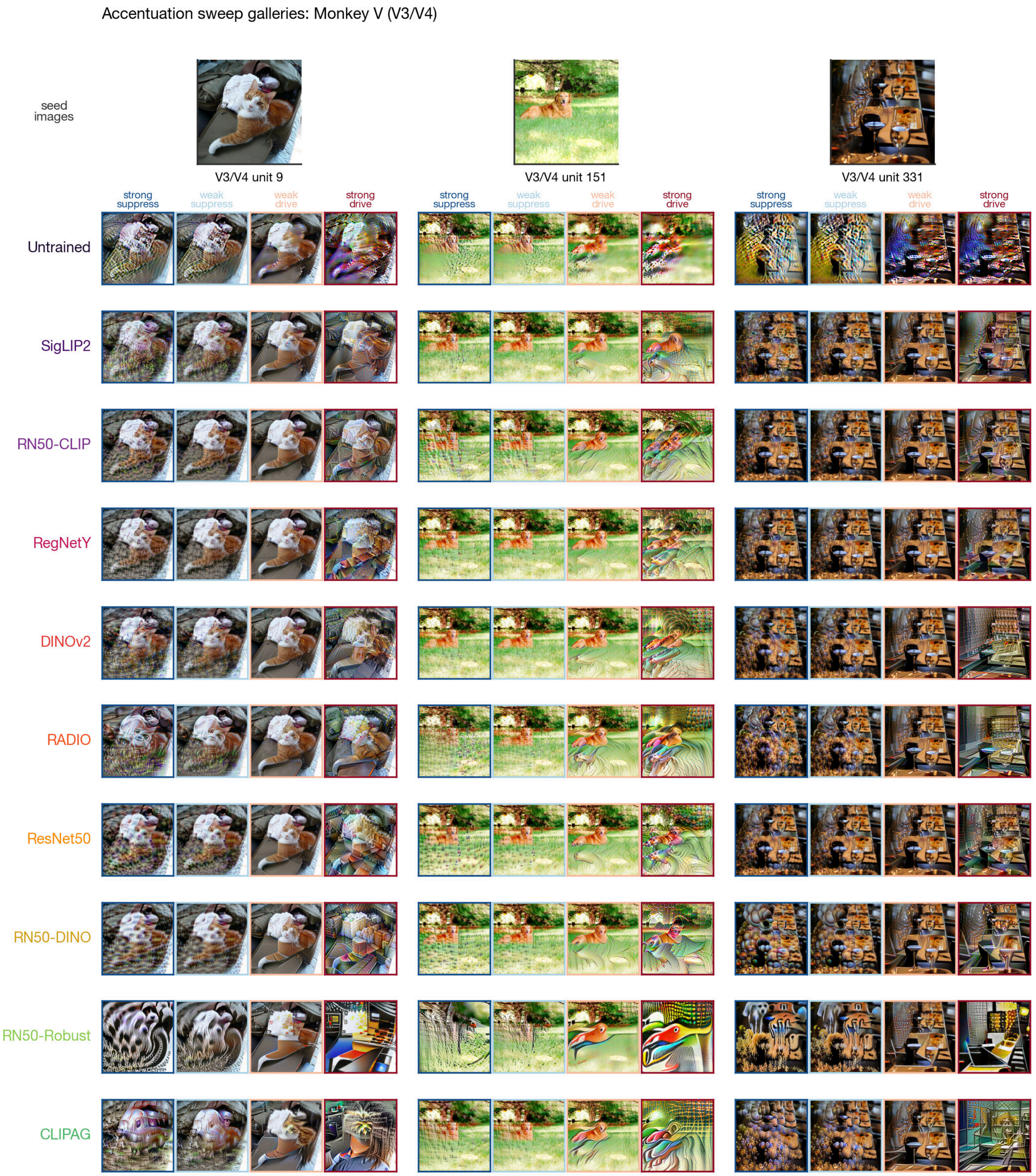


***Supplementary Figure 14. Accentuation sweep galleries – Monkey V (V3/V4).*** *As in Supplementary Fig. 12.*

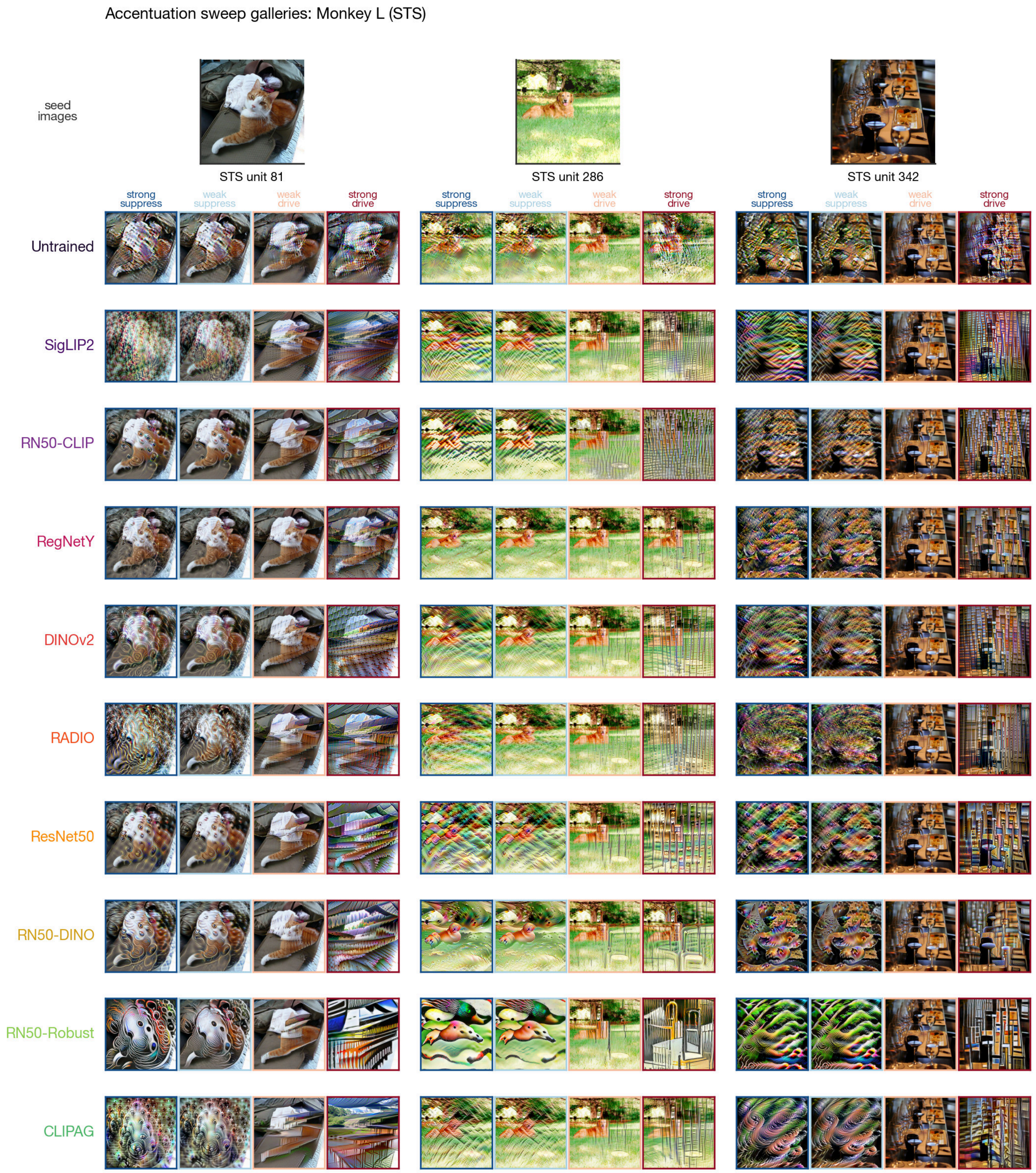


***Supplementary Figure 15. Accentuation sweep galleries – Monkey L (STS).*** *As in Supplementary Fig. 12.*

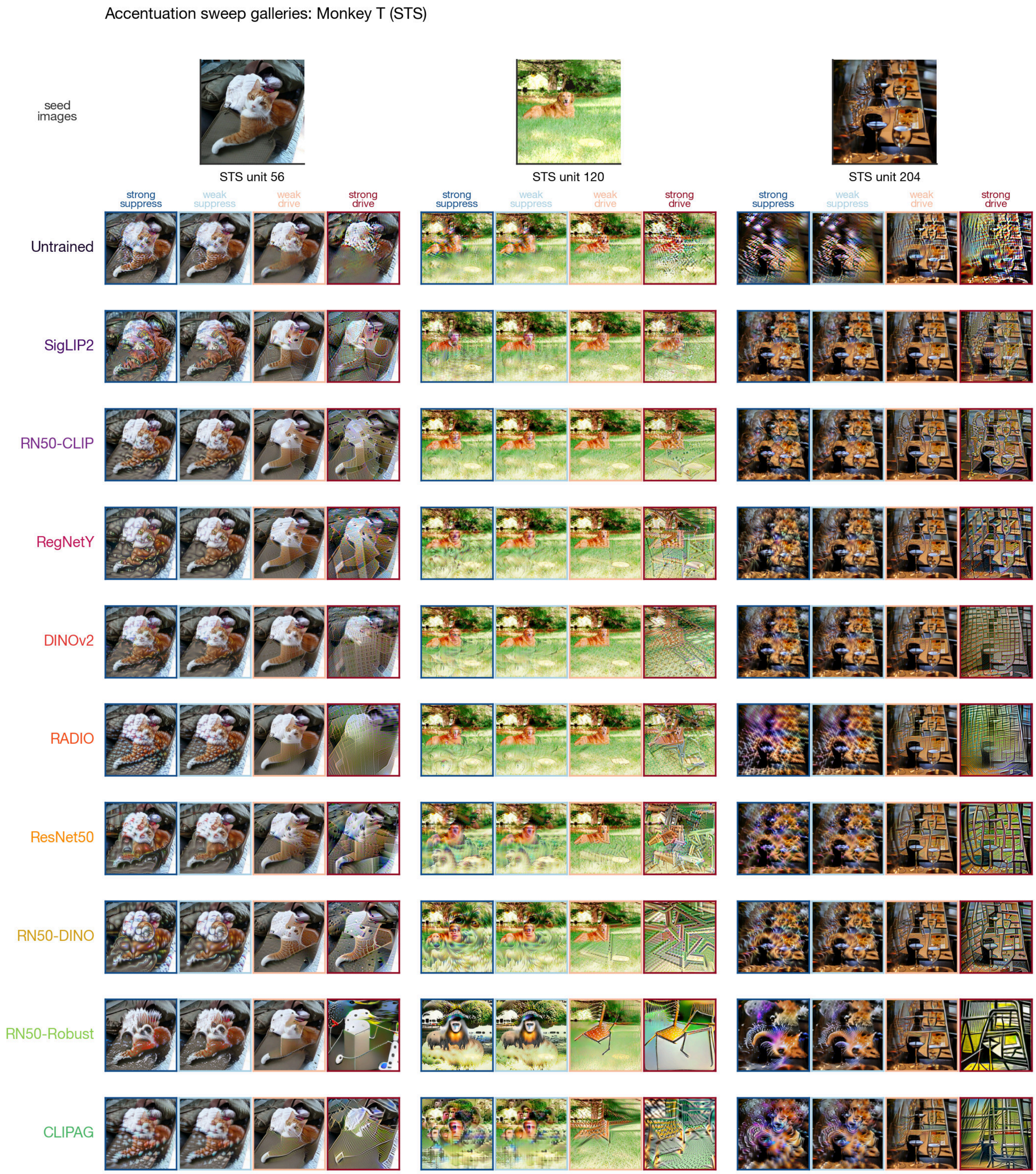


***Supplementary Figure 16. Accentuation sweep galleries – Monkey T (STS).*** *As in Supplementary Fig. 12.*

### Parametric neural control

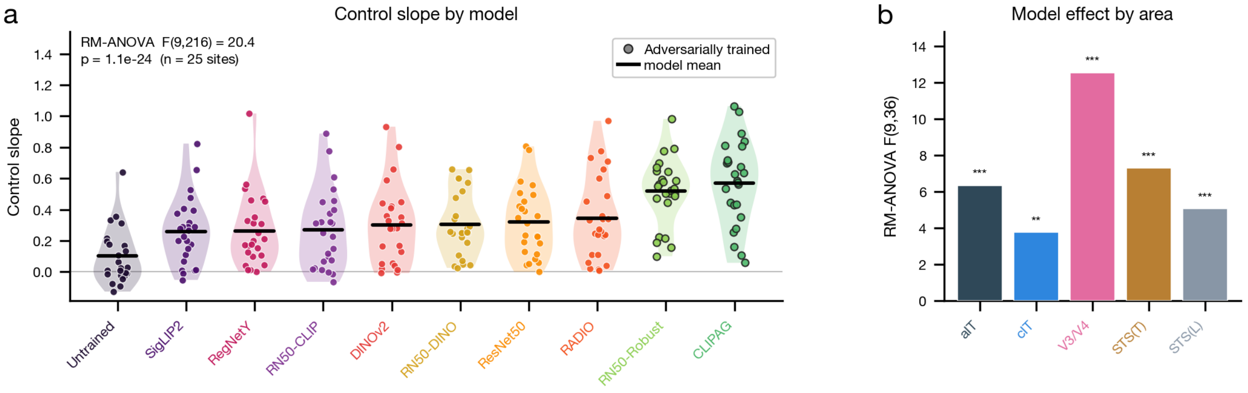


***Supplementary Figure 17. Model differences under the control-slope measure.*** *Repeating the Fig. 4 model-differentiation analysis using the control slope (slope of the OLS fit of measured against predicted responses ($z$-scored), using all accentuated stimuli for each site-model pair) in place of control $r$ recovers the same picture: models differ systematically, with the two adversarially trained models highest and the Untrained baseline lowest. (A) Per-model distribution of control slope across the 25 recorded sites (violin plus jittered swarm, one dot per site), with models ordered left to right by mean slope and each colored by model family (adversarially trained models, ResNet50-Robust and CLIPAG, highlighted with dark rings; Untrained as its own class). Black bars indicate model means; the gray line marks a slope of* 0*. A one-way repeated-measures ANOVA (model within-factor, $n = 25$ sites) confirms a strong model effect: $F(9, 216) = 20.4$, $p = 1.1 \times 10^{-24}$. (B) Per-area repeated-measures ANOVA $F(9, 36)$ (5 sites/area), colored by area/animal, with significance stars ($^*p < 0.05$, $^{**}p < 10^{-2}$, $^{***}p < 10^{-3}$). The model effect holds within every area (aIT, cIT, V3/V4, STS(T), STS(L)).*

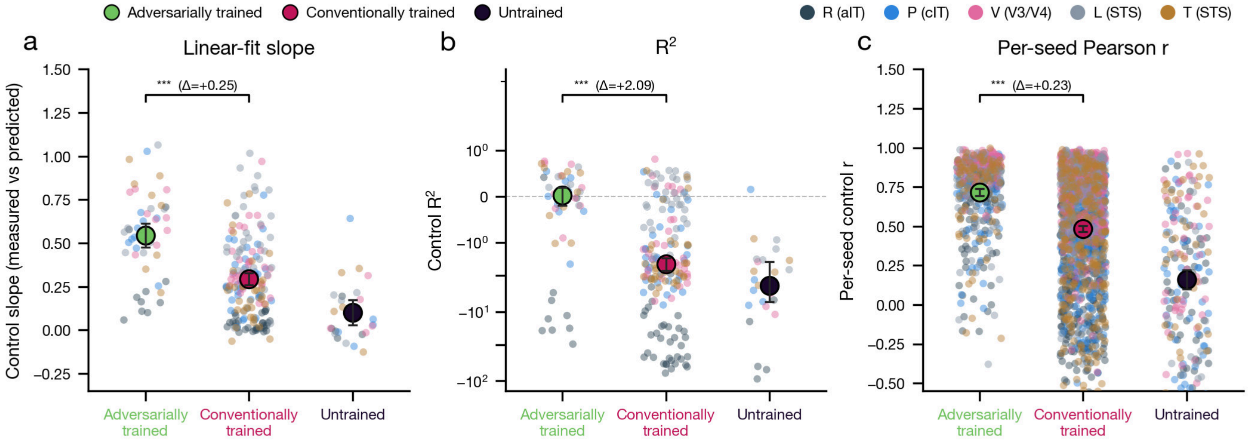


***Supplementary Figure 18. Control outcomes under alternative scoring metrics and per-seed aggregation.*** *The adversarially trained versus conventionally trained control comparison in the main text, scored as Pearson $r$ on seed-averaged accentuated sweeps, is re-run here under two alternative scores and one alternative aggregation. In each panel, one dot is an individual observation over the pooled accentuated stimuli, colored by monkey (Monkey R aIT, P cIT, V V3/V4, L STS, T STS), with the group central estimate and* 95% *CI overlaid, colored by training group (Adversarially trained green, Conventionally trained crimson, Untrained black). We report a two-sided Mann–Whitney $U$ test comparing adversarially trained against the conventionally trained models' scores. (A) Control slope, the OLS slope of measured versus predicted accentuated responses; group means $\pm$ $t$-based CIs; adversarially trained models exceed conventionally trained models ($p < 0.001$; $\Delta = +0.25$; separately, $\Delta = +0.26$ versus standard CNNs and $+0.24$ versus standard ViTs). (B) Control $R^2$, the coefficient of determination (symlog $y$-axis to accommodate the heavy negative tail); group medians with bootstrap CIs; adversarially trained models exceed conventionally trained models ($p < 0.001$; $\Delta = +2.09$; separately, $\Delta = +2.27$ versus standard CNNs and $+2.02$ versus standard ViTs). (C)*

*Per-seed control $r$, with each seed sweep scored separately rather than seed-averaged; group means $\pm$ $t$-based CIs; adversarially trained models exceed conventionally trained models ($p < 0.001$; $\Delta = +0.23$; separately, $\Delta = +0.22$ versus standard CNNs and $+0.25$ versus standard ViTs).*

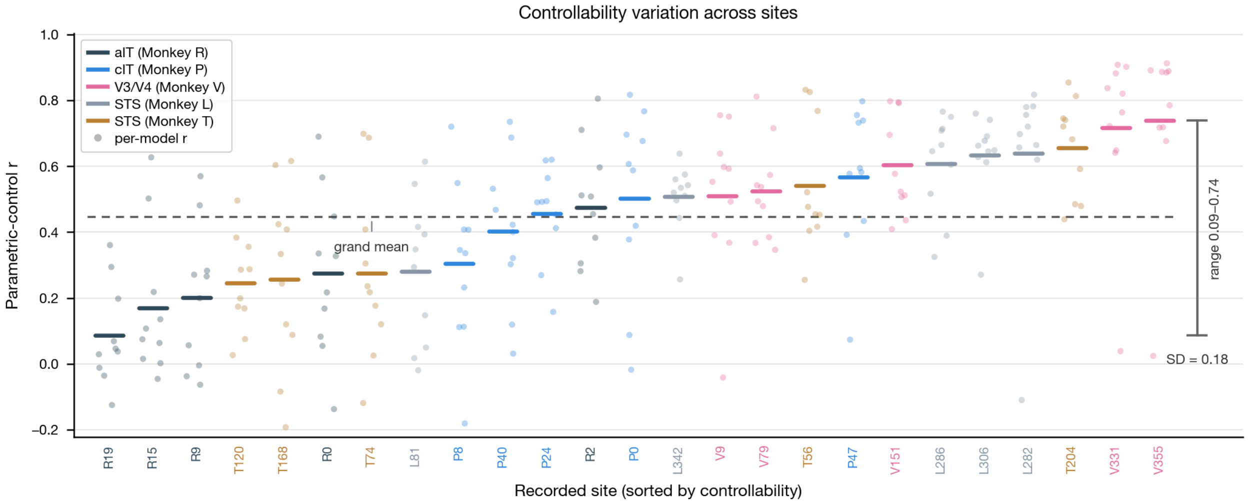


***Supplementary Figure 19. Controllability variation across recorded sites.*** *We plot the per-site control scores for the 25 measured sites, sorting them low-to-high and coloring by cortical area (aIT, cIT, V3/V4, STS). The control score here reflects the Pearson $r$ between a given model's predicted activation and the measured neural responses over its own personalized accentuated stimuli. Faint dots show the control $r$ across the 10 models, and the colored bar denotes the site mean. There are 250 total experiments shown. The dashed line indicates the grand mean across sites, and the annotation at right marks the min-to-max range of the site means. We find that mean control $r$ varies substantially across sites (SD $= 0.18$; range $0.09$–$0.74$), with the most controllable sites concentrated in V3/V4 and STS and the least controllable in aIT.*

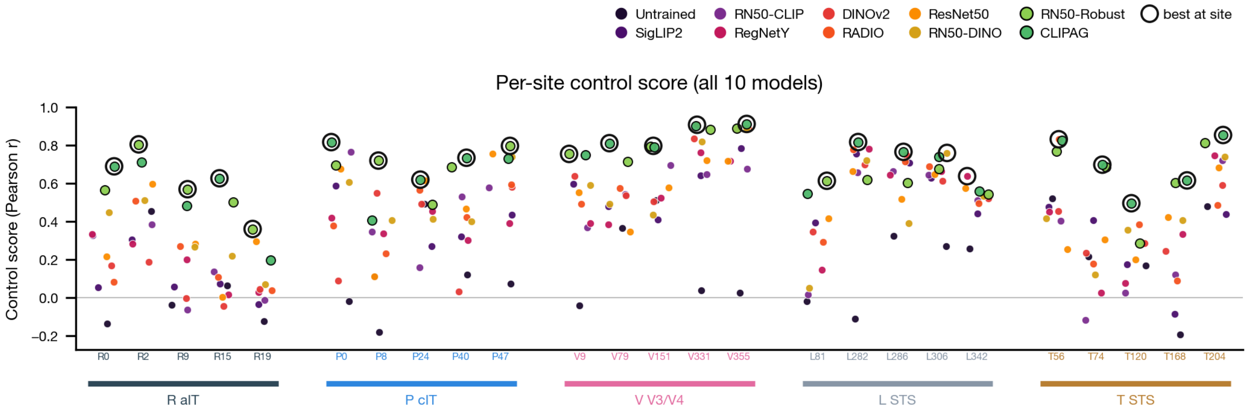


***Supplementary Figure 20. Per-site control scores for all ten models.*** *The same control score $r$ as defined in Supplementary Fig. S19 for each model $\times$ recording site ($n = 25$ across V3/V4, cIT, aIT, and STS in 5 macaques, tagged R/P/L/T/V), here emphasizing the ranking of models within each site. Per-site control $r$ for all 10 models, one point per model, colored by model and grouped by animal/area along the $x$-axis; adversarially trained models (ResNet50-Robust, CLIPAG) are drawn larger with black edges and the single best model at each site is ringed. An adversarially trained model is the top controller at $20/25$ sites ($80\%$).*

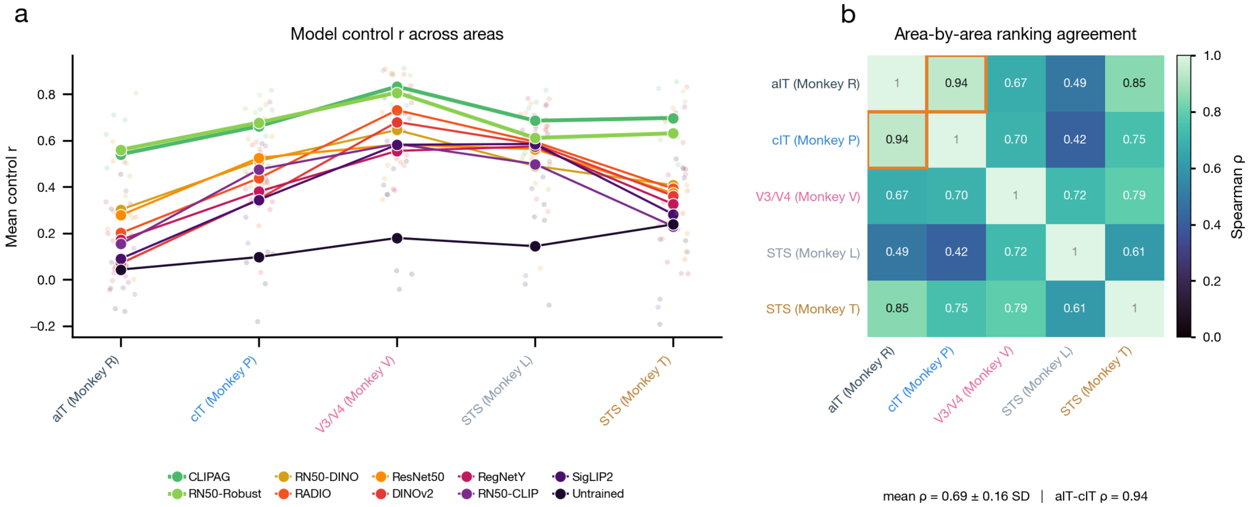


***Supplementary Figure 21. Cross-area consistency of model rankings.*** *Per-area, per-model mean control score (Pearson $r$ of predicted versus measured firing over pooled accentuated stimuli), for all 10 models (nine trained networks and the Untrained baseline), in five recording areas (one per animal): aIT, cIT, V3/V4, and two STS areas (STS-L, STS-T). (A) Each model's mean control score (y) across the five areas (x); one line per model, colored by model, with adversarially trained models (ResNet50-Robust, CLIPAG) drawn as thicker lines. Models are ordered by grand-mean control score; the rank ordering of models is preserved across areas (the two adversarially trained models highest throughout), while absolute control scores are lowest in aIT and peak in V3/V4. (B) Area-by-area matrix of Spearman rank correlations ($\rho$) between per-model control-score orderings, one value per off-diagonal cell (10 unique area pairs); color encodes $\rho \in [0, 1]$ (mako). The aIT/cIT pair, the strongest agreement ($\rho = 0.94$), is boxed in orange. Across the 10 pairs, mean $\rho = 0.69 \pm 0.16$ SD.*

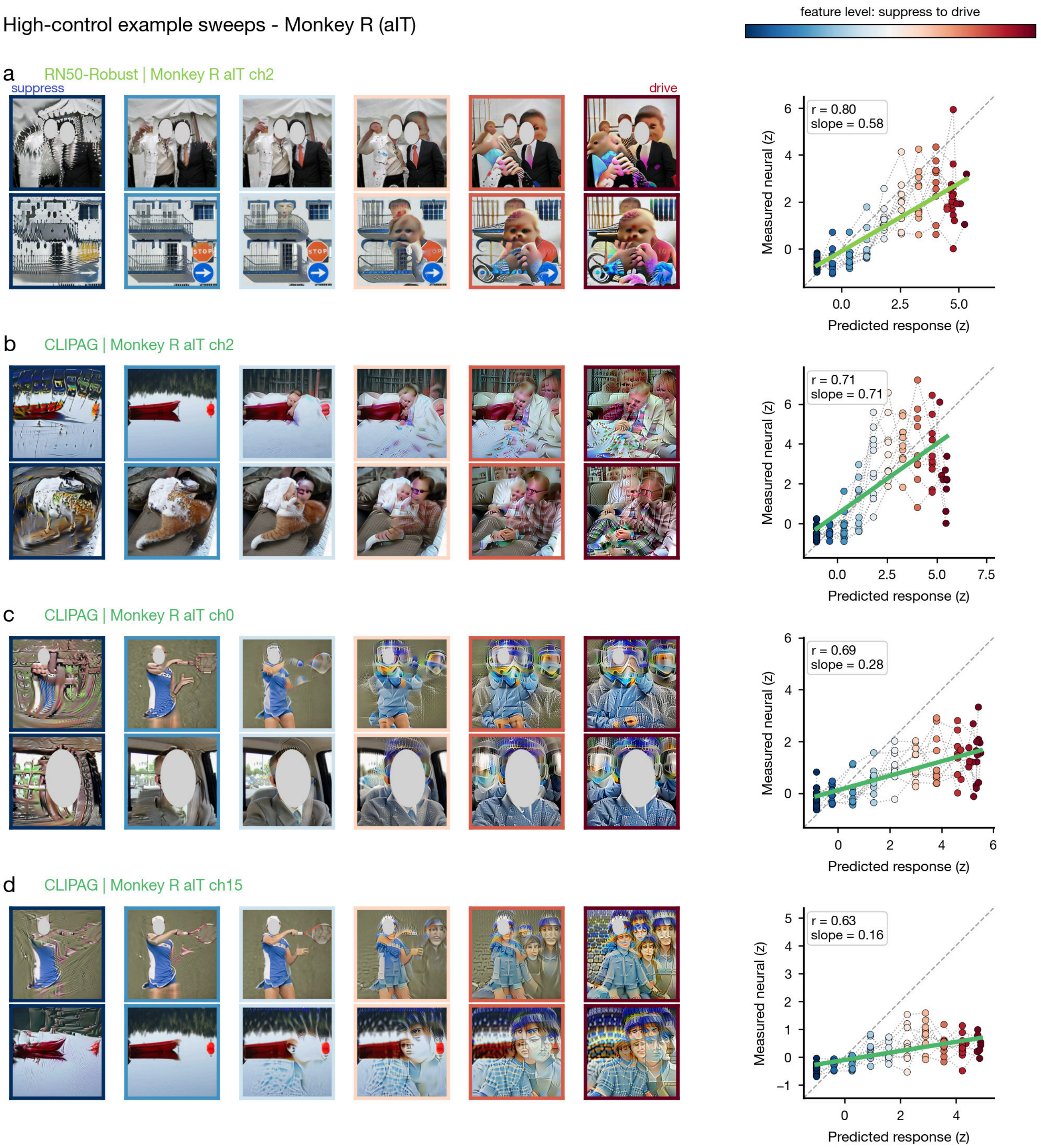


***Supplementary Figure 22. High-control example sweeps – Monkey R (aIT).*** *Each figure shows, for one macaque, the four site-model combinations with the highest parametric-control correlation. For each instance, we plot the two highest-$R^2$ seed sweeps (one seed per row, suppress levels to drive levels according to the color of the image borders) beside the predicted-versus-measured control scatter, in which thin lines link the eleven levels of each seed sweep. Dots are colored by feature level, and a solid line gives the least-squares fit with the pair's Pearson $r$ and slope outcome measures annotated. These examples illustrate strong predicted-to-measured correspondence across full accentuation sweeps.*

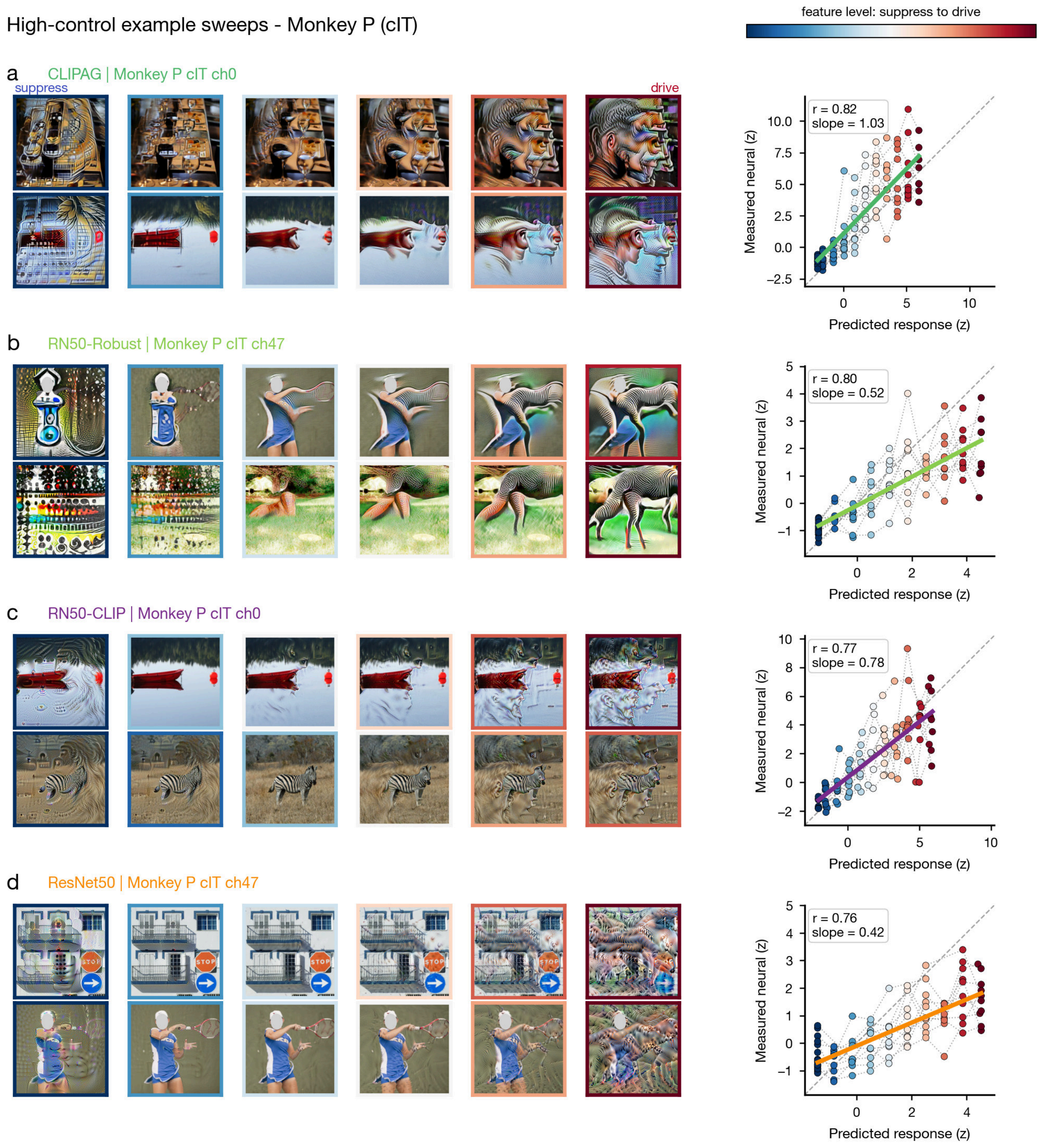


***Supplementary Figure 23. High-control example sweeps – Monkey P (cIT).*** *As in Supplementary Fig. 22.*

High-control example sweeps - Monkey V (V3/V4)

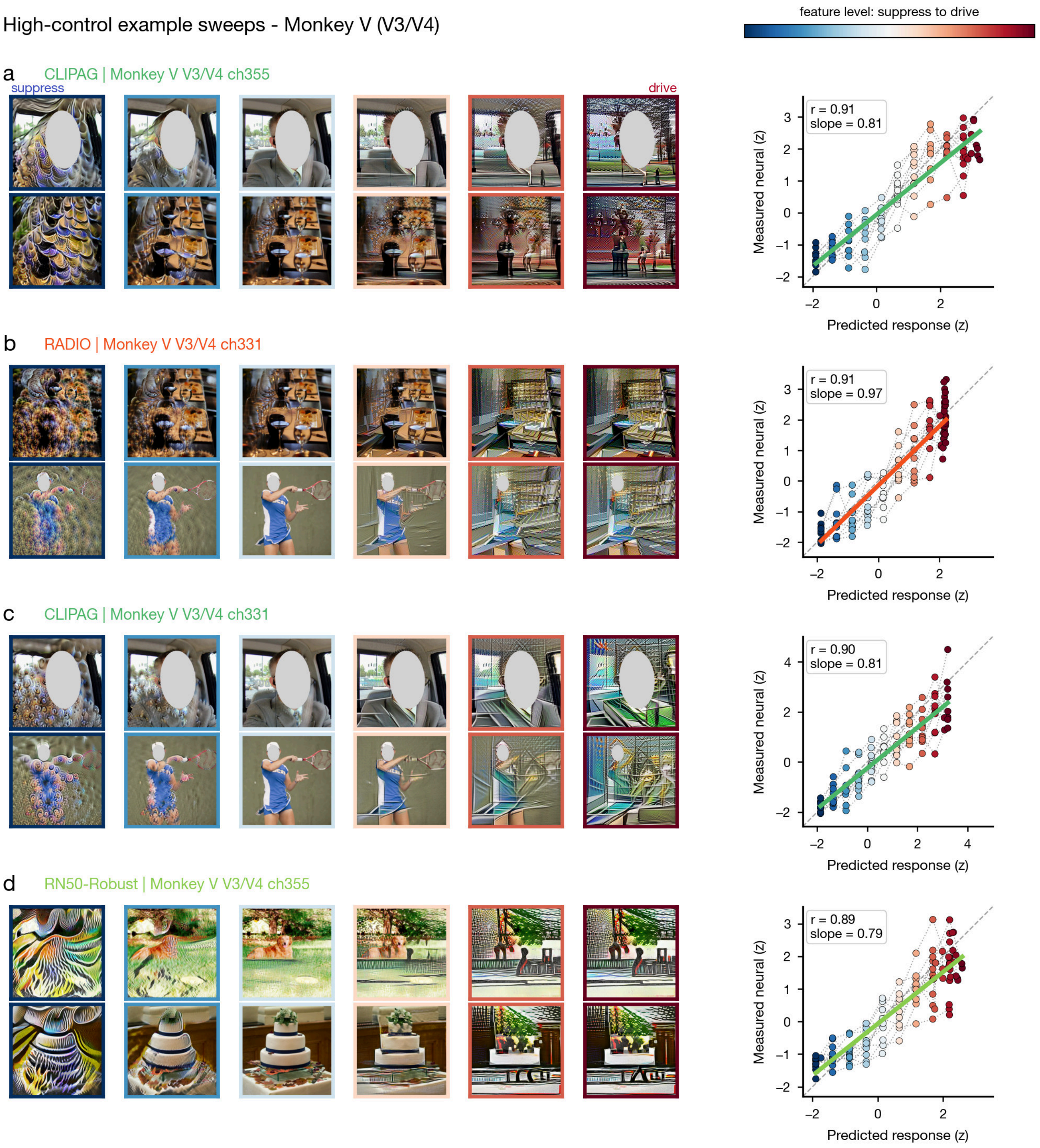


***Supplementary Figure 24. High-control example sweeps – Monkey V (V3/V4).*** *As in Supplementary Fig. 22.*

High-control example sweeps - Monkey L (STS)

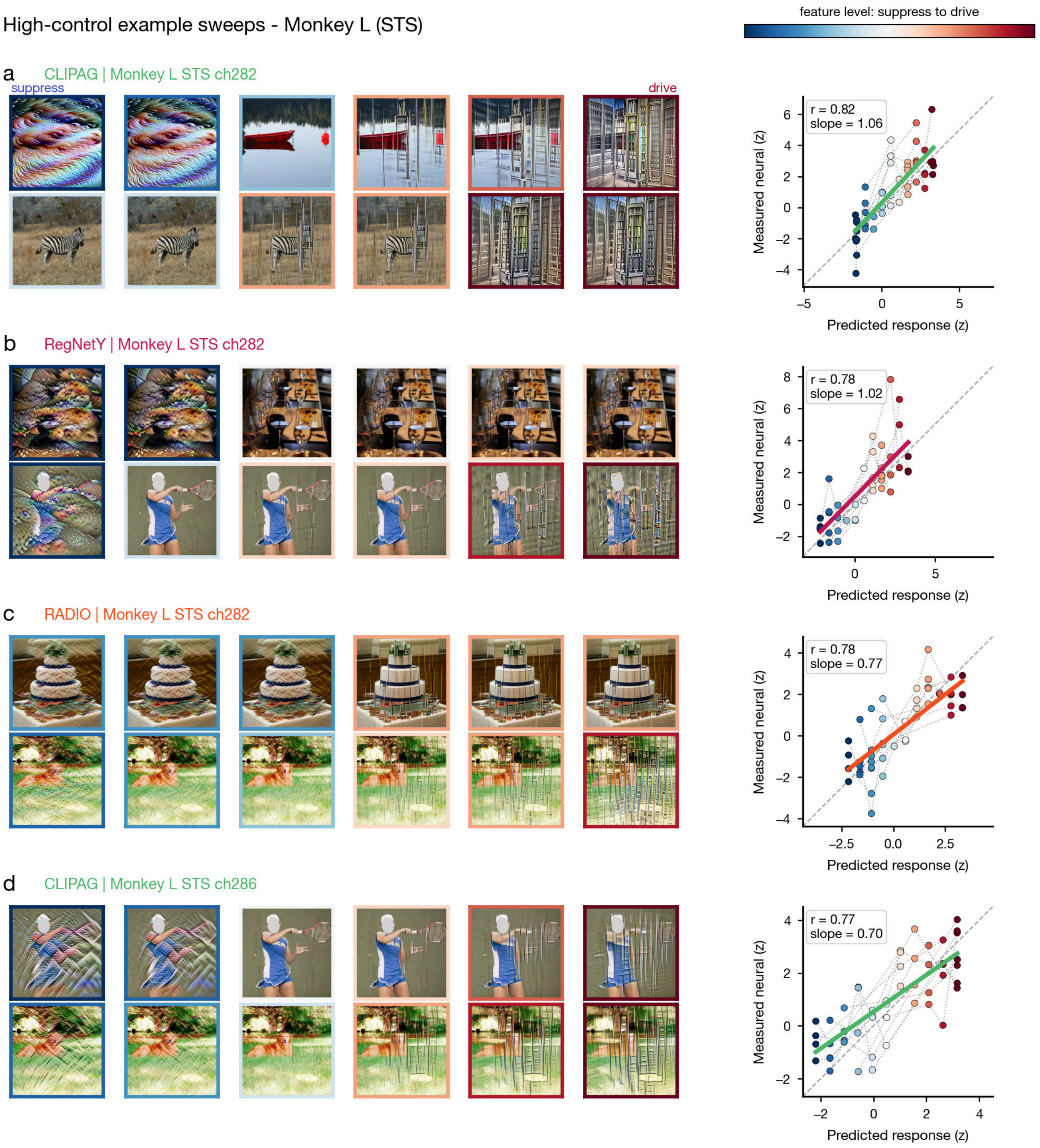


***Supplementary Figure 25. High-control example sweeps – Monkey L (STS).*** *As in Supplementary Fig. 22.*

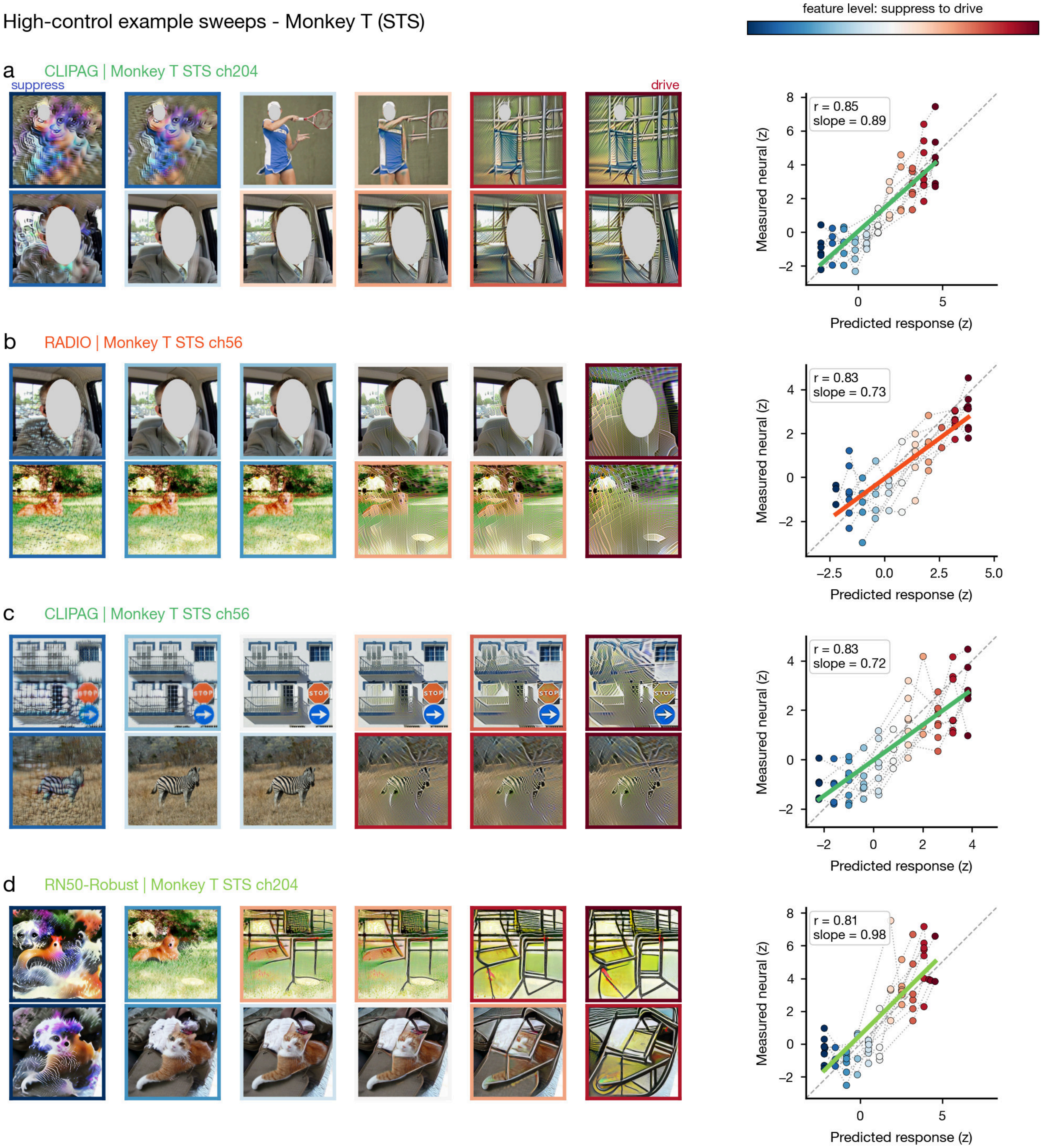


***Supplementary Figure 26. High-control example sweeps – Monkey T (STS).*** *As in Supplementary Fig. 22.*

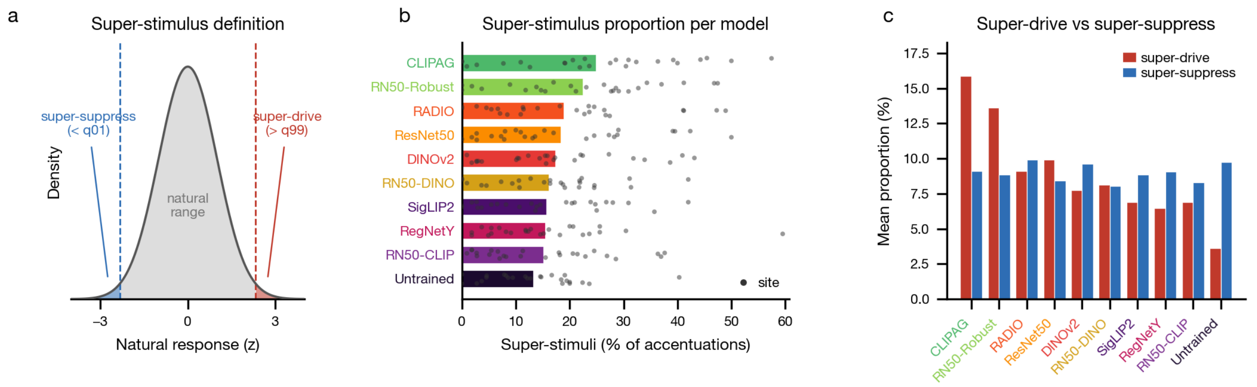


***Supplementary Figure 27. Super-stimulus proportion across models.*** *A super-stimulus is an accentuated image whose measured control response falls beyond the extremes of a site's natural response distribution; the two adversarially trained models produce the most. (A) Schematic defining the tails relative to a site's natural (calibration) response distribution: "super-drive" is a measured response above the* 99*th percentile (*$>$ q99*) and "super-suppress" is below the* 1*st percentile (*$<$ q01*), with the gray central region marking the natural range. (B) Per-model super-stimulus proportion (percent of a model's accentuated stimuli that are super-stimuli), mean over* $n = 25$ *sites (bars) with per-site points overlaid; models are ordered by their mean proportion and distinguished by color, with the adversarially trained models (CLIPAG, RN50-Robust) carrying the highest proportions and the Untrained baseline lowest. (C) Decomposition of the mean proportion into super-drive (red) versus super-suppress (blue) per model, same* 25 *sites.*

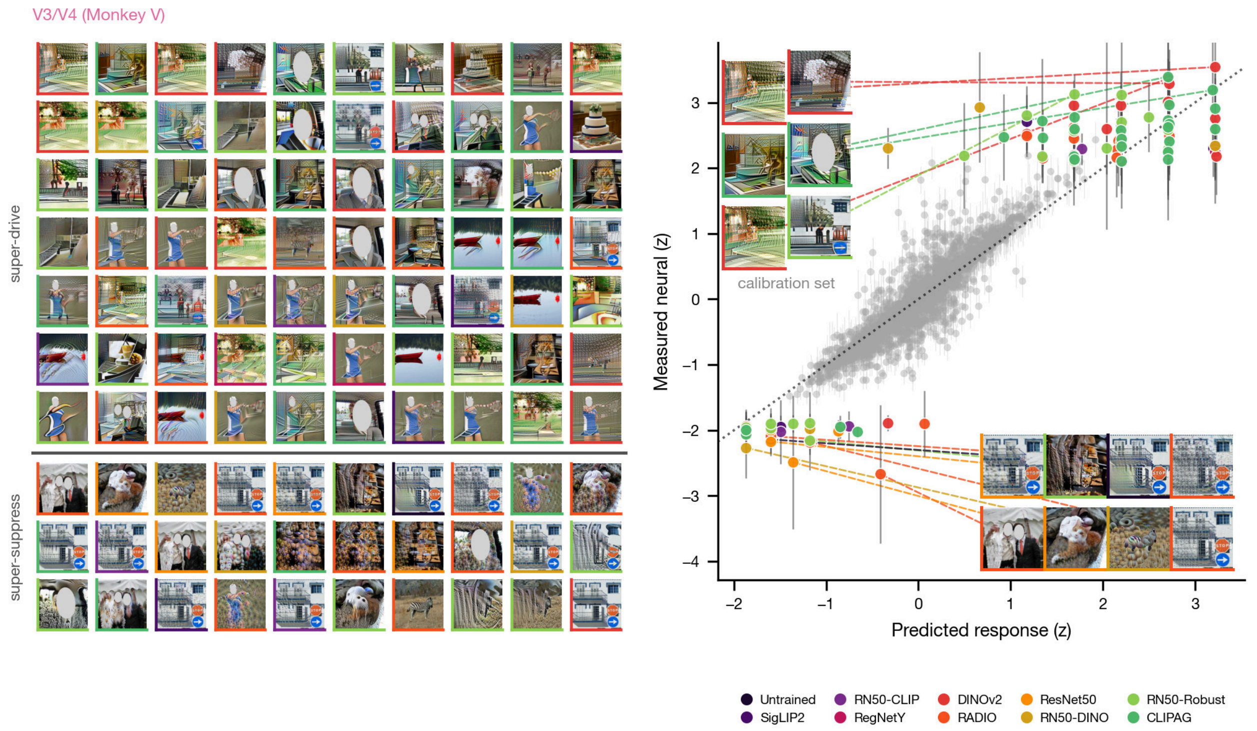


***Supplementary Figure 28. Super-stimulus response scatter, Monkey V.*** *Super-stimuli were accentuated images whose measured control response (with* $\geq 3$ *trial repeats) fell outside the site's natural-image calibration range:* $z >$ q99 *for super-drive and* $z <$ q01 *for super-suppress. (Left) A* $10 \times 10$ *gallery of the strongest super-stimuli, split into* 70 *super-drive (top) and* 30 *super-suppress (bottom). (Right) Predicted versus measured responses (*$z$ *units) for the same stimuli, shown as model-colored points with SEM whiskers. The gray calibration-set points with SEM whiskers follow the dotted identity line; the outermost super-stimuli are called out to their images with dashed leader lines. Image borders and points are colored by the generating model.*

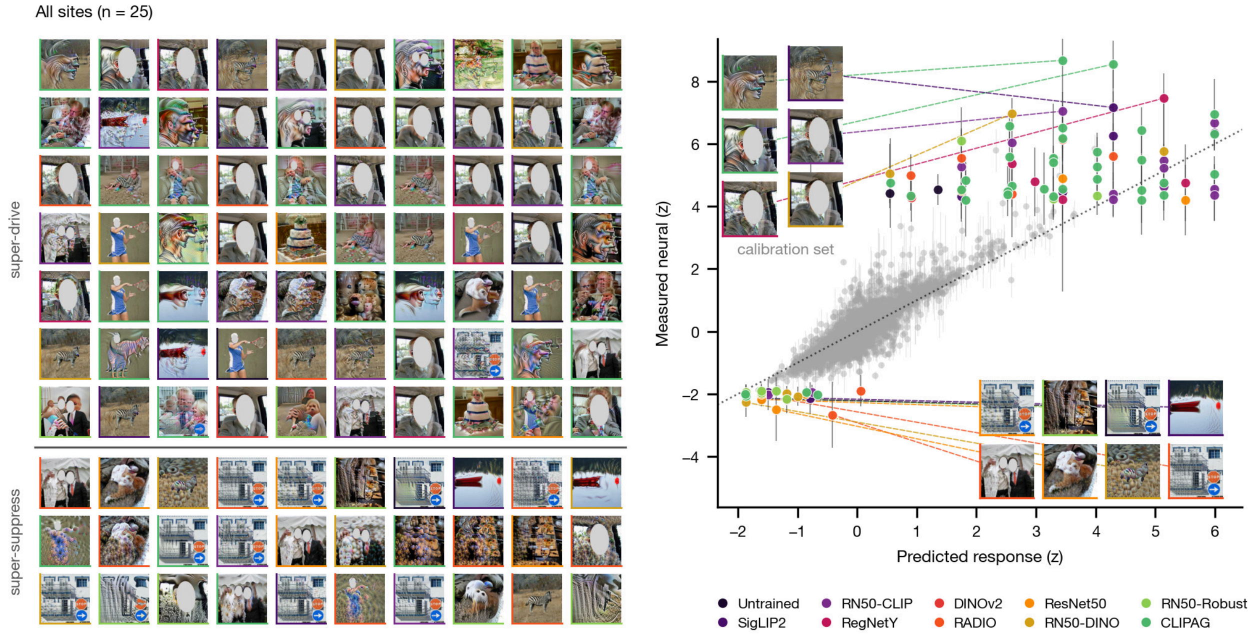


***Supplementary Figure 29. Super-stimulus response scatter, all sites.*** *Same format as above. Super-stimuli were accentuated images whose measured control response (with* $\geq 3$ *trial repeats) fell outside the site's natural-image calibration range:* $z > \mathrm{q99}$ *for super-drive and* $z < \mathrm{q01}$ *for super-suppress. (Left) A* $10 \times 10$ *gallery of the strongest super-stimuli pooled across all* $25$ *sites, split into* $70$ *super-drive (top) and* $30$ *super-suppress (bottom). (Right) Predicted versus measured responses (*$z$ *units) for the same stimuli, shown as model-colored points with SEM whiskers. The gray calibration-set points with SEM whiskers (*$n \leq 5000$ *pooled points) follow the dotted identity line; the outermost super-stimuli are called out to their images with dashed leader lines. Image borders and points are colored by the generating model.*

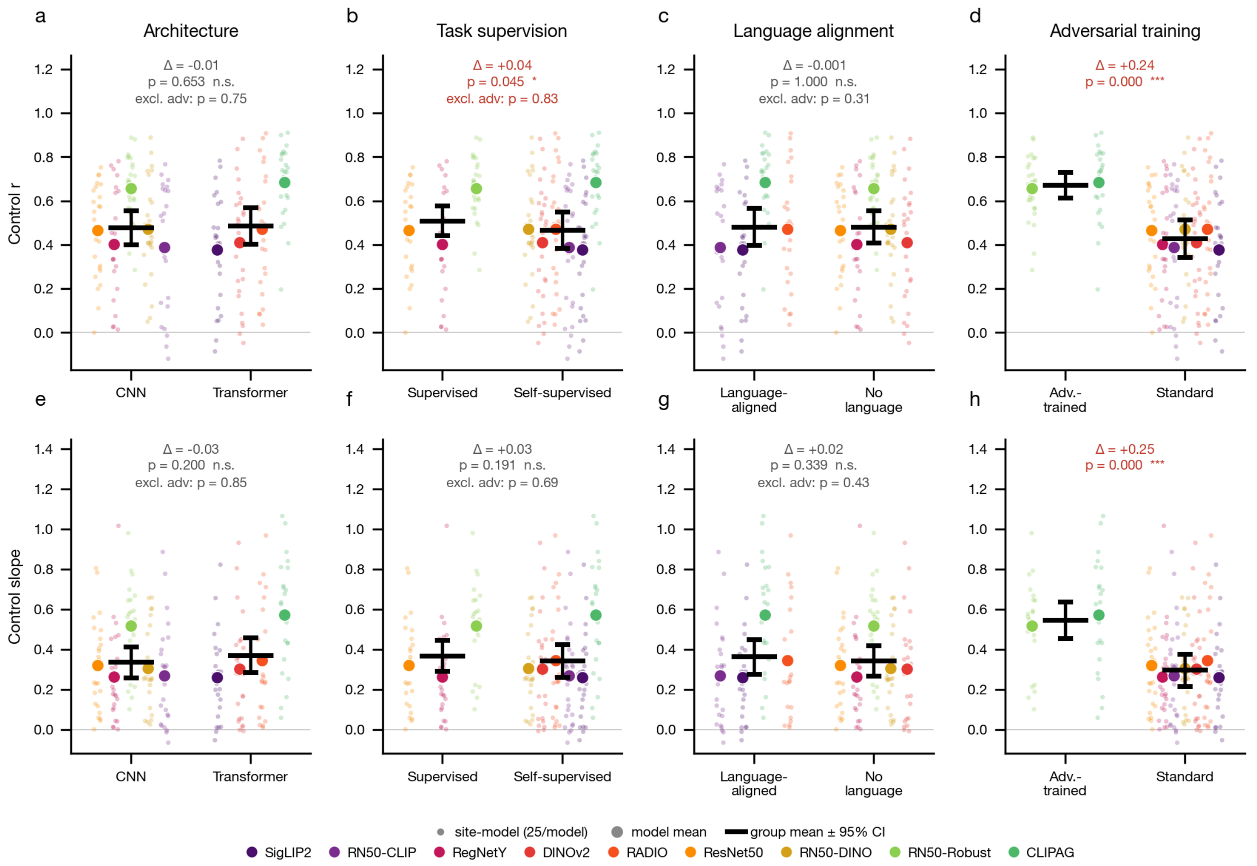


***Supplementary Figure 30. Neural control across model inductive bias groups.*** *Test of whether different inductive biases separate parametric control; each split is evaluated within site (Wilcoxon signed-rank over the 25 recording sites on the per-site difference of group-mean control), among the 9 trained models. Top row, control $r$; bottom row, control slope. Faint dots reflect the 25 site-level values per model, larger dots reflect per-model means colored by model; black bar is the group mean ±95% CI over sites; annotations show the group difference $\Delta$, its $p$, and the $p$ recomputed excluding the two adversarially trained models. (A) Convolutional versus transformer architecture ($\Delta = -0.01$, $p = 0.653$). (B) Label-supervised versus self-supervised objective ($\Delta = +0.04$, $p = 0.045$), the only weakly significant effect outside of adversarial training, which disappears once ResNet50-Robust and CLIPAG are removed (excl. adv $p = 0.83$). (C) Language-aligned versus vision-only ($\Delta = -0.001$, $p = 1.000$). (D) Adversarially trained versus standard ($\Delta = +0.24$, $p < 0.001$), the only strong dissociation. (E)–(H) The same analysis is repeated for the control slope outcome measure, with similar outcomes: architecture ($\Delta = -0.03$, $p = 0.200$), objective ($\Delta = +0.03$, $p = 0.191$), and language ($\Delta = +0.02$, $p = 0.339$) are all null, whereas adversarial training again separates the groups ($\Delta = +0.25$, $p < 0.001$).*

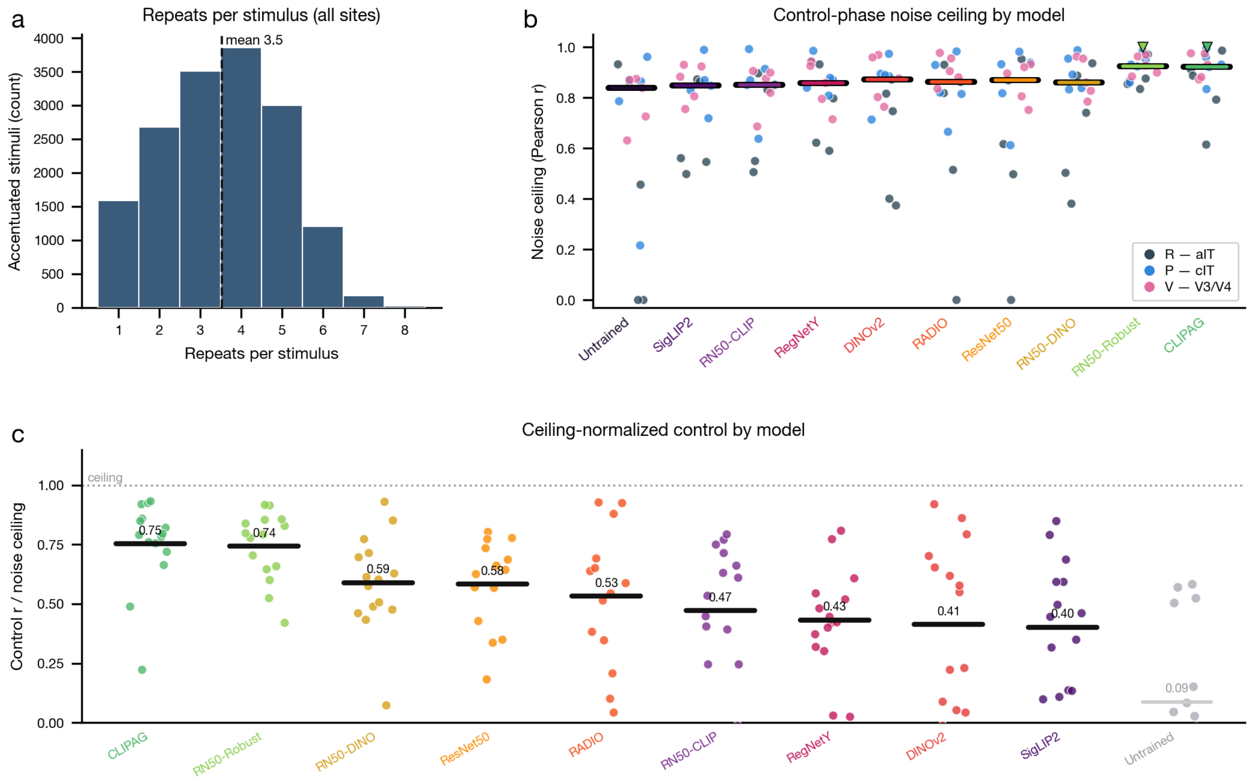


***Supplementary Figure 31. Reliability of control-phase responses.*** *(A) Distribution of repeat counts pooled across all accentuated stimuli and control sessions (dashed line, mean* $= 3.5$*). (B) Control-phase noise ceiling (NSD-style Pearson* $r$*) per encoding model, one dot per site-model combination where the ceiling is defined, colored by animal and area (R/aIT, P/cIT, V/V3/V4;* $n = 15$ *sites per model,* $150$ *total); the two STS animals (L, T) lacked sufficient same-day anchor repeats and are omitted. Thick bars indicate per-model medians; triangles and bold labels mark the two adversarially trained models (ResNet50-Robust, CLIPAG). We find that noise ceilings are broadly high across all* $10$ *models (medians* $\approx 0.8$*–*$0.9$*), including the Untrained baseline, confirming that measured control-phase responses are reliable. (C) Control score* $r$ *normalized by the control-phase noise ceiling shown in (B); the ranking of models is largely unchanged. Further, it suggests the achieved control scores of adversarially trained models are decent compared to the noise ceiling.*

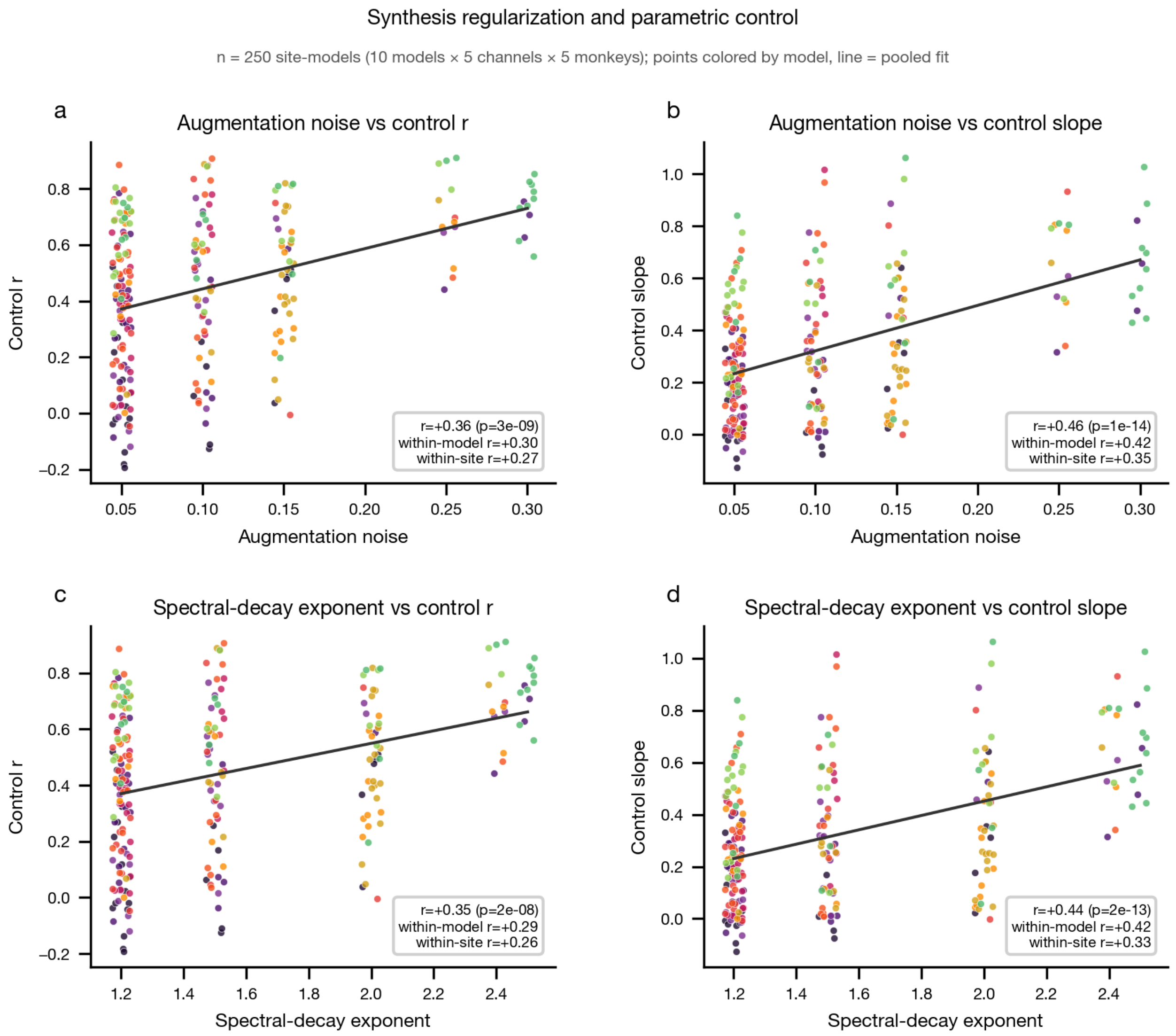


***Supplementary Figure 32. Synthesis regularization and parametric control.*** *The automatically selected synthesis regularization (augmentation noise and spectral-decay exponent) covaries with control fidelity across all* $n = 250$ *site-models (10 models* $\times$ *5 channels* $\times$ *5 monkeys). In (A)–(D) each dot is one site-model colored by model, the dark line is the pooled OLS fit, and inset text reports the pooled Pearson* $r$ *(with* $p$*), the within-model partial* $r$ *(predictor and outcome centered per model), and the within-site partial* $r$*. (A) Augmentation noise versus control score* $r$ *(Pearson correlation of predicted vs measured over accentuated stimuli): pooled* $r = +0.36$ *(*$p = 3 \times 10^{-9}$*), within-model* $r = +0.30$*, within-site* $r = +0.27$*. (B) Augmentation noise versus control slope (OLS gain): pooled* $r = +0.46$ *(*$p = 1 \times 10^{-14}$*), within-model* $r = +0.42$*, within-site* $r = +0.35$*. (C) Spectral-decay exponent versus control score: pooled* $r = +0.35$ *(*$p = 2 \times 10^{-8}$*), within-model* $r = +0.29$*, within-site* $r = +0.26$*. (D) Spectral-decay exponent versus control slope: pooled* $r = +0.44$ *(*$p = 2 \times 10^{-13}$*), within-model* $r = +0.42$*, within-site* $r = +0.33$*. Higher regularization tracks better control both across and within models.*

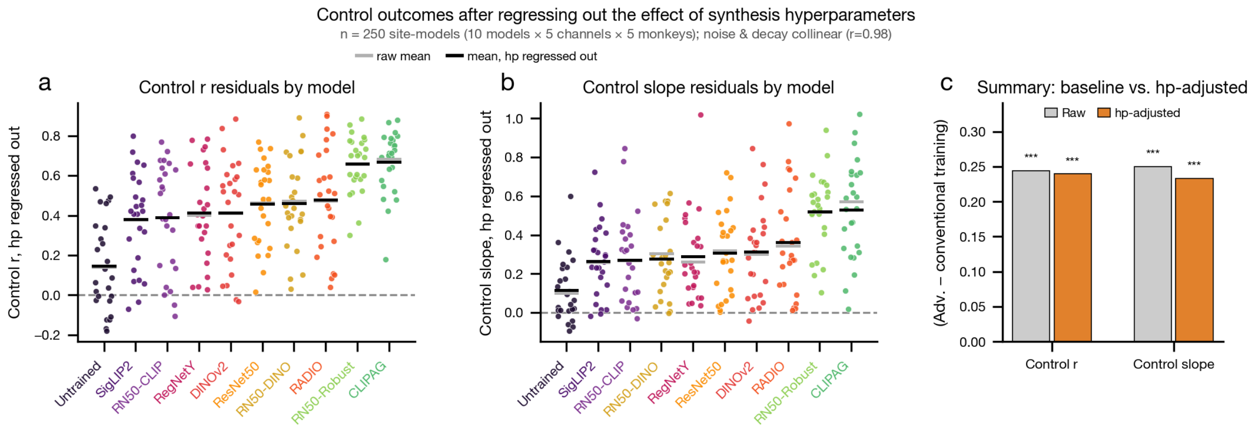


***Supplementary Figure 33. Control outcomes after regressing out the effect of synthesis hyperparameters.*** *This analysis examines whether the adversarially trained models' control advantage is related to the synthesis hyperparameters (hp) that were involved in generating the accentuations, i.e., the two regularization knobs, augmentation noise and spectral decay (Supplementary Fig. S6). Across all* $n = 250$ *site-models (10 models* $\times$ *5 sites* $\times$ *5 macaques), a single OLS model,* $\text{outcome} \sim \text{model} + \text{noise} + \text{decay} + \text{site}$, *was fit for each control outcome, and the part of the outcome attributed to the two knob terms was subtracted. The knobs rise together along the regularization ladder and are nearly collinear (*$r = 0.98$*), so their individual coefficients are not meaningful; only their combined contribution is used, and including both removes as much hp-related variance as possible. (A, B) Control* $r$ *(A) and control slope (B) with hp regressed out, one dot per site-model, colored by model; black bars, hp-adjusted model means; gray bars, raw means; dashed line at zero; models ordered by adjusted mean, adversarially trained models in bold. (C) We assess the gap in control outcomes between the adversarially trained and conventionally trained model families, comparing raw and hp-adjusted scenarios (raw* $p$ *from a within-site paired Wilcoxon over the* $25$ *sites; hp-adjusted* $p$ *from an ANCOVA with site fixed effects and site-clustered errors;* $***\ p < 10^{-3}$*).*

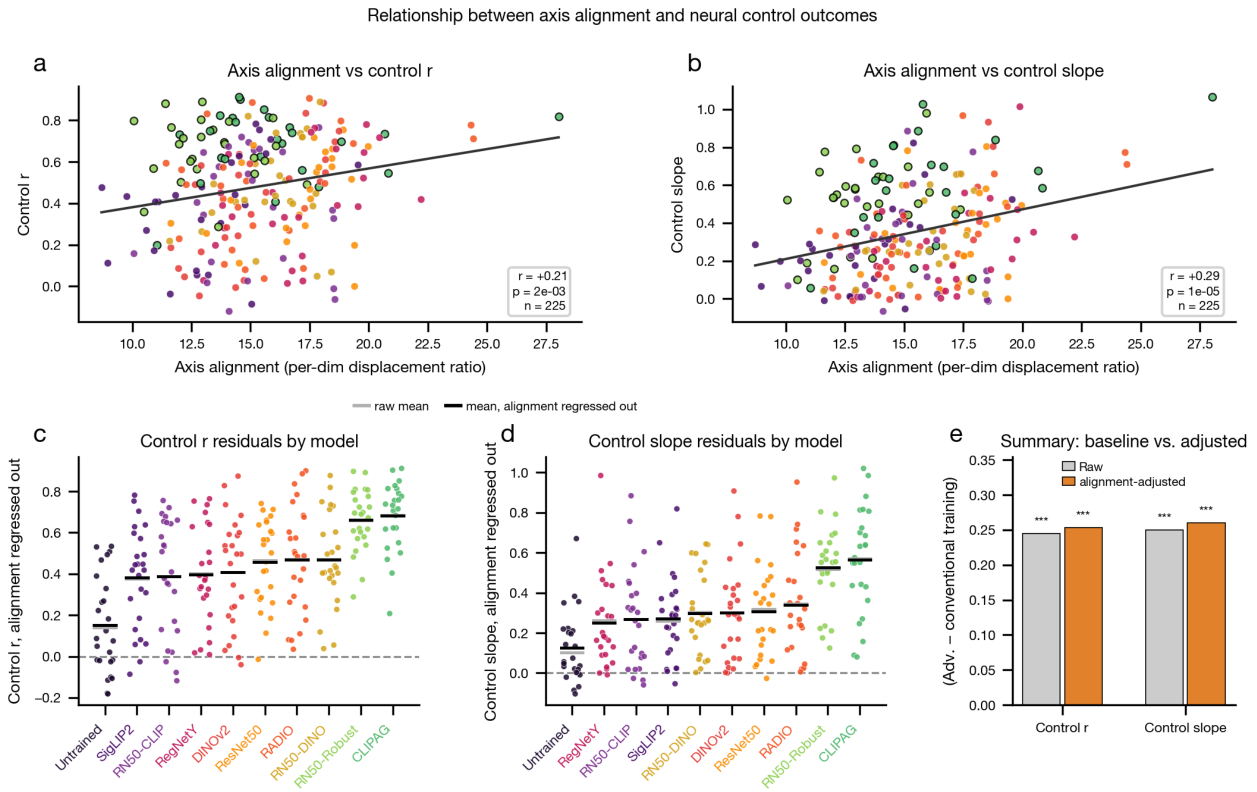

***Supplementary Figure 34. Relationship between axis alignment and neural control outcomes.*** *For each model-channel, axis alignment is the per-dimension displacement ratio* $\mathrm{DM}/(\mathrm{DOM}/\sqrt{n_{\mathit{off}}})$*, seed-averaged over the 10 accentuation seeds, quantifying how much of each parametric sweep lands on the encoding axis. (A) Control score (*$r$*) versus axis alignment across the trained model-channels (*$n = 225$*; 9 trained models, Untrained excluded), dots colored by model with adversarially trained models (ResNet50-Robust, CLIPAG) outlined in black; OLS fit and Pearson* $r = +0.21$ *(*$p = 2 \times 10^{-3}$*). (B) Same for control slope,* $r = +0.29$ *(*$p = 1 \times 10^{-5}$*). Alignment is weakly but positively associated with control, motivating adjustment. (C, D) Control r (C) and control slope (D) with alignment regressed out, one dot per site-model across all 10 models (alignment effects removed via a joint OLS fit with model and site terms, grand mean restored); black bars, alignment-adjusted model means; gray bars, raw means. The model ordering is preserved, with the two adversarially trained models remaining highest. (E) Adversarially-trained-minus-conventional group-mean gap for control r and control slope among the 9 trained models, raw (gray; p from a within-site paired Wilcoxon over the 25 sites) versus alignment-adjusted (orange; ANCOVA with site fixed effects and site-clustered errors); all four gaps are positive and significant (*$***\, p < 10^{-3}$*), showing the adversarially trained models' advantage is not explained by greater on-axis displacement.*

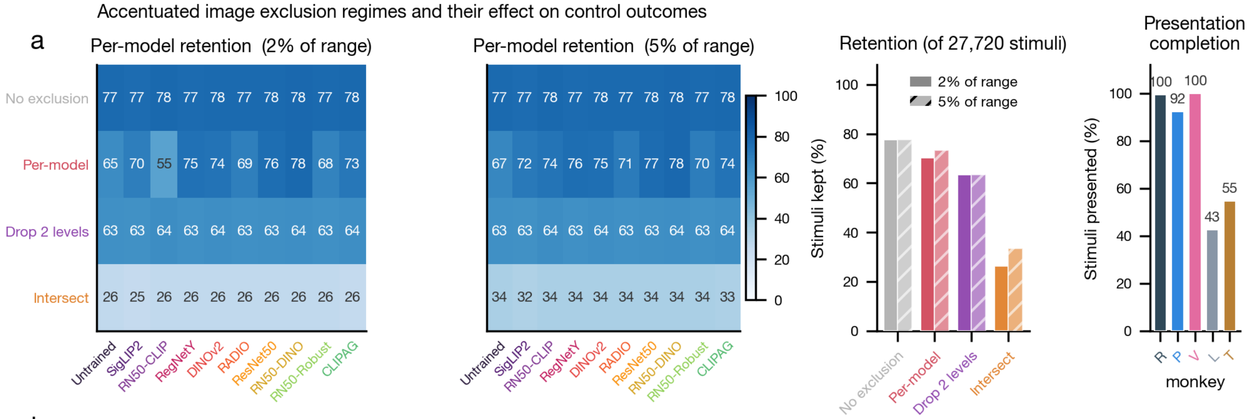


b

| Criterion | Definition | Stimuli kept 2% / 5% of range | Usable sweeps ≥ 5 2% / 5% of range | Seed-conds all ≥ 5 2% / 5% of range |
|---|---|---|---|---|
| No exclusion | all presented stimuli kept | 21,498 (78%) / 21,498 (78%) | 2191 / 2191 (of 2,500) | 155 / 155 (of 250) |
| Per-model | each model drops its own failed syntheses (stimulus sets differ across models) | 19,456 (70%) / 20,321 (73%) | 2080 / 2134 (of 2,500) | 123 / 142 (of 250) |
| Drop 2 levels | the two strongest drive levels dropped from every sweep (same levels for all models) | 17,584 (63%) / 17,584 (63%) | 1973 / 1973 (of 2,500) | 150 / 150 (of 250) |
| Intersect | a (site, seed, level) kept only if every model succeeded there (identical sets) | 7,260 (26%) / 9,240 (33%) | 870 / 1080 (of 2,500) | 87 / 108 (of 250) |

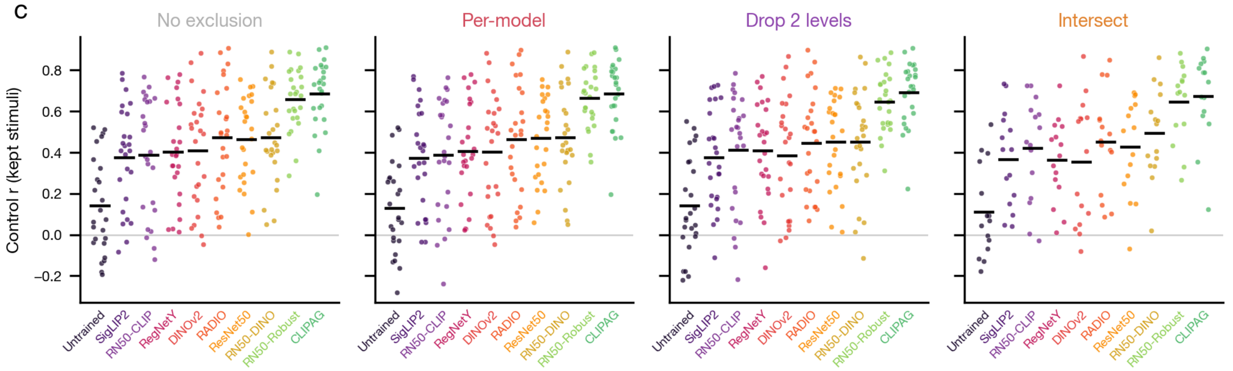


***Supplementary Figure 35. Control outcomes across accentuated-image exclusion regimes.*** *We defined failure of accentuation synthesis as occurring when the generating model's own post-hoc prediction for the saved image misses its intended target in either direction by more than a tolerance value* $\tau$ *(a fraction of the channel's target range). Dropping each model's own failures would compare models on unequal footing; we therefore derived a set of exclusion regimes to ensure fair comparison between models, and here we test whether control outcomes are sensitive to this choice. (A) Per-model retention under each regime at* $\tau = 2\%$ *and* $5\%$ *of range (heatmaps), overall retention of the*

27,720 *synthesized stimuli (bars), and presentation completion per monkey (the STS monkeys did not complete their stimulus sets,* 43% *and* 55% *presented; unpresented stimuli are excluded under every regime). (B) Summary table: what each regime keeps and how much survives. No exclusion retains all presented stimuli* (21,498 *of the synthesized set;* 78%*); per-model drops each model's own failed syntheses* (70%/73% *at* $\tau = 2\%/5\%$*); drop-2-levels removes the two strongest drive levels from every sweep* (63%)*; intersect keeps a (site, seed, level) cell only if every model succeeded there* (26%/33%)*. (C) Per-model control* $r$ *recomputed from the kept stimuli under each regime* ($\tau = 5\%$)*, one subplot per regime with fixed model order and a shared y-axis: the model ordering, and the adversarially trained models' advantage, are essentially unchanged across regimes.*

#### Controversial accentuations

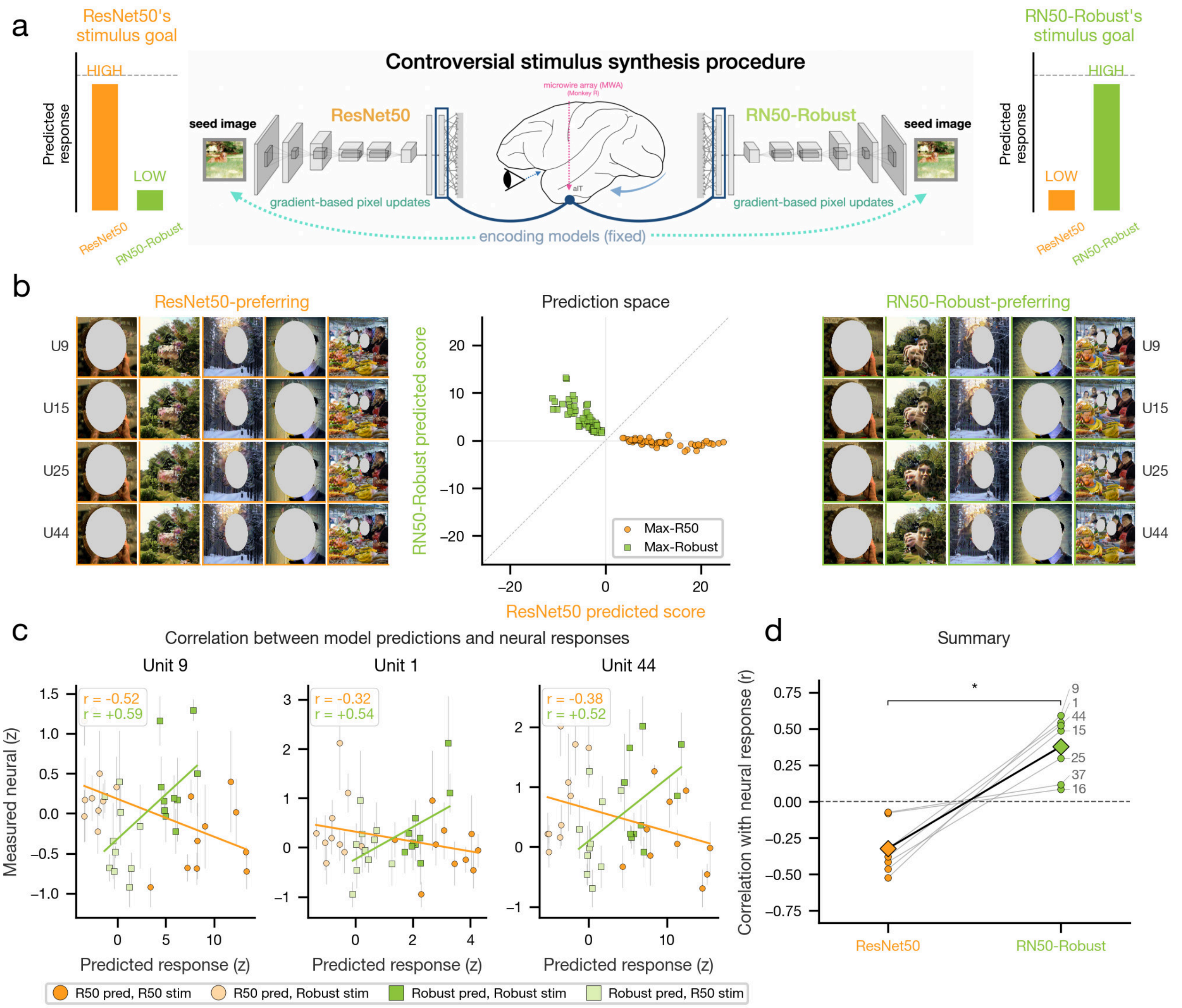


***Supplementary Figure 36. Controversial-accentuation experiment.*** *(A) Procedure for synthesizing controversial stimuli. For each neural site, ResNet50 and ResNet50-Robust encoding models define separate prediction axes. ResNet50-preferring stimuli were optimized for high predicted responses under ResNet50 and low predicted responses under ResNet50-Robust, and ResNet50-Robust-preferring stimuli were optimized for the reverse pattern. (B) Example controversial accentuations for four aIT sites. ResNet50-preferring stimuli are shown at left, and ResNet50-Robust-preferring stimuli at right. The center plot shows the prediction space for one example site, with each stimulus plotted*

*according to its predicted response under ResNet50 and ResNet50-Robust; orange circles indicate ResNet50-preferring stimuli, and green squares indicate ResNet50-Robust-preferring stimuli. (C) Relationship between model predictions and neural responses for three example aIT sites. Neural responses are in standardized firing rate units, and Pearson correlations are shown for each model evaluated on its predictions over both models' preferred stimuli. (D) Summary across the seven targeted aIT sites. Each dot indicates the Pearson correlation between one site's observed responses and predictions from ResNet50 or ResNet50-Robust, evaluated over both groups of stimuli (20 stimuli per site); diamonds indicate across-site means, and gray lines connect paired correlations within a site (Wilcoxon signed-rank test). ResNet50-Robust predictions correlate positively with neural responses, whereas ResNet50 predictions generally correlate negatively.*

Controversial-accentuation stimuli (Monkey R, aIT; 7 sites x 10 seeds x 2 directions = 140)

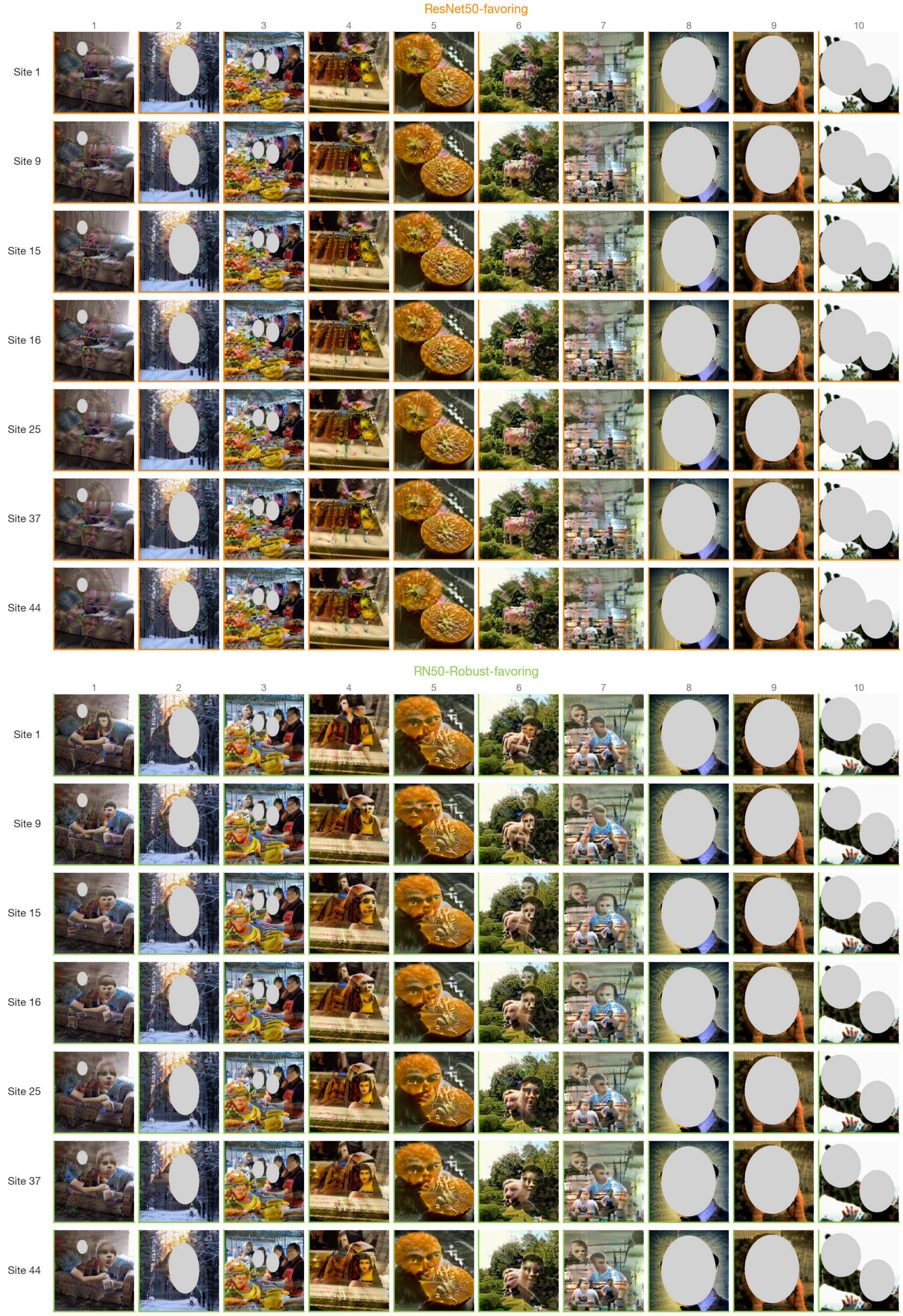

***Supplementary Figure 37. Complete controversial-accentuation stimulus set.*** *The full set of* 140 *controversial accentuated stimuli from the ResNet50 versus ResNet50-Robust experiment (Monkey R, area aIT,* 7 *sites), each synthesized to drive one model's encoding prediction while suppressing the other's. Top mosaic, ResNet50-favoring stimuli (orange borders); bottom mosaic, ResNet50-Robust-favoring stimuli (green borders). Rows index the* 7 *targeted aIT sites (channels* 1, 9, 15, 16, 25, 37, 44*); columns index the* 10 *seed images. Thus* 7 *sites* $\times$ 10 *seeds* $\times$ 2 *target directions* $=$ 140 *stimuli (*70 *per direction). Each image was optimized for a fixed* 2,048 *NAdam steps under the most heavily regularized decorrelated-spectrum regime (augmentation noise* 0.30*, spectral decay* $\alpha = 2.5$*) on a* $768 \times 768$*-px canvas.*

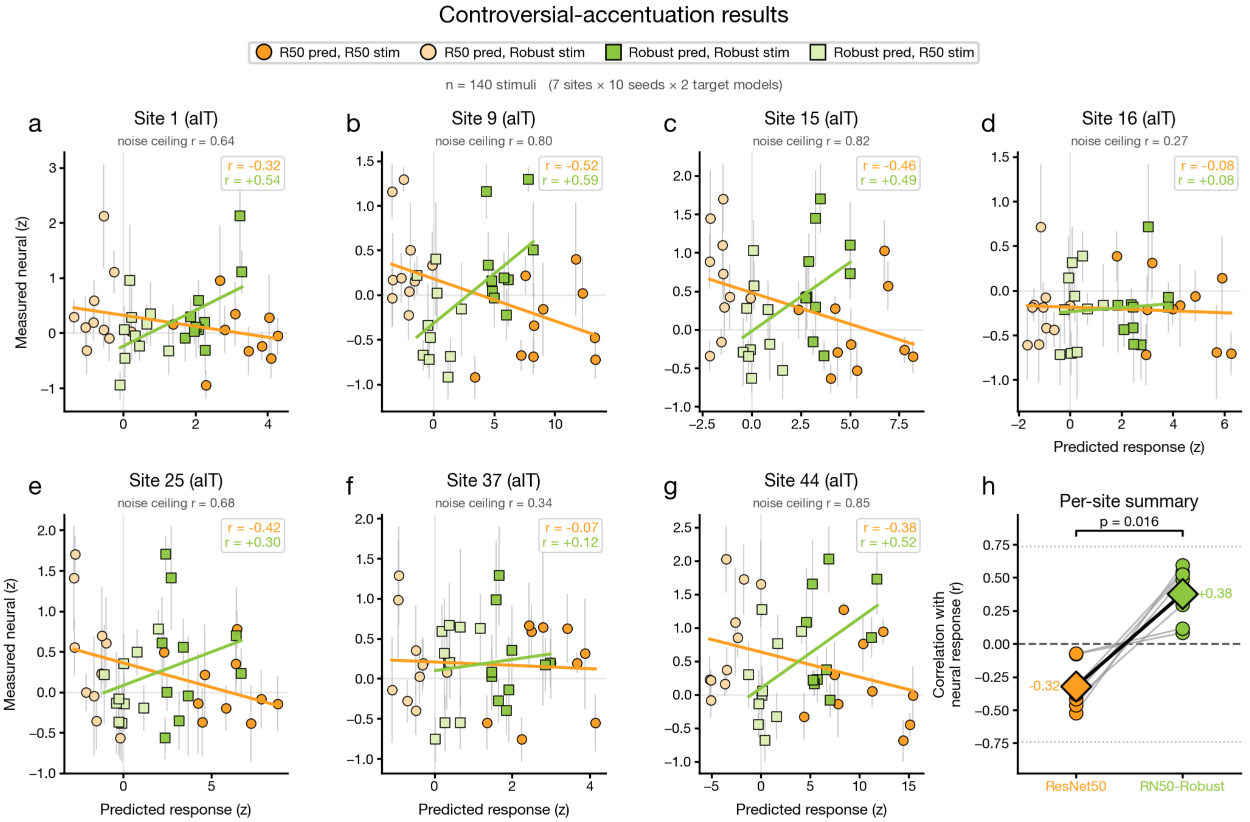


***Supplementary Figure 38. Controversial-accentuation results.*** *Site-by-site results of the controversial-accentuation experiment (Monkey R, aIT), in which each image was synthesized to drive one model's predicted response up while suppressing the other's, forcing the two models to make opposing predictions for the same stimuli. (A-G) Per-site scatter of measured neural response (*$z$*-scored,* $y$*) versus model prediction (*$x$*) for the seven targeted aIT sites (channels* 1, 9, 15, 16, 25, 37, 44*;* $n = 140$ *stimuli total,* 7 *sites* $\times$ 10 *seeds* $\times$ 2 *target models). Orange circles show standard ResNet50 predictions and green squares show ResNet50-Robust predictions (Lasso encoders); full color marks stimuli favoring the plotting model's family, faded color the opposing family. Lines represent least-squares fits per model, boxed values provide the per-site correlations per model, while the subtitle shows the maximum achievable correlation noise ceiling per site. Adversarially trained model predictions better track the neural response (positive* $r$*) while standard ResNet50 predictions are generally anticorrelated with it (negative* $r$*). (H) Per-site summaries: Pearson* $r$ *with the neural response for ResNet50 predictions versus ResNet50-Robust predictions, with diamonds and connecting line marking group means (ResNet50* $-0.32$*, RN50-Robust* $+0.38$*). Gray dotted lines mark the pooled achievable* $\pm$ *noise-ceiling envelope. Adversarially trained models exceed standard across sites (Wilcoxon signed-rank* $p = 0.016$*), indicating that standard ResNet50's brain alignment derives largely from features shared with its adversarially trained counterpart.*

### Adversarial sensitivity and input-gradient structure

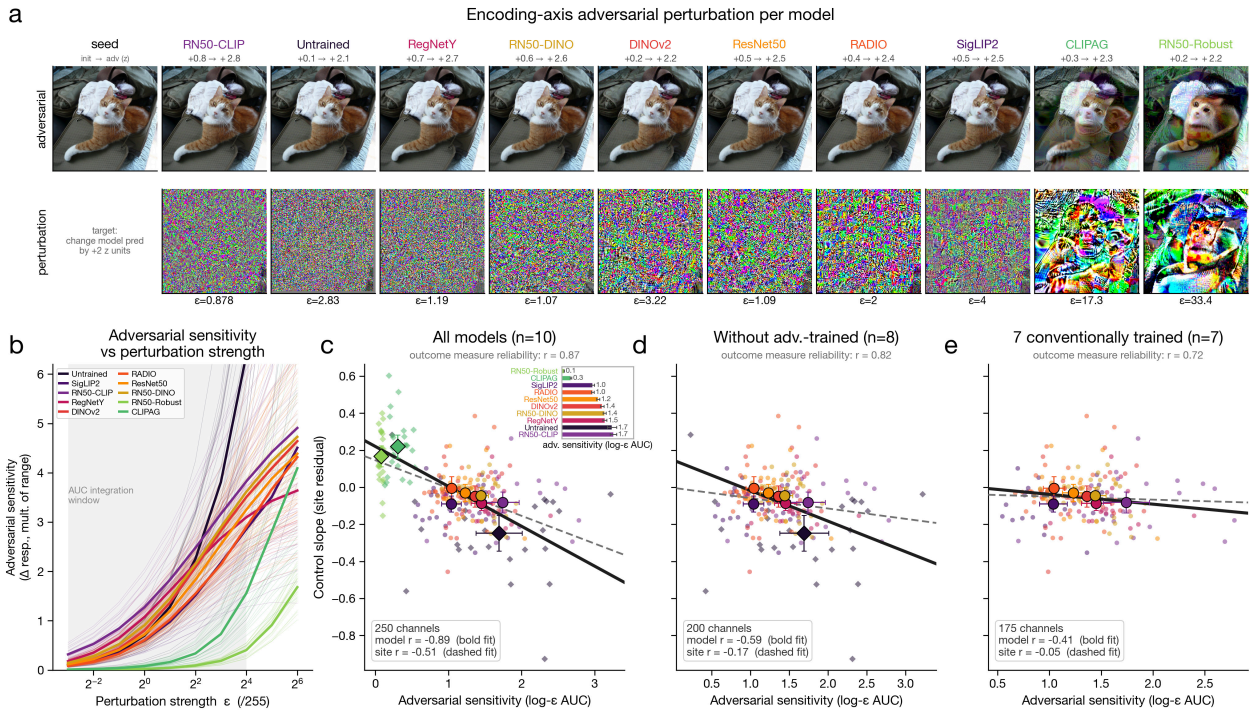


***Supplementary Figure 39. Encoding-axis adversarial sensitivity and neural control.*** *Although only two backbones were explicitly adversarially trained, we can measure adversarial sensitivity continuously for every one of the* 250 *fitted encoding axes. This is because each backbone plus its encoding readout is a differentiable input-output function. We implement attacks using pixel-space projected gradient descent (PGD). (A) Mosaic: for each model, the top row shows the minimally perturbed version of our cat seed image such that the model's prediction for the example site (Monkey P ch8) goes up by* $+2$ *z; the bottom row shows the perturbation itself, with the minimal* $\epsilon$ *annotated beneath. We implement a sweep of* $\epsilon$ *values such that we can identify the minimal* $\epsilon$ *that achieves almost exactly a* $+2$ *z update of the prediction. Columns are ordered by per-model sensitivity value. (B) We plot encoding adversarial sensitivity against perturbation strength* $\epsilon$*. Faint traces show the* 25 *trajectories per model and bolder traces show the model mean. The shaded area marks the* $\epsilon$ *window that is being integrated for the AUC metric. (C)–(E) Adversarial sensitivity (log-*$\epsilon$ *AUC on the* 100 *held-out NSD images) versus site-residualized control slope (residualized against the per-site mean over the* 9 *trained models) for three model subsets (all* 10 *models; without the adversarially trained models,* $n = 8$*; the* 7 *conventionally trained models, with the untrained and both adversarially trained models excluded); dashed line = site-level fit, bold line = model-mean fit; panel (C) carries the per-model sensitivity inset. Sensitivity tracks control over the full model set, but the relationship collapses once the adversarially trained models are removed.*

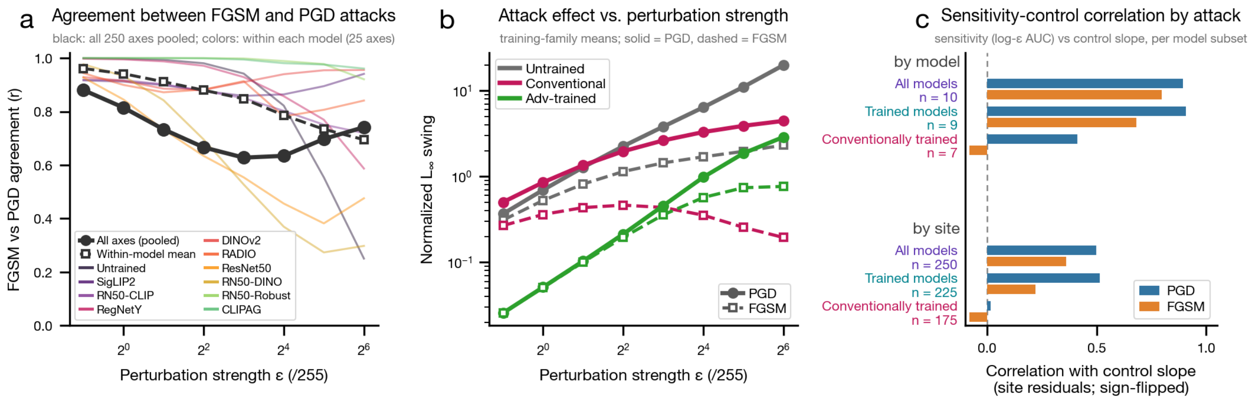


***Supplementary Figure 40. Adversarial attack-method comparisons.*** *Validation that the adversarial-sensitivity probe used in the main analysis is stable across different methods of implementing adversarial attacks. (A) We plot the agreement between a single-step FGSM attack and a multi-step PGD attack across perturbation budgets* $\epsilon$ *(log-2 axis,* $2^{-1}$ *to* $2^6$ *over* $255$*). Here agreement varies as a function of perturbation magnitude, reaching a minimum of approximately* $0.63$ *at intermediate budgets while remaining substantially above zero throughout. Thus, FGSM and PGD yield broadly consistent rankings of readout sensitivity across the full range of perturbation strengths. (B) Normalized* $L_\infty$ *swing as a function of* $\epsilon$ *for untrained (gray), conventionally trained (crimson), and adversarially trained (green) model groups, shown separately for PGD (solid circles) and FGSM (dashed open squares). Across all perturbation budgets and for both attack methods, adversarially trained models show the lowest sensitivity, occupying a distinct low-swing regime relative to the other model groups. (C) The relationship between adversarial sensitivity and control is reproduced using FGSM. Horizontal bars show the correlation between adversarial sensitivity (log-*$\epsilon$ *AUC over* $[0.125, 16]/255$*, the main-figure measure) and the control slope, computed from site residuals and oriented such that the relationship across the full model set is positive, for PGD (blue) and FGSM (orange). Correlations are shown at both the model level (top: all* $10$ *models,* $9$ *trained models, and* $7$ *conventionally trained models) and the site level (bottom:* $n = 250$*,* $225$*, and* $175$*). FGSM reproduces the PGD relationship across the full model set and the subset of nine trained models. However, for both attacks, this relationship collapses toward zero when considering only the seven conventionally trained models, with a small sign reversal for FGSM. Thus, the sensitivity-control relationship is specifically driven by the inclusion of the two adversarially trained models.*

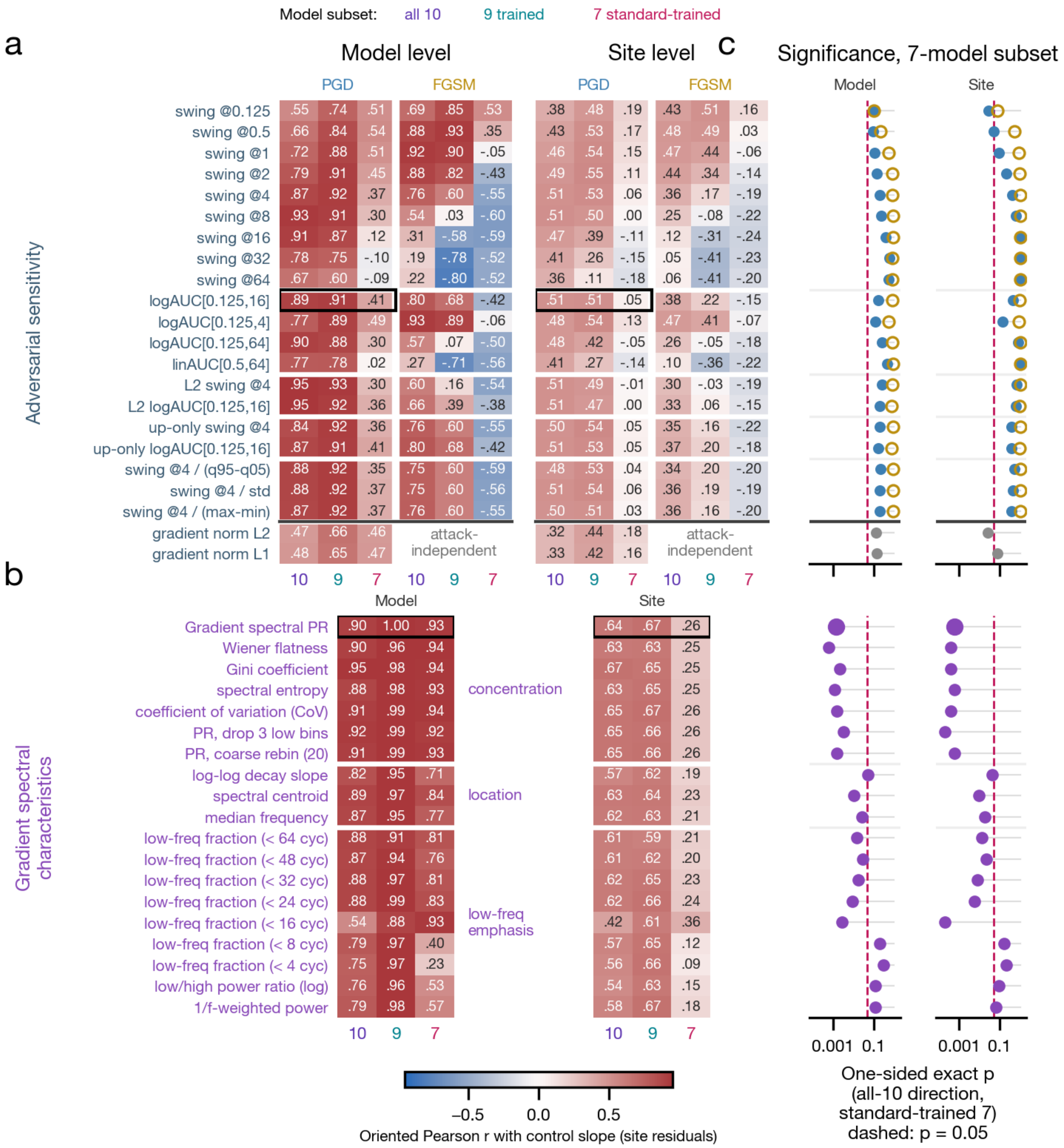

***Supplementary Figure 41. Alternative metrics for estimating adversarial sensitivity and gradient spectral concentration.*** *We find that while adversarial sensitivity correlates with control over the full model set, the relationship diminishes once the adversarially trained models are removed, and this trend replicates across essentially every tested alternative formulation of adversarial sensitivity. This includes multiple attack methods: PGD and FGSM. In contrast, the relationship between gradient-spectral summaries and control outcomes survives, regardless of which summary statistic is used. All analyses here use sensitivity measured on held-out NSD images (never involved in fitting the encoding models or as synthesis seeds); control slope values are site-residualized against the* 9 *trained models' per-site mean. Model-level significance across the* 7 *conventionally trained model subset is assessed here with an exact permutation test: all* 7! *permutations of the seven model-mean control values are examined relative to the corresponding predictor values (unshuffled), Pearson* $r$ *is recomputed for each permutation, and the observed correlation is evaluated against this null distribution. The plotted correlation values are sign-flipped so that each measure's association with*

*control across the full 10-model set is positive, and the correlations for the reduced model subsets use the same orientation, such that any negative values indicate a reversal of the relationship observed across the 10-model set. (A) Adversarial-sensitivity measures crossed with attack method (PGD and matched single-step FGSM) and model subset (all 10 / 9 trained / 7 conventionally trained): oriented Pearson r with control slope at the model level and the site level. The measures here span per-ϵ swing (0.125–64/255), log-AUC windows, linear AUC, the $L_2$ attack model, up-only attacks, and alternative response normalizers; the input-gradient norms are attack-independent. The boxed entry is the main-figure default (PGD log-AUC over $[0.125, 16]/255$). (B) Gradient spectral characteristics under the same model subsets (attack-independent, so no PGD/FGSM pairing), grouped by concentration, location, and low-frequency emphasis; the boxed entry is the main-figure gradient-spectral measure (the participation ratio is an exact monotone transform of the CoV: $\mathrm{PR} = N/(1 + \mathrm{CoV}^2)$). (C) Significance for the 7 conventionally trained model subset: one-sided exact-permutation p-values for each measure at the model level (left) and the site level (right), oriented in the all-10-model direction; the dashed line marks $p = 0.05$.*

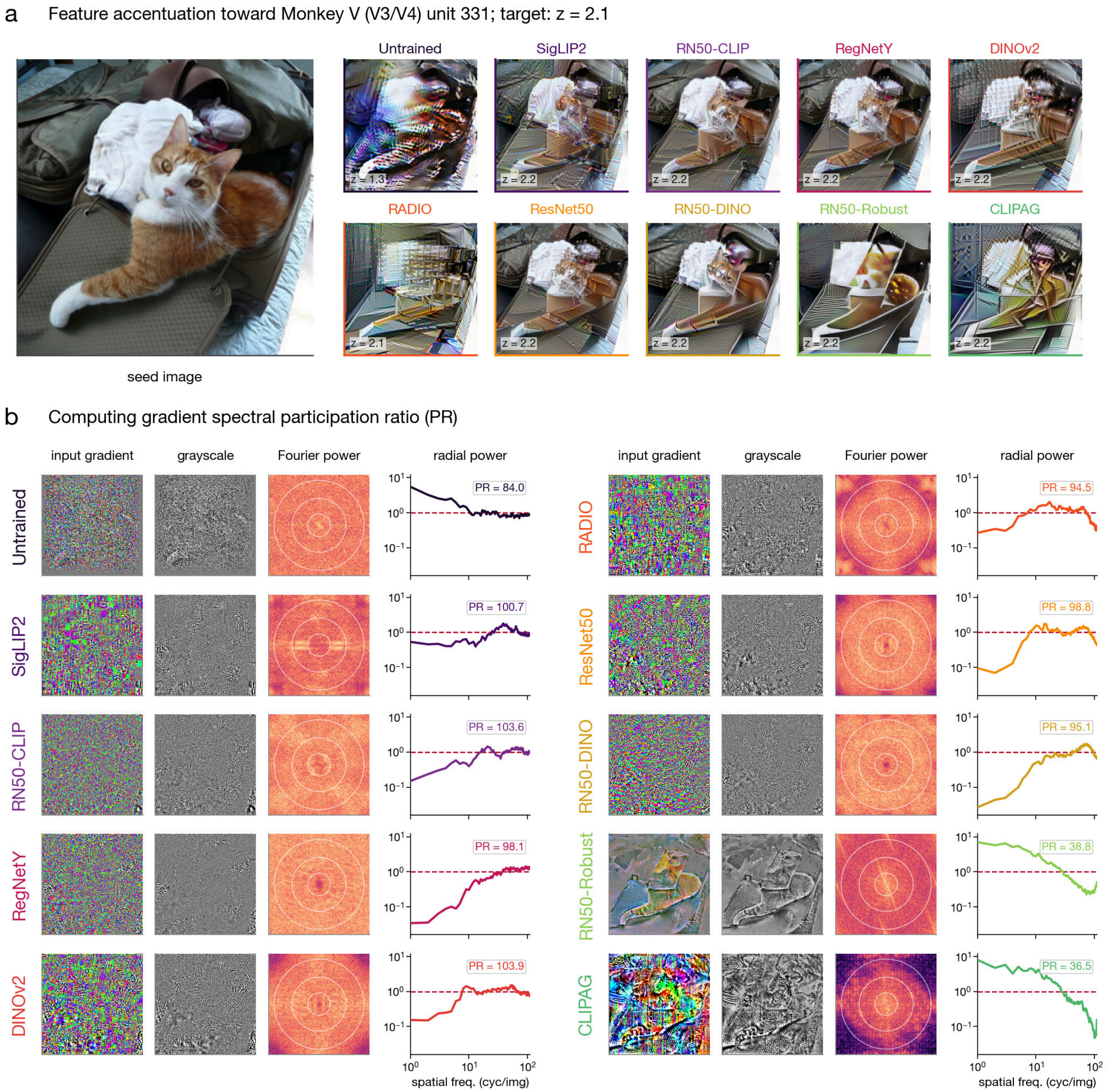


***Supplementary Figure 42. Computing gradient spectral participation ratio (PR).*** *Feature accentuations and the input-gradient spectrum pipeline for one natural seed image at a single recording site (Monkey V V3/V4, unit 331). (A)*

*The natural seed image $x$ (a cat) alongside a $2 \times 5$ mosaic of the ten models' feature accentuations of that seed toward the site, each driven to the highest response reachable by all trained models (target $z = 2.1$; the Untrained AlexNet is exempt and shown at its own maximum); the achieved $z$ is printed on each thumbnail. (B) The PR pipeline for the same seed and site, one row per model (two blocks of five, in Fig. 4 order) across four steps. First column: the input gradient of the site's encoding readout, $\partial(\mathbf{w} \cdot \phi(x))/\partial x$, from one backward pass of the fitted encoding prediction through the model to the image (three-channel; displayed as $0.5 + \mathrm{grad/s.d.}$). Second column: the gradient reduced to grayscale by averaging over the RGB channels (signed-square-root stretched for visibility; the input to the Fourier transform). Third column: its 2D Fourier power spectrum $|\mathcal{F}|^2$ (fftshift-centered, log scale; white rings mark radial frequency bins). Fourth column: radial averaging of the 2D power gives the 1D radial power profile (log-log; spatial frequency $1$–$111$ cyc/img), normalized so its arithmetic mean (red dashed) equals one, so all rows share one axis. The spectral summary used here is the participation ratio of the gradient frequency profile, $\mathrm{PR} = \big(\sum_f P_f\big)^2 / \sum_f P_f^2$. We use this to convey the effective number of frequency bins carrying power. PR tends to be higher for the conventionally trained models, whose gradient spectra are more broadband, and relatively lower for the adversarially trained models, whose power concentrates more at low frequencies. The per-site and per-model spectral PR reported in Fig. 5 are computed from gradients estimated over $100$ held-out natural images, while the seed-based pipeline shown here is an illustrative example. Critically, the computation of the gradient PR depends only on the encoding model. It is independent of the feature-accentuation synthesis procedure and the resulting stimuli.*

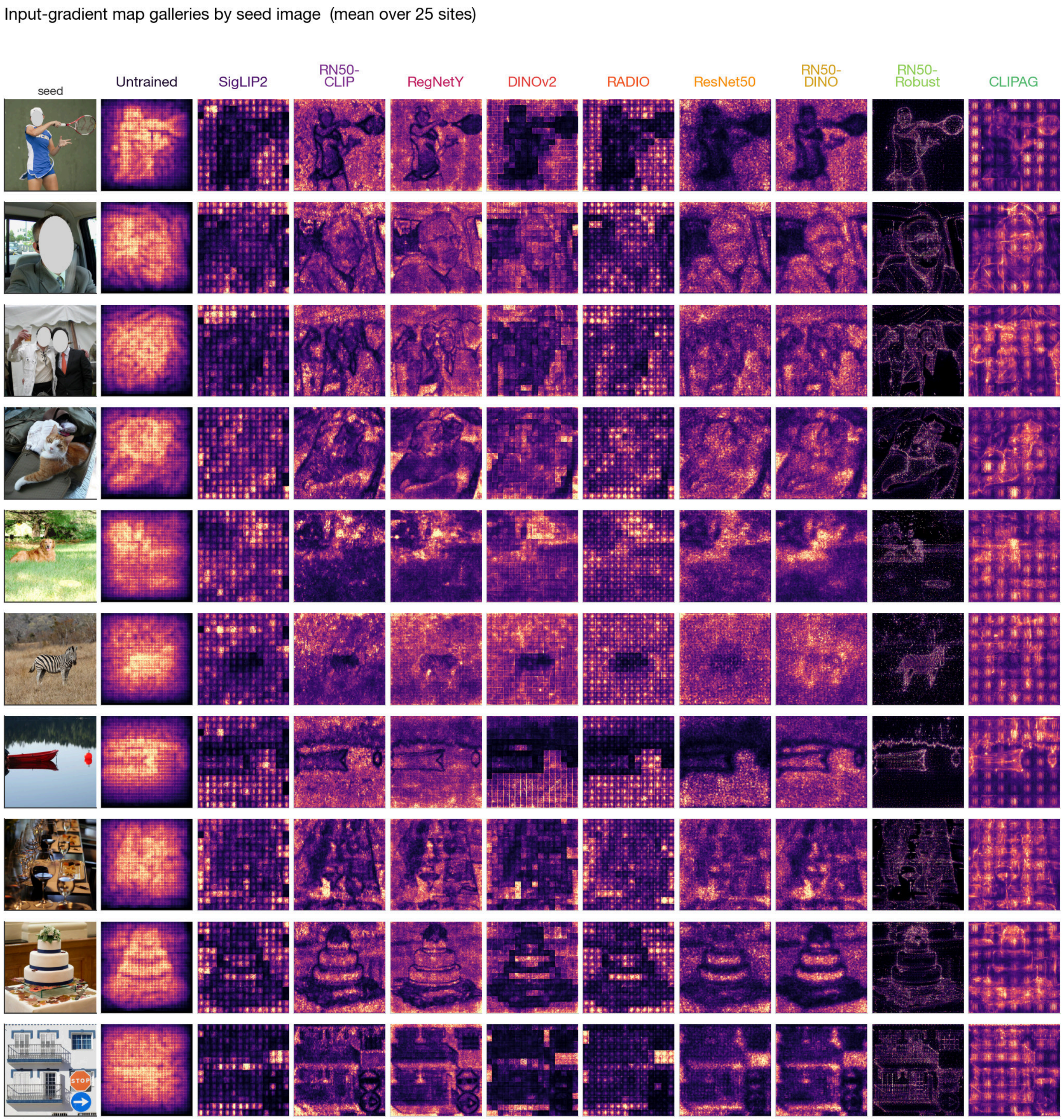


***Supplementary Figure 43. Input-gradient map galleries by seed image.*** *Site-averaged input-gradient saliency maps for each of the ten encoding models, arranged as ten seed rows by ten model columns, plus a left reference-seed column showing the natural seed images. Each cell in the ten model columns shows a given model's mean gradient map for that seed. These are computed as the $L_2$ magnitude of the model's input-output gradient across color channels, normalized per-map and then averaged over all 25 site-specific encodings for that model. These maps use a common magma color scale (shared $v_{\max}$ reflects the 99th percentile across all models).*

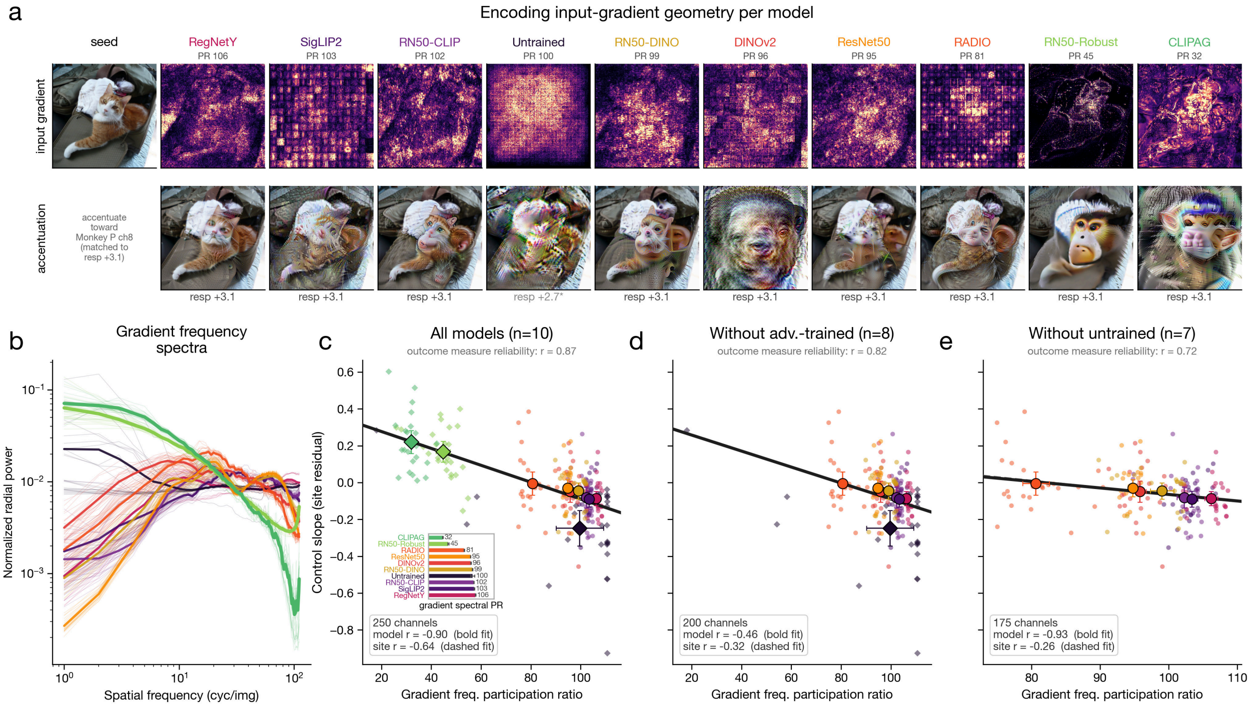


***Supplementary Figure 44. Encoding-gradient geometry and neural control.*** *Here we show a standalone version of the gradient-geometry content that is displayed in main Fig. 5, via the same layout. All spectral measures use the input-gradient power spectra computed over the* 100 *held-out NSD images (the same probe set as used in main Fig. 5). (A) Mosaic: the top row shows a seed image (cat) and each model's input-gradient saliency map (brighter = larger gradient magnitude), ordered left to right by gradient spectral participation ratio (PR value above each map), from the least concentrated models on the left (RegNetY PR* 106*, SigLIP2* 103*) to the two adversarially trained models on the right (RN50-Robust* 45*, CLIPAG* 32*); the bottom row shows the accentuation of that seed toward an example cIT site (Monkey P ch8), each matched to an achieved response of* $+3.1$ $z$ *(Untrained is capped at* $+2.7$*, asterisk). Adversarially trained models tend to have gradient sensitivity located around surfaces and objects, reflecting lower spatial frequencies, whereas the untrained and standard models tend to show more diffuse, high-frequency gradients, including visible transformer patchification. (B) Per-model gradient frequency spectra: we plot normalized radial power versus spatial frequency (cyc/img, log-log); thin lines show per-site data, bold lines show per-model mean data. (C)–(E) Site-residualized control slope versus gradient spectral PR for three different model subsets, with larger markers showing per-model means (*$\pm 95\%$ *CI; diamonds are used for the untrained and the two adversarially trained models, circles are used for others). A dashed site-level linear fit and a bold model-level linear fit are plotted; lower PR (greater low-frequency concentration in the spatial frequency spectra) is associated with stronger control outcomes. (C) All* 10 *models (*250 *model-site axes): model* $r = -0.90$*, site* $r = -0.64$ *(outcome-measure reliability* $r = 0.87$*); the inset bars give per-model mean PR, from CLIPAG* 32 *(most concentrated) to RegNetY* 106 *(least). (D) With the two adversarially trained models removed (*$n = 8$*;* 200 *model-site axes): model* $r = -0.46$*, site* $r = -0.32$ *(reliability* $r = 0.82$*). (E) The* 7 *conventionally trained models (untrained and both adversarially trained models excluded;* 175 *model-site axes): model* $r = -0.93$*, site* $r = -0.26$ *(reliability* $r = 0.72$*).*

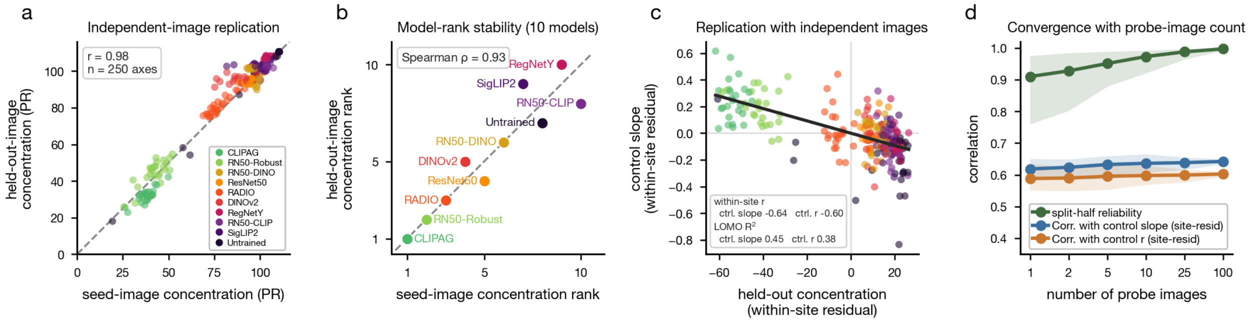


***Supplementary Figure 45. Robustness of gradient spectral participation ratio to probe images and analysis choices.*** *Gradient spectral participation ratio (PR) is a property of each fitted encoding axis; here we assess whether PR is stable across different image sets used to compute it, i.e., the* $100$ *held-out NSD images versus the* $10$ *natural seed images used for feature accentuation. (A) Independent-image replication: we measure per-axis PR on the* $10$ *synthesis seeds vs. on the* $100$ *held-out images, and the two sets achieve nearly perfect agreement (*$r = 0.98$*,* $n = 250$ *axes; points colored by model). (B) Stability of model rankings: the* $10$*-model ordering of mean PR is conserved whether PR is estimated using the* $10$ *seeds or the* $100$ *held-out images (Spearman* $\rho = 0.93$*); the two adversarially trained models, CLIPAG and RN50-Robust, remain the two most concentrated (lowest PR). (C) Held-out-image PR predicts neural control on independent images: within-site (residualized, with mean performance subtracted across the models per site) PR vs. control slope (points colored by model, black regression line); within-site* $r = -0.64$ *(control slope outcome measure) and* $-0.60$ *(control* $r$ *outcome measure), and leave-one-model-out cross-validated* $R^2 = 0.45$ *(control slope) and* $0.38$ *(control* $r$*). (D) Varying the number of images used to assess gradient spectral PR: we compute split-half reliability of the per-axis PR estimate (green) and its site-residualized association with control slope (blue) and control* $r$ *(orange), as a function of the number of probe images (*$1$*–*$100$*; bootstrap subsampling of the* $100$ *held-out images, mean and* $95\%$ *CI shaded). All estimates stabilize within a small number of images.*

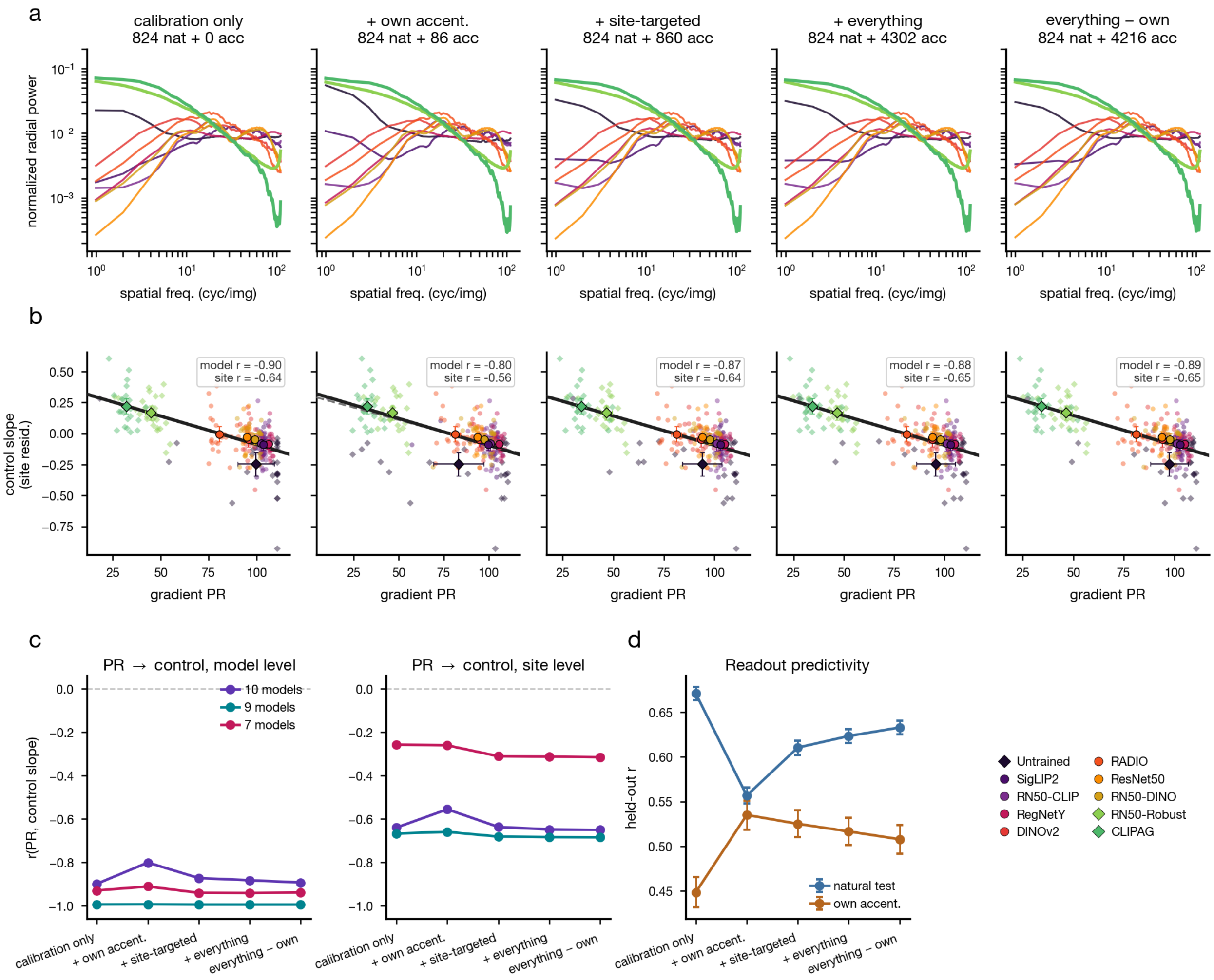


***Supplementary Figure 46. Gradient spectra and their prediction of control across encoding training diets.*** *The main-figure gradient-spectra analyses redone across a ladder of encoding-readout training diets, asking whether fitting the readout on more (or cleaner) data changes the gradient spectra, or how well their spectral participation ratio (PR) predicts neural control. Diets per site-model pair (mean fitting-set sizes over the 250 pairs; within each pair every diet uses the same natural calibration set, mean* $n = 824.2$ *images, and differs only in the accentuated images added): (d0) calibration images only (the synthesis-time readouts, as in the main figure;* $+0$ *accentuated); (d1) plus the pair's own accentuations (*$+86$*); (d2) plus all site-targeted accentuations from every model (*$+860$*); (d3) plus all accentuated stimuli (the pooled refit;* $+4{,}302$*); (d4) everything minus the pair's own control-scored images (zero circularity;* $+4{,}216$*). (A) Per-model mean held-out gradient spectra per diet. (B) Gradient spectral PR versus site-residualized control slope per diet, exact main-figure recipe (site dots plus model means with* $95\%$ *CI, dashed site-level fit, bold model-mean fit; held-out* $100$*-NSD-image profiles;* $9$*-trained residual reference). (C) Summary of model- and site-level correlations across diets for the* $10/9/7$ *model subsets. (D) Readout predictivity along the ladder (natural-image validation* $r$ *plus own-accentuation* $r$*; out-of-fold when the diet contains those images). The gradient spectra, and their prediction of neural control, are essentially unchanged across training diets.*

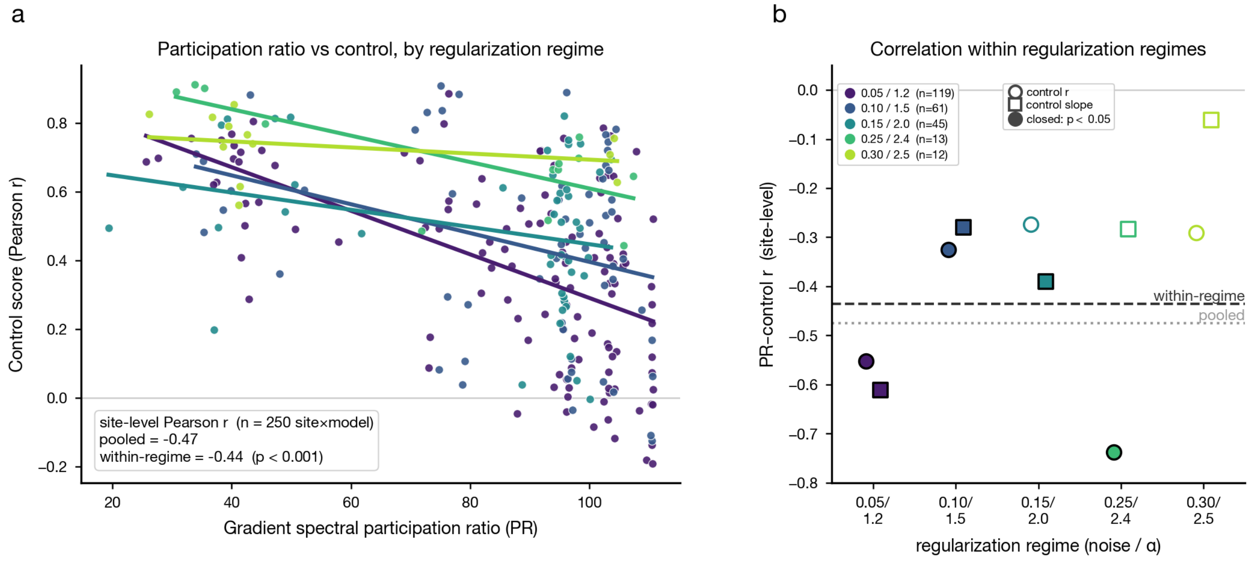


***Supplementary Figure 47. Gradient spectral participation ratio and control within matched synthesis-regularization regimes.*** *Because accentuation stimuli were synthesized under a Fourier-domain regularizer (an augmentation-noise / spectral-decay* $\alpha$ *rung selected automatically per site-model), gradient spectral concentration might in principle predict control only because low-frequency-concentrated gradients suit that optimizer rather than because they reflect a brain-like property. To test this, the* $n = 250$ *site-models were stratified by their selected regularization regime (five discrete rungs;* $n = 119, 61, 45, 13, 12$*) and the PR–control relationship was re-evaluated within regimes. (A) Gradient spectral participation ratio (PR, the effective number of frequency bins of the input-gradient radial power spectrum over* $1$–$111$ *cyc/img) versus control score (Pearson* $r$ *of predicted vs. measured responses over the accentuated set), each site-model colored by its regularization regime (augmentation noise /* $\alpha$*) with a per-regime ordinary-least-squares fit; the pooled and within-regime (regime-mean-centered) correlations are inset. (B) Per-regime Pearson correlation between PR and control score (circles) and control slope (squares), colored by regime; the dashed line marks the pooled within-regime correlation and the dotted line the raw pooled correlation. Matching on regularization attenuates the PR–control correlation only from* $r = -0.47$ *to* $r = -0.44$ *(control score;* $p < 0.001$*), and the correlation is negative (lower PR, stronger control) in all five regimes, indicating that the relationship reflects the models' intrinsic gradient structure rather than an artifact of the synthesis optimizer.*

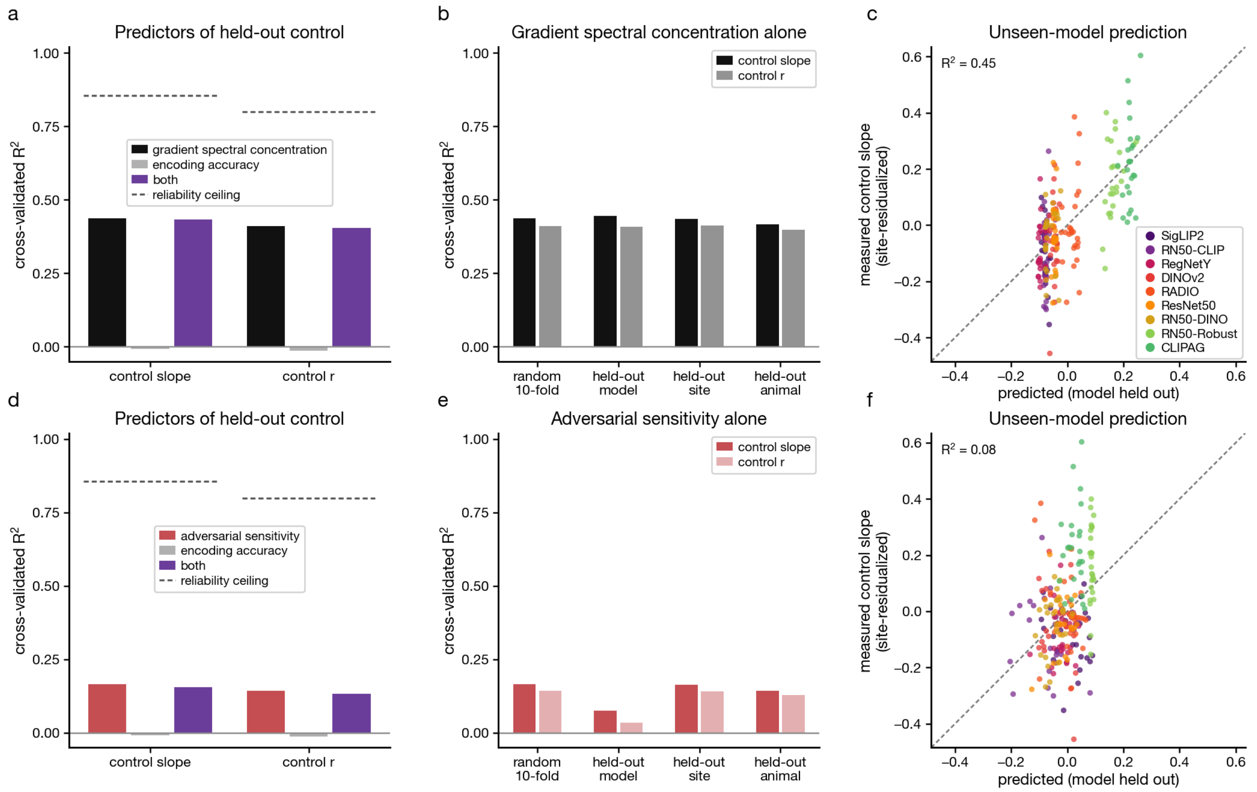


***Supplementary Figure 48. Cross-validated prediction of parametric control among the trained models.*** *What predicts held-out parametric control among the* 9 *trained models (*225 *encoding axes): gradient spectral concentration (top row) versus adversarial sensitivity (bottom row), each against natural-image encoding accuracy. The untrained model is excluded as an out-of-distribution extreme that inflates every correlation and breaks leave-one-model-out evaluation on the site-residualized outcome. Outcomes are control slope and control $r$, each site-residualized against the per-site mean over the* 9 *trained models. Predictors are the gradient spectral participation ratio on the* 100 *held-out natural images, first-order adversarial sensitivity (both main-figure quantities), and the calibration-phase encoding validation $r$; models are ordinary least squares with a single global slope per predictor (the only design under which an unseen animal can be predicted at all), z-scored within each training fold. (A, D)* 10*-fold cross-validated $R^2$ for the row's predictor alone, encoding accuracy alone, and both together, against the seed split-half explainable ceiling (dashed). (B, E) The row's predictor alone under progressively harder held-out regimes: random folds, whole held-out models, whole held-out sites, and whole held-out animals. (C, F) Measured versus predicted site-residualized control slope with the model itself held out (leave-one-model-out). Spectral PR generalizes across every held-out regime, whereas adversarial sensitivity and encoding accuracy do not.*

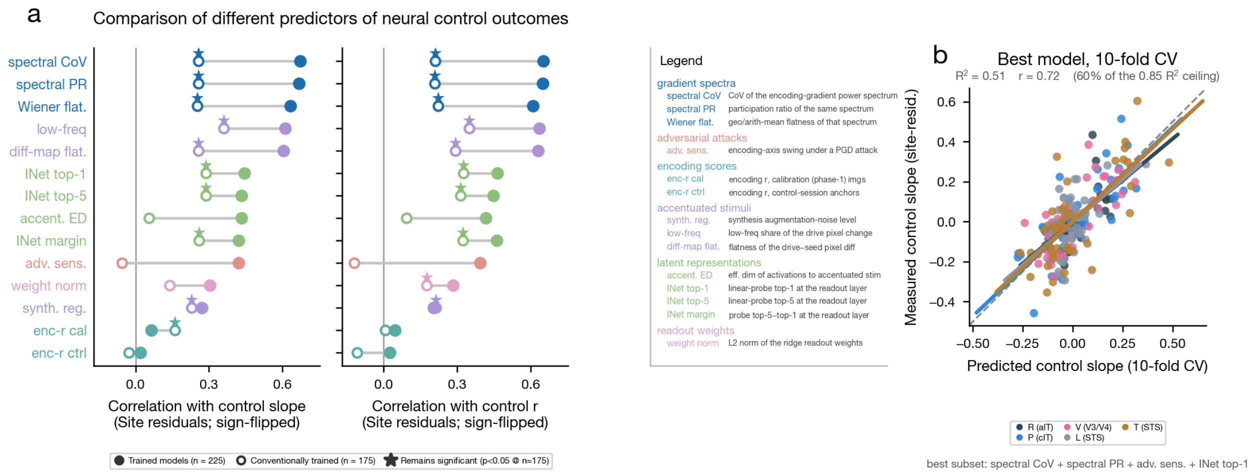


***Supplementary Figure 49. Predicting neural control: single-predictor leaderboard and cross-validated best model.*** *Site-residualized, cross-validated comparison of candidate predictors of parametric control, contrasting the trained models ($n = 225$ site-models) with the conventionally-trained subset ($n = 175$). (A) Single-predictor site-level correlation with control slope (left) and control $r$ (right; sign-flipped site residuals); each candidate predictor is a dumbbell with the trained-models value filled and the conventionally-trained value open, and a star marking predictors that remain significant ($p < 0.05$) at $n = 175$. Predictors are grouped by what they measure (legend): gradient spectra (spectral CoV, spectral PR, Wiener flatness), adversarial attacks (encoding-axis swing under a PGD attack), encoding scores (calibration- and control-phase encoding $r$, both over held-out images), accentuated stimuli (synthesis regularization, low-frequency share of the drive pixel change, flatness of the drive–seed pixel difference), latent representations (effective dimensionality of activations to accentuated stimuli; ImageNet linear-probe top-1, top-5, and margin at the readout layer), and readout weights (L2 norm of the ridge readout weights). The gradient-spectral summaries lead and retain significance among the conventionally trained models, whereas adversarial sensitivity collapses there (open marker crosses zero). (B) The best predictor subset (spectral CoV + spectral PR + adversarial sensitivity + ImageNet top-1), identified by exhaustively scoring every subset of the eleven site-varying predictor groups by repeated 10-fold cross-validated $R^2$ under a per-animal design (per-animal intercepts and slopes), shown as predicted versus measured site-residualized control slope, one color per monkey with per-monkey fits and the identity line; $R^2 = 0.51$, $r = 0.72$, reaching $60\%$ of the $0.85$ split-half $R^2$ ceiling. We note that the same cross-validation folds were used to score and select predictors; best-model $R^2$ as reported here therefore reflects an optimistic upper bound on the outcome measure.*
